# Volumetric alteration of olfactory bulb and immune-related molecular changes in olfactory epithelium in first episode psychosis patients

**DOI:** 10.1101/2021.05.03.442464

**Authors:** Kun Yang, Jun Hua, Semra Etyemez, Adrian Paez, Neal Prasad, Koko Ishizuka, Akira Sawa, Vidyulata Kamath

## Abstract

**Background:** Olfactory dysfunction has been reproducibly reported in patients with psychosis, including first episode psychosis (FEP) patients. Consistently, structural abnormalities in the olfactory bulb (OB), a key region of the peripheral olfactory system, have also been reported in psychotic disorders. Meanwhile, air pollution and viral infections in the upper respiratory tract, including those of SARS-CoV-2, are reportedly risk factors for brain dysfunction and mental disorders. These risk factors can disturb the olfactory epithelium (OE) that is located adjacent to the OB and connected via direct neuronal projections. Nevertheless, it is unknown how a disturbance of the OE possibly affects the OB in the pathophysiological context of psychotic disorders.

**Methods:** We examined the volume of the OB in FEP patients and healthy controls from 3 Tesla magnetic resonance imaging and molecular expression profiles of olfactory neuronal cells (ONCs) enriched from freshly biopsied OE.

**Results:** We observed a reduction of the OB volume in FEP patients compared with healthy controls. We also observed a significant alteration in gene expression profiles in the ONCs from FEP patients, supporting the pathological changes in the OE. Among such molecular changes, immune-related molecules and pathways were underscored in association with the OB volume changes in FEP patients.

**Conclusions:** Our data support the OB and OE pathologies in FEP patients. Immune-related molecular changes in the OE can biologically link adverse factors in the nasal cavity, such as air pollution and viral infection, with the OB structural change, both of which have been reported for psychotic disorders.

## 1. Introduction

Deficits in olfactory function have been reproducibly reported in patients with psychotic disorders (Moberg et al., 2014). In these studies, three psychophysical measures (odor identification, odor discrimination, and odor detection threshold) have been frequently used (Cohen et al., 2012; Ishizuka et al., 2010; Kamath et al., 2018; Kiparizoska and Ikuta, 2017; Kopala et al., 1995; Malaspina et al., 1994; Moberg et al., 2014; Turetsky et al., 2009). Importantly, these deficits are observed in youths at-risk for psychosis (Brewer et al., 2003; Kotlicka-Antczak et al., 2017; Woodberry et al., 2010). Accordingly, olfactory dysfunction is also one of the key deficits in patients with first episode psychosis (FEP) (Chen et al., 2018b; Kamath et al., 2014, 2018, 2019). Furthermore, olfactory deficits appear to predict poor outcomes in patients with schizophrenia (SZ) and can identify patients at high risk of developing unremitting negative symptoms, particularly anhedonia (Good et al., 2010; Lin et al., 2015).

Olfactory neurons in the olfactory epithelium (OE) that are exposed to the nasal cavity receive odorants and project to glomeruli in the olfactory bulb (OB) where they synapse on mitral and tufted cells (Mori and Sakano, 2011). Mitral and tufted cells then project to the primary olfactory cortex, where the information from olfactory neurons is transmitted to the higher cortex, such as the orbitofrontal cortex (OFC) and medial prefrontal cortex (mPFC), as well as other central brain regions that are involved in emotion and cognition (Mori and Sakano, 2021). It is important to note that the external information perceived in the olfactory system can be transmitted with fewer synapses (only two, by bypassing the thalamus) to the central brain regions for emotion and cognition compared with other perceptions (Su et al., 2009).

Deficits in the peripheral olfactory system (e.g., the OE and OB), or the central olfactory system, or both can elicit the aforementioned olfactory deficits (Sawa and Cascella, 2009). Meanwhile, volumetric changes of the OB, mainly by magnetic resonance imaging (MRI) assessment, have been reported in multiple brain disorders (Chen et al., 2014, 2018a; Li et al., 2016; Petekkaya et al., 2020). A reduction in the OB volume has also been reported in psychiatric disorders, such as major depressive disorders and SZ (Asal et al., 2018; Gul et al., 2015; Negoias et al., 2010; Nguyen et al., 2011; Rottstädt et al., 2018; Rottstaedt et al., 2018a, 2018b; Turetsky et al., 2000). Importantly, there are reports that indicate a volume reduction in subjects at high risk for psychosis (Turetsky et al., 2003, 2018), suggesting that the changes found in SZ and related disorders may not be due to secondary effects of medication. More recently, SARS-CoV-2 infected patients with olfactory dysfunction have reportedly shown smaller OB volumes compared with healthy subjects (Kandemirli et al., 2021), and a case report also indicated shrinkage of the OB in the course of SARS-CoV-2 infection (Chiu et al., 2021).

Compared with many reports about the OB in brain disorders, much less literature is available on the biological assessment of the OE in psychotic disorders. Some studies were performed with postmortem OE, which includes a seminal work by Arnold reporting a significant change in cell fate from stem cells to mature neurons (Arnold et al., 2001). Some of these results, but not all, were reproduced by another group (Minovi et al., 2015), which may be due to the heterogeneity of SZ pathophysiology (Owen et al., 2016). An outstanding question is whether the pathological features observed in postmortem OE from chronic SZ patients may represent the signatures of a long-term compensation to the key pathological events when psychosis occurs and is active. Accordingly, it is important to conduct biological assessments of the OE freshly biopsied from patients with early-stage psychosis.

Air pollution has been highlighted as a major risk factor for SZ and related psychotic disorders (Gao et al., 2017; Newbury et al., 2019; Pedersen et al., 2004), and can elicit pathological and immune/inflammatory responses in the OE at the biological levels (Ajmani et al., 2016). Viral infection can also be a major adverse event for the OE that is exposed to the nasal cavity. Studies on herpes simplex virus and Alzheimer’s Disease have suggested a link between viral infection in the nasal cavity and brain disorders (Eimer et al., 2018; Itzhaki et al., 2016, 2020; Readhead et al., 2018). More recently, intranasal infection with SARS-CoV-2 is underscored in the context of olfactory dysfunction, possibly followed by deficits in higher brain function and the development of psychosis (Kabbani and Olds, 2020; Zubair et al., 2020). These data suggest that peripheral olfactory system disturbances may underlie, as an important risk factor or driver, multiple neuropsychiatric disorders, including SZ and related psychotic disorders (Taquet et al., 2021; Varatharaj et al., 2020). Nevertheless, the causal impact and underlying mechanism of the peripheral olfactory pathology on brain disorders remain to be elucidated.

In the present study, we first aimed to build a working hypothesis that can link the following three observations in SZ, or at least in a subset of SZ: (1) adverse events in the nasal cavity are a major risk factor for disease; (2) possible changes in neural cell fate in the OE; and (3) volume reduction of the OB. Meanwhile, a recent basic science publication indicated that the stress response to induced inflammation in the OE could alter the cell fate of basal stem cells [horizontal basal stem cells (HBCs) in a mouse model] (Chen et al., 2019). In this animal model, subsequent OB shrinkage has also been reported (Chen et al., 2019). Inspired by these reports in basic science, we hypothesized that at least a subset of SZ patients might show molecular changes relevant to a stress response in the OE, which may also be associated with a volumetric alteration of the OB. To address this specific question, we conducted an unbiased molecular expression study on biopsied neuronal cells from the OE together with a volumetric assessment of the OB by using 3 Tesla magnetic resonance imaging (MRI) data from the same set of FEP patients and healthy controls. The biopsied neuronal cells (olfactory neuronal cells, ONCs) were used as molecular surrogates of neuronal cells, in particular those for stem cells and immature neuronal cells (Evgrafov et al., 2020; Horiuchi et al., 2013; Kano et al., 2013; Lavoie et al., 2017; Takayanagi et al., 2020).

## 2. Methods

This study was conducted in accordance with The Code of Ethics of the World Medical Association (1964 Declaration of Helsinki) and was approved by the Johns Hopkins School of Medicine Institutional Review Board. Written informed consent was obtained for all participants.

### 2.1. Study Participants

Adolescents and young adults between 18 and 35 years old were recruited from within and outside of the Johns Hopkins Hospital. Exclusion criteria included history of traumatic brain injury, neurologic condition, intellectual disability, cancer, viral infection, active substance abuse, and conditions affecting olfaction (e.g, history of sinus surgery, rhinoplasty, chronic rhinosinusitis). Controls were screened and excluded for a family history of schizophrenia or schizophrenia-spectrum disorder. Patients must be within 24 months of the onset of psychotic manifestations as assessed by study team psychiatrists using the Structured Clinical Interview for DSM-IV (SCID) and information from available medical records.

There were originally 24 FEP patients and 24 HC subjects with both OB volumes and RNA-Seq data from ONCs available for the present study. Subjects that self-reported smoking tobacco and cannabis were excluded from the study. In addition, one subject with an unusually large OB (z-score > 3) was considered as an outlier and excluded from the study. Together, the present study finally investigated the data from 22 HC subjects and 16 FEP patients. Patients were diagnosed with schizophrenia (*n* = 9), schizoaffective disorder (*n* = 2), bipolar disorder with psychosis (*n* = 3), major depressive disorder with psychosis (*n* = 1), and substance-induced psychosis (*n* = 1). The majority of patients, except one patient, were medicated. Antipsychotic medication dosages were converted to chlorpromazine (CPZ) equivalents using published reference tables using the Defined Daily Doses (DDDs) method (Leucht et al., 2016).

### 2.2. 3T MRI scans and OB volume

All images are acquired on a 3T Philips MRI scanner (Philips Healthcare, Best, The Netherlands). The three-dimensional, T1-weighted, magnetization-prepared rapid acquisition with gradient echo (MPRAGE) scan was acquired for each participant with the following parameters: 1mm isotropic voxel, repetition time (TR) = 8.1ms, echo time (TE) = 3.7ms, flip angle = 8°, field of view (FOV) = 256mm x 256mm x 170mm, 170 slices, duration = 4minutes 16seconds.

The FMRIB Software Library (FSL, https://fsl.fmrib.ox.ac.uk/fsl/) software was used to normalize the T1-weighted MPRAGE images into the MNI space. Subsequent analysis was performed in Insight ToolKitSNAP (ITK-SNAP) (www.itksnap.org). The OB was manually identified and segmented on coronal images, with additional corrections made in the sagittal and axial planes. Volumetric measures were obtained using the built-in functions in ITK-SNAP at the level of the anterior cribriform plate following the procedures established previously (Turetsky et al., 2000). Two experienced researchers (A.G.P and N.P.) performed the segmentation separately and were blinded to participant information. After all segmentation was completed, discrepancies between the two researchers were assessed and final measurements were agreed upon. The intraclass correlation coefficient for inter-rater reliability between raters (AGP, NP) was 0.91.

### 2.3. ONCs obtained via nasal biopsy

The nasal biopsy was conducted at the Johns Hopkins Otolaryngology Clinic. The biopsy procedure was described previously (Kano et al., 2013; Narayan et al., 2014). ONCs were enriched from nasal biopsied tissue as follows: first, the olfactory neuroepithelial tissue was incubated with 2.4 U/mL Dispase II for 45 min at 37°C, and mechanically minced into small pieces. Then, the tissue pieces were further treated with 0.25 mg/mL collagenase A for 10 min at 37°C. Dissociated cells were gently suspended, and centrifuged to obtain pellets. Cell pellets were resuspended in DMEM/F12 supplemented with 10% FBS and antibiotics (D-MEM/F12 medium), and tissue debris was removed by transferring only the cell suspension into a new tube. Cells were then plated on a 6-well plate (plate A) in fresh D-MEM/F12 medium for 2 days. We next transferred cells floating or loosely attached to plate A into a new 6 well plate (plate B), and further incubated these cells for another 5 days. Then, cells floating or loosely attached to plate B were transferred into another new 6 well plate (plate C). Cells in plate C were further incubated to be confluent, collected as ONCs, and stored in liquid nitrogen. When it was time to conduct downstream experiments, ONCs were recovered from the liquid nitrogen and cultured in D-MEM/F12 medium with media changes every 2-3 days. In summary, without any genetic and chemical reprogramming and conversion, we enriched the neuronal cell population (Horiuchi et al., 2013; Mor et al., 2013).

### 2.4. Molecular expression profiles (RNA-sequencing) of ONCs

Total RNA was isolated from ONCs using the RNeasy Plus Mini Kit (Qiagen). RNA quality was assessed on the Agilent Fragment Analyzer using an RNA High Sensitivity (DNF-472) and quantified using a Qubit 4 RNA BR kit (Thermo Fisher). RNA libraries were prepared with 500ng total RNA. Library generation was accomplished using the NEBNext Ultra II Directional RNA Library Prep Kit for Illumina (E7760 and E7490) following the NEBNext Poly(A) mRNA Magnetic Isolation Module protocol. Libraries were enriched using 11 cycles of PCR amplification. Library quality and quantification was assessed on the Agilent Fragment Analyzer using a High Sensitivity NGS Kit (DNF-474) and a Qubit 4 RNA BR kit (Thermo Fisher). Samples were then normalized to 4nM and pooled in equimolar amounts. Paired-End Sequencing was performed using Illumina’s NovaSeq6000 S4 200 cycle kit.

FastQC (Andrews, 2010) was used to check the quality of reads. High-quality data were obtained from raw data by using cutadapt (Martin, 2011) to remove adapters, primers, and reads with low quality (option -q 10) or shorter than 20 nt. Hisat2 (option --dta) (Pertea et al., 2016) was used to map the clean reads to the human genome, version GRCh38 (Genome Reference Consortium Human Build 38). Stringtie (Pertea et al., 2016) was used to assemble and merge transcripts and estimate transcript abundance. A Python script (prepDE.py) provided by the Stringtie developer (Pertea et al., 2016) was used to create count tables for further analysis.

### 2.5. Two-group comparison analysis

General linear regression analysis was performed to compare the volumes of the total (referred to as OB), left (referred to as OB_L), and right (referred to as OB_R) OB between FEP patients and HC subjects. Considering that subjects with larger heads may have larger OBs, the ratio between OB, OB_L, and OB_R volumes and intracranial volume was used in the analysis. In the general linear regression, the dependent variables were the volume and the diagnosis group (FEP or HC), and the covariates included age, gender, and race.

Differential expression analysis was performed to compare the expressional profiles between FEP patients and HC subjects using R library *DESeq2* (Love et al., 2014). Age, gender, race and the 3 hidden/unknown confounding factors identified by *sva* (Leek et al., 2012) were included as covariates in the design formula. The Benjamini and Hochberg (BH) procedure was used for multiple comparison correction.

### 2.6. Correlation analysis

General linear regression analysis was performed to evaluate the correlation between expressional profiles and the OB_R volume. Genes with low expression levels (TPM (Transcripts Per Million) values less than 0.1 in more than 20% of the total samples) were excluded from the analysis. Log transformed TPM values and the ratio between the OB_R volume and intracranial volume were used as the independent and dependent variables in the general linear regression analysis, respectively. For the HC group, age, gender, and race were included as covariates, while for the FEP group, age, gender, race, duration of illness (DOI), and CPZ dose were included as covariates. The BH procedure was used for multiple comparison correction. In addition, permutation testing (1000-fold) was conducted to assess the significance of the results from correlation analyses performed with adjustment for confounding variables.

### 2.7. Pathway and GWAS enrichment analysis

Gene set enrichment analysis (GSEA) (Subramanian et al., 2005) on the Reactome Pathway Database (Fabregat et al., 2018) was conducted to identify molecular pathways altered in FEP patients compared to HC subjects and pathways associated with the OB_R volume. GSEA is a threshold-free method that analyzes all genes based on their statistical significance. It searches for pathways whose genes are enriched at the top or bottom of the ranked gene list. In the analysis to identify pathways altered in FEP patients compared to HC subjects, all genes (including genes differentially and not differentially expressed between the FEP and HC groups) were ranked by the Wald statistic from *DESeq2* (Love et al., 2014), where the top and bottom of the ranked gene list are genes differentially increased and decreased in FEP patients compared to HC subjects, respectively. In the analysis to identify pathways associated with the OB_R volume, all genes (including genes correlated and not correlated with OB_R volumes) were ranked by t values from the correlation analysis between the OB_R volume and gene expression levels, where the top and bottom of the ranked gene list were genes positively and negatively associated with the OB_R volume, respectively. In the analyses for identifying pathways altered in FEP patients compared to HC subjects and for identifying pathways associated with the OB_R volume, the GSEAPreranked module was used and pathways with less than 10 genes or more than 500 genes were excluded (options -set_max 500 -set_min 10). The BH procedure was used for multiple comparison correction.

A GWAS enrichment analysis was performed to check if risk genes for immune-related disorders or brain disorders were over-represented in the genes associated with the OB_R volume compared with those not correlated with the OB_R volume. The NHGRI-EBI GWAS Catalog (MacArthur et al., 2017) was used to identify risk genes. Disorders with an immune response as the core mechanism or symptom were selected as immune-related disorders (**Table S1**). Brain disorders included schizophrenia, bipolar disorder, mood disorder, autism, and Alzheimer’s disease. Genes with p-values smaller than 1E-8 and an odd ratio larger than 1 from the GWAS were considered as risk genes. Over-representation was assessed using a chi-square test for the following 2×2 contingency table:

|  | Risk Gene | Not Risk Gene |
| --- | --- | --- |
| Correlated with the OB_R volume |  |  |
| Not correlated with the OB_R volume |  |  |

## 3. Results

### 3.1. Study participants

We initially collected the data of OB volumes and gene expressional profiles from 24 HC subjects and 24 FEP patients. In the present study, after excluding one outlier and individuals that self-reported smoking tobacco and cannabis, we analyzed the data from 22 HC subjects and 16 FEP patients. There were no significant differences in age, gender, race, and intracranial volume between HC subjects and FEP patients (**Table 1**). Consistent with our previous study (Kamath et al., 2018), we observed significant reductions in psychophysical olfactory performances (odor discrimination and odor identification) in FEP patients compared to HC subjects (**Table 1**).

**Table 1.** Demographic summary. Significant results are highlighted in bold. Abbreviations: FEP, first episode psychosis; HC, healthy control; sd, standard deviation; AA, African American; and Cauc, Caucasian.

| Characteristics | FEP (16) | HC (22) | P Value |
| --- | --- | --- | --- |
| Age<br>(*mean ± sd*) | 22.81 ± 4.03 | 23.50 ± 2.79 | 0.56 |
| Gender<br>(*male/female*) | 10/6 | 9/13 | 0.32 |
| Race<br>(*AA/Cauc/Asian/Hispanic/Others*) | 9/4/2/1/0 | 10/6/2/2/2 | 0.86 |
| Intracranial volume<br>(*mean ± sd*) | 1.28E6 ± 1.59E6 | 1.25E6 ± 1.08E6 | 0.50 |
| Odor discrimination<br>(*mean ± sd*) | 9.40 ± 2.44 | 11.71 ± 2.31 | **7.60E-03** |
| Odor identification<br>(*mean ± sd*) | 10.47 ± 2.13 | 11.95 ± 1.80 | **0.03** |

### 3.2. Alterations in OB structure in FEP patients compared with HC subjects

We observed a significant reduction in the volume of the total OB (p-value = 0.048) and OB_R (p-value = 0.030) in FEP patients compared to HC subjects (**Figure 1**). The difference in the OB_L volume between FEP patients and HC subjects didn’t reach significance (p-value = 0.245) (**Figure 1**). The significant difference observed in the total OB volume was mainly contributed by the changes in the OB_R.

**Figure 1.**
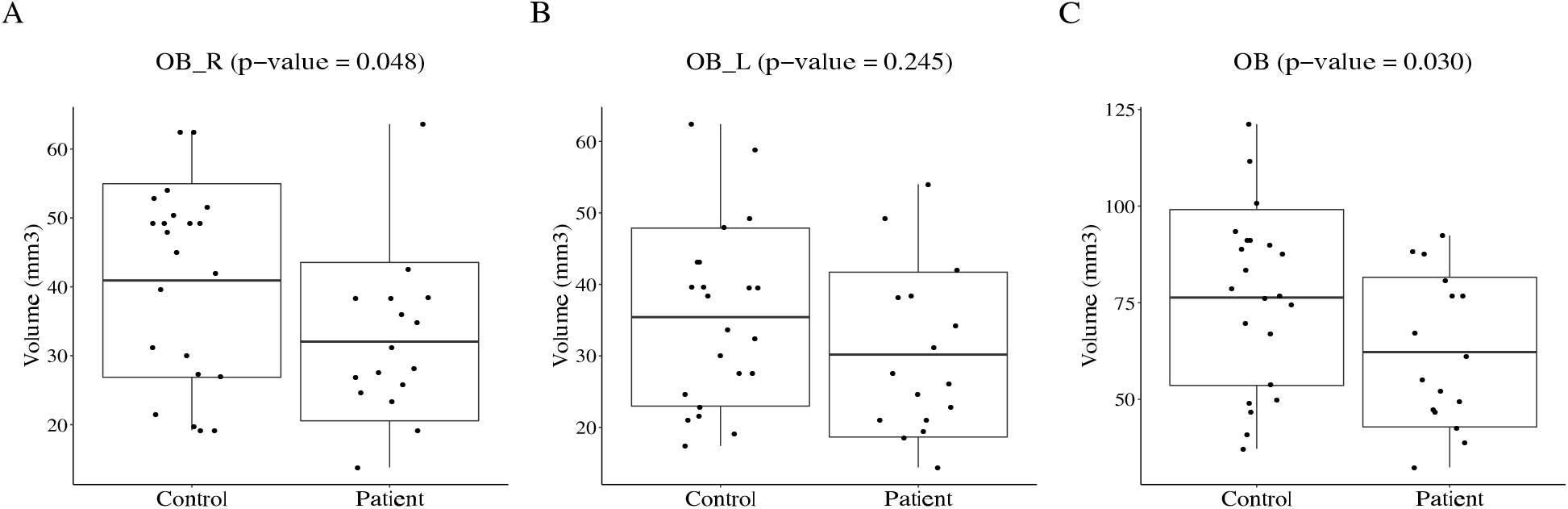
Boxplots of olfactory bulb (OB) volumes in FEP patients and healthy controls. We compared the volumes of the right OB (A), left OB (B), and entire OB (C) between FEP patients and healthy controls. After adjusting for potential confounding factors including age, gender, race, and smoking status, FEP patients were found to have a significantly smaller OB, especially in the right hemisphere, compared to healthy controls. The box represents the standard deviation, the solid line in the middle of the box shows the mean value of the volume. Each black dot represents an individual subject.

### 3.3. Alterations in gene expressional profiles in ONCs in FEP patients compared with HC subjects

Significant alterations in the gene expressional profiles in ONCs after multiple comparison correction were observed in FEP patients compared to HC subjects (**Figure S1**, **Table S2**). GSEA (Subramanian et al., 2005) on the Reactome Pathway Database (Fabregat et al., 2018) identified 138 significant pathways overrepresented in genes altered in FEP patients (**Table S3**). We next grouped these 138 pathways based on the parent-child hierarchical structure of pathways provided by the Reactome Pathway Database. Signal transduction, metabolism of proteins, and the immune system were found to be the top 3 parent networks with the most significant pathways (**Figure 2**).

**Figure 2.**
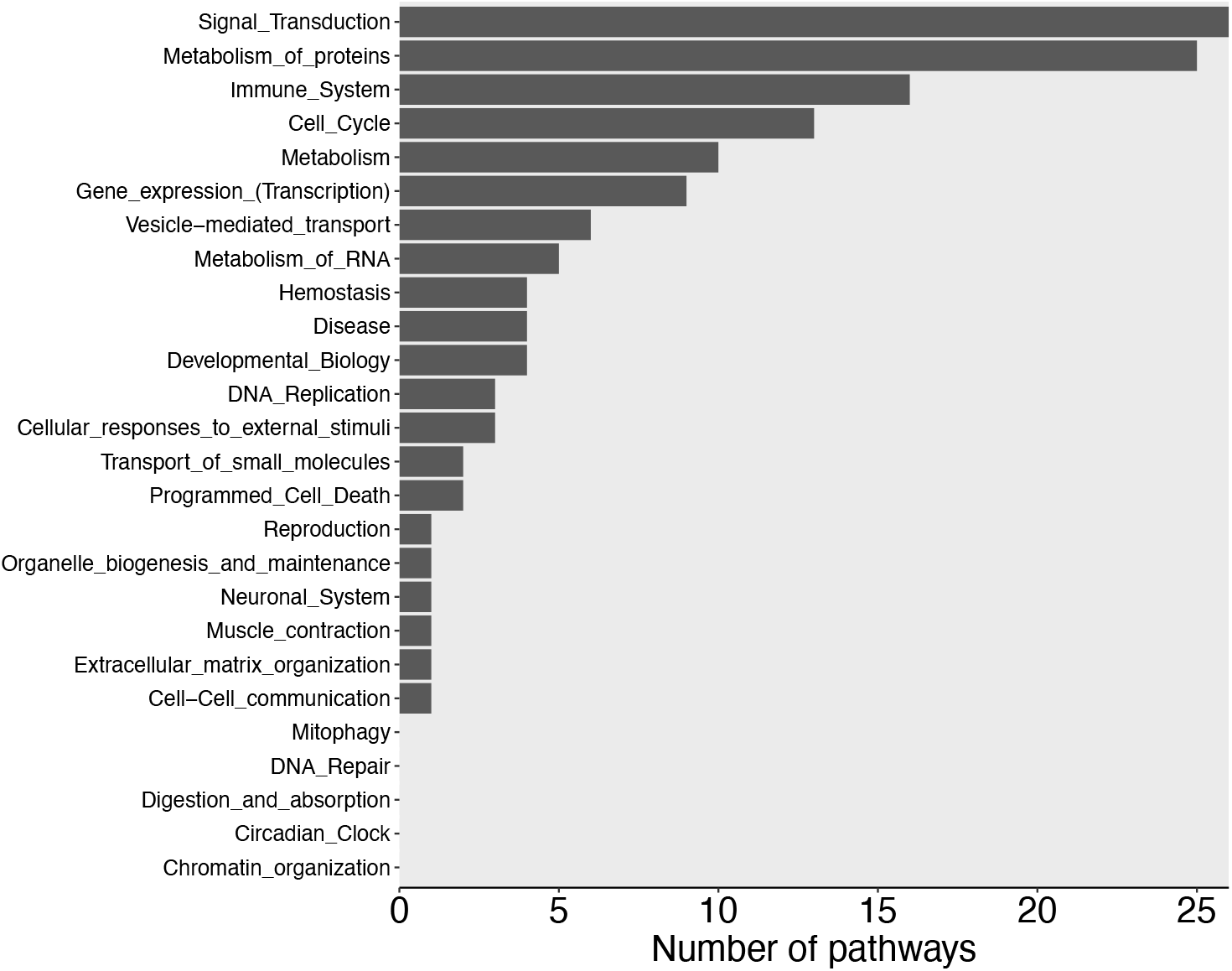
Gene set enrichment analysis (GSEA) results of genes altered in FEP patients compared to healthy controls. GSEA identified 138 significant pathways, which were further grouped based on the parent-child hierarchical structure of pathways provided by the Reactome Pathway Database. The y-axis has all the parent networks from Reactome Pathway Database and the x-axis showed the number of significant child pathways under the corresponding parent network.

### 3.4. Molecular pathways associated with the volume of the OB_R

After identifying a smaller OB volume, especially in the right hemisphere, in FEP patients, we next asked whether this shrinkage was associated with molecular pathways. General linear regression was performed to evaluate the correlations between gene expressional levels in ONCs and the OB_R volume. Since no correlation reached the significance cutoff after multiple comparison correction, we performed a permutation test. Genes with p-values less than 0.05 from both the general linear regression and permutation test were considered genes associated with the OB_R volume. Accordingly, we identified 1,197 (**Table S4**) and 646 (**Table S5**) genes associated with the OB_R volume in FEP patients and HC subjects, respectively. Enrichment analysis found that risk genes for immune-related disorders (p-value = 4.563E-07) (**Table 2**) and brain disorders (p-value = 1.564E-11) (**Table S6**) were significantly overrepresented in genes associated with the OB_R volume in FEP patients (see details about the definition of immune-related disorders and brain disorders in the Methods section). On the other hand, no overrepresentation of risk genes for immune-related disorders (p-value = 1) and brain disorders (p-value = 1) were observed in genes associated with the OB_R volume in HC subjects.

**Table 2.**
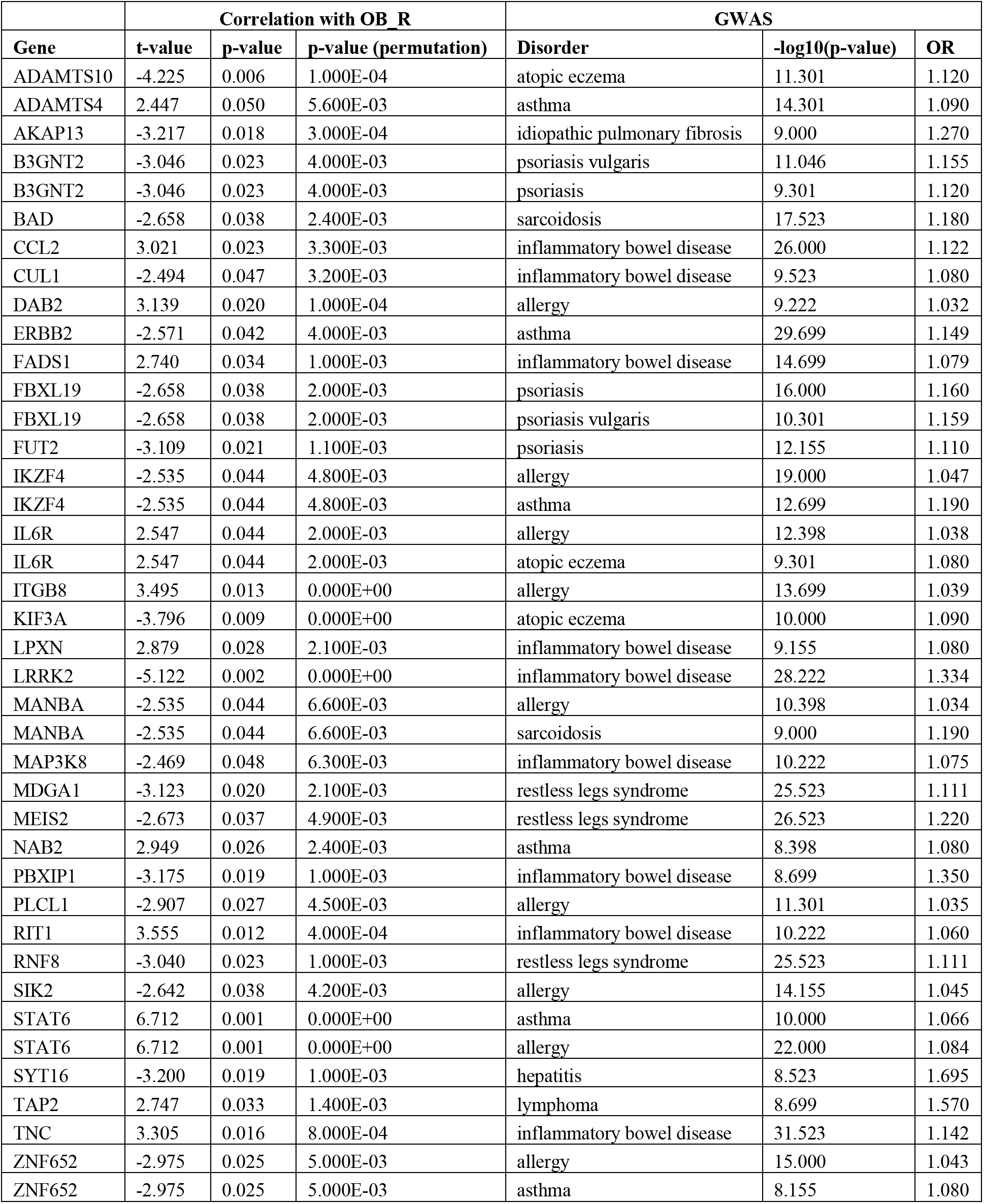
Genes that were significantly correlated with the right olfactory bulb (OB_R) volume in FEP patients and also identified by a genome-wide association study (GWAS) as risk genes for immune-related disorders.

We next performed gene set enrichment analysis (GSEA) (Subramanian et al., 2005) on the Reactome Pathway Database (Fabregat et al., 2018) and identified 88 significant pathways overrepresented in genes associated with the OB_R volume in FEP patients (**Table S7**). Metabolism of proteins, signal transduction, and the immune system were the top 3 parent networks with the most significant pathways (**Figure S2A**). In HC subjects, we identified 17 significant pathways associated with the OB_R volume; metabolism of proteins, metabolism of RNA, and organelle biogenesis and maintenance were the top 3 parent networks with the most significant pathways (**Figure S2B, Table S8**). Together with the differential expression analysis between FEP and HC groups, we found that immune-related pathways were altered in FEP patients and this alteration was associated with the change in the OB_R structure in patients.

## 4. Discussion

We have conducted an exploratory study to address a specific hypothesis to test whether specific pathophysiological events in the OE can account for possible alterations in the OB and overall olfactory dysfunction in FEP patients. Even in this small cohort, we could reproduce the observations reported by multiple groups that OB volume reduction (Asal et al., 2018; Nguyen et al., 2011; Turetsky et al., 2000) and olfactory functional deficits are associated with psychotic disorders (Kamath et al., 2018, 2019; Moberg et al., 2014). To address the pathophysiological events in the OE in FEP patients at the molecular levels, we conducted an unbiased expression study with olfactory neuronal cells freshly biopsied from FEP patients in comparison with those from HC subjects. In the molecular study, we observed that gene expression profiles were altered in FEP patients, where molecules for signal transduction, metabolism, and immune system were enriched as altered molecules. More importantly, among them, an alteration in immune-related molecules and pathways was further highlighted in correlation with the change of the OB volume. Given that many basic biology studies have supported OB volume changes as a result of the immune/inflammatory response in the OE (Hasegawa-Ishii et al., 2020), these results suggest that, in at least in a subset of SZ, the OB volume changes elicited by or associated with the immune/inflammatory response in the OE may exist as the disease pathophysiology.

Although the present exploratory study has provided supportive evidence to our working hypothesis on the unique role of OE and OB in the pathophysiology of FEP subjects, this also raised several important questions to be clarified in future studies. First, we observed a selective reduction in the OB volume on the right side. Possible asymmetry has been proposed as an outstanding question in olfactory deficits in psychotic disorders (Asal et al., 2018; Turetsky et al., 2003). There were studies support the right-specific alternation (Nguyen et al., 2011; Turetsky et al., 2003), but others did not find such laterality (Asal et al., 2018; Turetsky et al., 2000, 2018). Sex-associated changes in olfactory deficits have also been frequently discussed (Boesveldt et al., 2017; Chen et al., 2018b; Sorokowski et al., 2019). Possible differences between affective and non-affective psychosis in olfactory deficits are also an important question (Kamath et al., 2018). The sample size of the present study is too small to address these, and future studies are warranted.

Possible changes in OE pathology in psychotic disorders have been suggested (Arnold et al., 1998, 2001; Pantazopoulos et al., 2013; Smutzer et al., 1998). Nevertheless, the data in classic neuropathology have not been perfectly consistent (Arnold et al., 2001; Minovi et al., 2015), partly because of biological heterogeneity in the same diagnostic criteria. Several studies addressed possible molecular changes in postmortem OE, but all of them took, to the best of our knowledge, a candidate molecular approach and studied only a limited number of molecules (Pantazopoulos et al., 2013; Smutzer et al., 1998). More fundamentally, although all these postmortem OE studies are informative, it is important to figure out whether the pathological features observed in postmortem OE may represent the signatures of a long-term compensation to the key pathological events when psychosis occurs and is active. Accordingly, the use of biopsied OE samples for biological assessment has recently become active. Investigators have applied comprehensive molecular approaches, including unbiased RNA-seq and proteomic assessments, to neuronal cell cultures originating from biopsied OE, and report significant, pathological changes at the molecular level (Abrams et al., 1999; Brown et al., 2014; English et al., 2015, 2015; Evgrafov et al., 2020; Féron et al., 1999; Ghanbari et al., 2004; Girard et al., 2011; Mackay-Sim, 2012; Matigian et al., 2010; Mor et al., 2013; Rhie et al., 2018; Solís-Chagoyán et al., 2013; Wolozin et al., 1993). Nevertheless, these excellent studies with OE-derived neuronal cells were not directly linked to the data of olfactory neurocircuitry. In this sense, although we admit the exploratory nature of the present study, the present study provides a new avenue of studying the OE pathology in association with the OB volume, which is expected to affect the higher cortex.

Future studies may include the investigation of a relationship among adverse events in the nasal cavity, OE pathology at both the histological and molecular levels, structural/functional brain imaging data of the OB and upper neurocircuitry, as well as clinical phenotypes, including olfactory functional changes. To define cell-type specificity in a molecular study of the OE, the introduction of digital special profiling (although ONCs may represent profiles of progenitor cells) (Merritt et al., 2020) may also open a new window for schizophrenia/psychosis research.

