## Supplementary materials for "Volumetric alteration of olfactory bulb and immune-related molecular changes in olfactory epithelium in first episode psychosis patients"

### Table S1. Immune-related disorders for the GWAS enrichment analysis.

| Traits |
|---|
| acquired immunodeficiency syndrome (AIDS) |
| allergic rhinitis |
| allergy |
| amyloid light-chain (AL) amyloidosis |
| asthma |
| atopic eczema |
| B-cell acute lymphoblastic leukemia |
| cryoglobulinemia |
| dengue Hemorrhagic Fever |
| dilated cardiomyopathy |
| duodenal ulcer |
| hepatic fibrosis |
| hepatitis |
| hepatitis C induced liver cirrhosis |
| HIV-1 infection |
| hodgkins lymphoma |
| idiopathic pulmonary fibrosis |
| immune system disease |
| inflammatory bowel disease |
| influenza A (H1N1) |
| lymphoma |
| malaria |
| marginal zone B-cell lymphoma |
| mixed cellularity |
| monoclonal gammopathy |
| multiple myeloma |
| myositis |
| osteitis deformans |
| osteoarthritis |
| pancreatitis |
| periodontitis |
| psoriasis |
| psoriasis vulgaris |
| psoriatic arthritis |
| recalcitrant atopic dermatitis |
| restless legs syndrome |
| sarcoidosis |
| seasonal allergic rhinitis |
| staphylococcus aureus infection |
| Stevens-Johnson syndrome |
| susceptibility to viral and mycobacterial infections |
| toxic epidermal necrolysis |
| tuberculosis |
| visceral Leishmaniasis |
| Vogt-Koyanagi-Harada disease |

**Table S2. Differential expression analysis between first episode psychosis patients and healthy controls.**

R library *DESeq2* was used to perform differential expression analysis. Age, gender, race and the 3 hidden/unknown confounding factors identified by *sva* were included as covariates in the design formula. The Benjamini and Hochberg (BH) procedure was used for multiple comparison correction. The top 20 genes are listed in the table. The meaning of columns: baseMean, mean of normalized counts for all samples; lfcSE, standard error of the log2FoldChange; stat, Wald statistic, the log2FoldChange divided by its standard error; padj, p-value corrected for multiple testing.

| Gene | baseMean | log2FoldChange | lfcSE | stat | pvalue | padj |
|---|---|---|---|---|---|---|
| FAM155A | 174.909 | -3.435 | 0.543 | -6.324 | 2.551E-10 | 3.533E-06 |
| RAD9A | 1163.661 | 0.410 | 0.081 | 5.046 | 4.509E-07 | 3.123E-03 |
| CYTL1 | 467.084 | -3.838 | 0.805 | -4.767 | 1.874E-06 | 7.552E-03 |
| SLC1A2 | 48.914 | 1.993 | 0.421 | 4.736 | 2.181E-06 | 7.552E-03 |
| HMBS | 3354.686 | -0.294 | 0.067 | -4.383 | 1.170E-05 | 0.032 |
| KANSL1L | 1248.535 | 0.355 | 0.085 | 4.181 | 2.906E-05 | 0.057 |
| ST6GAL1 | 596.768 | -1.214 | 0.295 | -4.123 | 3.747E-05 | 0.057 |
| HAS3 | 3596.416 | -0.355 | 0.086 | -4.116 | 3.858E-05 | 0.057 |
| TANGO6 | 3596.416 | -0.355 | 0.086 | -4.116 | 3.858E-05 | 0.057 |
| YWHAE | 45723.339 | -0.331 | 0.081 | -4.082 | 4.470E-05 | 0.057 |
| ZNF560 | 66.755 | -2.010 | 0.494 | -4.067 | 4.763E-05 | 0.057 |
| SPOCD1 | 15777.072 | -0.909 | 0.224 | -4.060 | 4.897E-05 | 0.057 |
| SFRP4 | 158.752 | -4.229 | 1.072 | -3.944 | 8.000E-05 | 0.084 |
| PAQR8 | 1252.279 | 0.495 | 0.126 | 3.925 | 8.656E-05 | 0.084 |
| IGF2BP1 | 737.196 | -1.857 | 0.481 | -3.861 | 1.131E-04 | 0.084 |
| KIAA0391 | 13120.083 | -0.231 | 0.060 | -3.859 | 1.139E-04 | 0.084 |
| PSMA6 | 13120.083 | -0.231 | 0.060 | -3.859 | 1.139E-04 | 0.084 |
| ALDH4A1 | 18089.363 | -0.583 | 0.152 | -3.842 | 1.219E-04 | 0.084 |
| IFFO2 | 18089.363 | -0.583 | 0.152 | -3.842 | 1.219E-04 | 0.084 |
| TAS1R2 | 18089.363 | -0.583 | 0.152 | -3.842 | 1.219E-04 | 0.084 |

**Table S3. Significant pathways overrepresented in genes altered in first episode psychosis patients.**

Meaning of columns: size, the number of genes in the child pathway; NES, normalized enrichment score; FDR, false discovery rate.

| Child pathway | Parent network | size | NES | FDR |
|---|---|---|---|---|
| ABC-FAMILY PROTEINS MEDIATED TRANSPORT | Transport_of_small_molecules | 75 | -2.427 | 0.000 |
| ACTIVATED PKN1 STIMULATES TRANSCRIPTION OF AR (ANDROGEN RECEPTOR) REGULATED GENES KLK2 AND KLK3 | Signal_Transduction | 44 | -1.843 | 0.012 |
| ACTIVATION OF NF-KAPPAB IN B CELLS | Immune_System | 62 | -2.689 | 0.000 |
| ACTIVATION OF SMO | Signal_Transduction | 14 | 2.199 | 0.006 |
| ADP SIGNALLING THROUGH P2Y PURINOCEPTOR 1 | Hemostasis | 18 | -1.798 | 0.017 |
| ADP SIGNALLING THROUGH P2Y PURINOCEPTOR 12 | Hemostasis | 15 | -1.675 | 0.046 |
| AMYLOID FIBER FORMATION | Metabolism_of_proteins | 64 | -1.971 | 0.003 |
| ANTIGEN PRESENTATION: FOLDING; ASSEMBLY AND PEPTIDE LOADING OF CLASS I MHC | Immune_System | 25 | -2.109 | 0.000 |
| ANTIGEN PROCESSING: UBIQUITINATION & PROTEASOME DEGRADATION | Immune_System | 283 | -1.670 | 0.047 |
| APC/C:CDH1 MEDIATED DEGRADATION OF CDC20 AND OTHER APC/C:CDH1 TARGETED PROTEINS IN LATE MITOSIS/EARLY G1 | Cell_Cycle | 69 | -2.786 | 0.000 |
| ASSOCIATION OF TRIC/CCT WITH TARGET PROTEINS DURING BIOSYNTHESIS | Metabolism_of_proteins | 37 | -1.675 | 0.046 |
| ASYMMETRIC LOCALIZATION OF PCP PROTEINS | Signal_Transduction | 61 | -2.493 | 0.000 |
| AUF1 (HNRNP D0) BINDS AND DESTABILIZES MRNA | Metabolism_of_RNA | 52 | -2.810 | 0.000 |
| AUTODEGRADATION OF CDH1 BY CDH1:APC/C | Cell_Cycle | 62 | -2.636 | 0.000 |
| CDK-MEDIATED PHOSPHORYLATION AND REMOVAL OF CDC6 | DNA_Replication | 69 | -2.659 | 0.000 |
| CDT1 ASSOCIATION WITH THE CDC6:ORC:ORIGIN COMPLEX | DNA_Replication | 56 | -2.753 | 0.000 |
| CELL SURFACE INTERACTIONS AT THE VASCULAR WALL | Hemostasis | 58 | -1.763 | 0.023 |

| | | | | |
|---|---|---|---|---|
| CELL-EXTRACELLULAR MATRIX INTERACTIONS | Cell-Cell_communication | 12 | -1.800 | 0.017 |
| CHK1/CHK2(CDS1) MEDIATED INACTIVATION OF CYCLIN B:CDK1 COMPLEX | Cell_Cycle | 13 | -1.851 | 0.011 |
| CILIUM ASSEMBLY | Organelle_biogenesis_and_maintenance | 14 | -2.015 | 0.001 |
| CITRIC ACID CYCLE (TCA CYCLE) | Metabolism | 22 | -1.993 | 0.002 |
| CLEC7A (DECTIN-1) SIGNALING | Immune_System | 73 | -2.616 | 0.000 |
| COMPLEX I BIOGENESIS | Metabolism | 49 | -2.234 | 0.000 |
| CONDENSATION OF PROPHASE CHROMOSOMES | Cell_Cycle | 52 | -1.826 | 0.014 |
| COOPERATION OF PDCL (PHLP1) AND TRIC/CCT IN G-PROTEIN BETA FOLDING | Metabolism_of_proteins | 34 | -2.083 | 0.000 |
| COPI-DEPENDENT GOLGI-TO-ER RETROGRADE TRAFFIC | Vesicle-mediated_transport | 88 | -2.207 | 0.000 |
| COPI-INDEPENDENT GOLGI-TO-ER RETROGRADE TRAFFIC | Vesicle-mediated_transport | 45 | -2.220 | 0.000 |
| COPI-MEDIATED ANTEROGRADE TRANSPORT | Vesicle-mediated_transport | 93 | -2.376 | 0.000 |
| COPII-MEDIATED VESICLE TRANSPORT | Vesicle-mediated_transport | 65 | -1.905 | 0.006 |
| CROSS-PRESENTATION OF SOLUBLE EXOGENOUS ANTIGENS (ENDOSOMES) | Immune_System | 43 | -2.724 | 0.000 |
| CROSSLINKING OF COLLAGEN FIBRILS | Extracellular_matrix_organization | 18 | -1.675 | 0.046 |
| CYCLIN A/B1/B2 ASSOCIATED EVENTS DURING G2/M TRANSITION | Cell_Cycle | 25 | -1.802 | 0.016 |
| DECTIN-1 MEDIATED NONCANONICAL NF-KB SIGNALING | Immune_System | 57 | -2.818 | 0.000 |
| DEFECTIVE CFTR CAUSES CYSTIC FIBROSIS | Disease | 57 | -2.633 | 0.000 |
| DEGRADATION OF AXIN | Signal_Transduction | 52 | -2.712 | 0.000 |
| DEGRADATION OF BETA-CATENIN BY THE DESTRUCTION COMPLEX | Signal_Transduction | 65 | -2.680 | 0.000 |
| DEGRADATION OF DVL | Signal_Transduction | 54 | -2.644 | 0.000 |
| DEGRADATION OF GLI1 BY THE PROTEASOME | Signal_Transduction | 56 | -2.607 | 0.000 |
| DEGRADATION OF GLI2 BY THE PROTEASOME | Signal_Transduction | 56 | -2.603 | 0.000 |
| DEPOSITION OF NEW CENPA-CONTAINING NUCLEOSOMES AT THE CENTROMERE | Cell_Cycle | 55 | -1.769 | 0.022 |

| | | | | |
|---|---|---|---|---|
| DETOXIFICATION OF REACTIVE OXYGEN SPECIES | Cellular_responses_to_external_stimuli | 30 | -1.727 | 0.031 |
| DNA METHYLATION | Gene_expression_(Transcription) | 43 | -1.957 | 0.003 |
| DOWNSTREAM TCR SIGNALING | Immune_System | 78 | -2.581 | 0.000 |
| ENOS ACTIVATION | Metabolism | 11 | -1.819 | 0.014 |
| EPHB-MEDIATED FORWARD SIGNALING | Developmental_Biology | 41 | -1.876 | 0.008 |
| ER-PHAGOSOME PATHWAY | Immune_System | 79 | -2.766 | 0.000 |
| EUKARYOTIC TRANSLATION TERMINATION | Metabolism_of_proteins | 92 | -2.894 | 0.000 |
| FBXL7 DOWN-REGULATES AURKA DURING MITOTIC ENTRY AND IN EARLY MITOSIS | Cell_Cycle | 52 | -2.801 | 0.000 |
| FCERI MEDIATED NF-KB ACTIVATION | Immune_System | 74 | -2.576 | 0.000 |
| FOLDING OF ACTIN BY CCT/TRIC | Metabolism_of_proteins | 10 | -2.052 | 0.001 |
| FORMATION OF A POOL OF FREE 40S SUBUNITS | Metabolism_of_proteins | 100 | -2.924 | 0.000 |
| FORMATION OF THE TERNARY COMPLEX; AND SUBSEQUENTLY; THE 43S COMPLEX | Metabolism_of_proteins | 51 | -2.339 | 0.000 |
| FORMATION OF TUBULIN FOLDING INTERMEDIATES BY CCT/TRIC | Metabolism_of_proteins | 21 | -2.439 | 0.000 |
| G ALPHA (Z) SIGNALLING EVENTS | Signal_Transduction | 36 | -1.830 | 0.013 |
| G BETA:GAMMA SIGNALLING THROUGH PI3KGAMMA | Signal_Transduction | 32 | -1.790 | 0.018 |
| G BETA:GAMMA SIGNALLING THROUGH PLC BETA | Signal_Transduction | 15 | -1.811 | 0.015 |
| G2/M CHECKPOINTS | Cell_Cycle | 48 | -2.794 | 0.000 |
| GAP JUNCTION ASSEMBLY | Vesicle-mediated_transport | 19 | -1.947 | 0.003 |
| GENERIC TRANSCRIPTION PATHWAY | Gene_expression_(Transcription) | 327 | 2.052 | 0.023 |
| GLI3 IS PROCESSED TO GLI3R BY THE PROTEASOME | Signal_Transduction | 56 | -2.549 | 0.000 |
| GLUCAGON SIGNALING IN METABOLIC REGULATION | Metabolism | 19 | -1.669 | 0.047 |
| GLUCAGON-TYPE LIGAND RECEPTORS | Signal_Transduction | 15 | -1.691 | 0.042 |
| GLUCONEOGENESIS | Metabolism | 25 | -1.821 | 0.014 |
| GLYOXYLATE METABOLISM AND GLYCINE DEGRADATION | Metabolism | 23 | -1.693 | 0.041 |
| GTP HYDROLYSIS AND JOINING OF THE 60S RIBOSOMAL SUBUNIT | Metabolism_of_proteins | 111 | -2.977 | 0.000 |

| | | | | |
|---|---|---|---|---|
| HEDGEHOG 'ON' STATE | Signal_Transduction | 67 | -2.394 | 0.000 |
| HEDGEHOG LIGAND BIOGENESIS | Signal_Transduction | 54 | -2.664 | 0.000 |
| HH MUTANTS THAT DON'T UNDERGO AUTOCATALYTIC PROCESSING ARE DEGRADED BY ERAD | Disease | 52 | -2.684 | 0.000 |
| HSP90 CHAPERONE CYCLE FOR STEROID HORMONE RECEPTORS (SHR) | Cellular_responses_to_external_stimuli | 49 | -2.226 | 0.000 |
| INTERLEUKIN-1 SIGNALING | Immune_System | 81 | -2.495 | 0.000 |
| KERATINIZATION | Developmental_Biology | 35 | -1.951 | 0.003 |
| L13A-MEDIATED TRANSLATIONAL SILENCING OF CERULOPLASMIN EXPRESSION | Metabolism_of_proteins | 110 | -2.958 | 0.000 |
| LDL CLEARANCE | Transport_of_small_molecules | 18 | -2.095 | 0.000 |
| MAJOR PATHWAY OF RRNA PROCESSING IN THE NUCLEOLUS AND CYTOSOL | Metabolism_of_RNA | 177 | -2.844 | 0.000 |
| MAPK6/MAPK4 SIGNALING | Signal_Transduction | 84 | -2.286 | 0.000 |
| MEIOTIC RECOMBINATION | Cell_Cycle | 60 | -1.700 | 0.039 |
| MEIOTIC RECOMBINATION | Reproduction | 60 | -1.700 | 0.039 |
| MHC CLASS II ANTIGEN PRESENTATION | Immune_System | 104 | -2.124 | 0.000 |
| MITOCHONDRIAL PROTEIN IMPORT | Metabolism_of_proteins | 60 | -2.273 | 0.000 |
| MITOCHONDRIAL TRANSLATION ELONGATION | Metabolism_of_proteins | 86 | -1.931 | 0.004 |
| MITOCHONDRIAL TRANSLATION INITIATION | Metabolism_of_proteins | 86 | -1.975 | 0.002 |
| MITOCHONDRIAL TRANSLATION TERMINATION | Metabolism_of_proteins | 86 | -1.990 | 0.002 |
| MITOTIC PROMETAPHASE | Cell_Cycle | 101 | -1.686 | 0.043 |
| NEDDYLATION | Metabolism_of_proteins | 215 | -1.695 | 0.040 |
| NEUTROPHIL DEGRANULATION | Immune_System | 374 | -2.167 | 0.000 |
| NIK: NONCANONICAL NF-KB SIGNALING | Immune_System | 56 | -2.814 | 0.000 |
| NONSENSE MEDIATED DECAY (NMD) ENHANCED BY THE EXON JUNCTION COMPLEX (EJC) | Metabolism_of_RNA | 114 | -2.729 | 0.000 |
| NONSENSE MEDIATED DECAY (NMD) INDEPENDENT OF THE EXON JUNCTION COMPLEX (EJC) | Metabolism_of_RNA | 94 | -2.874 | 0.000 |
| O-LINKED GLYCOSYLATION OF MUCINS | Metabolism_of_proteins | 31 | -1.678 | 0.045 |
| ORC1 REMOVAL FROM CHROMATIN | DNA_Replication | 68 | -2.577 | 0.000 |

| | | | | |
|---|---|---|---|---|
| OXYGEN-DEPENDENT PROLINE HYDROXYLATION OF HYPOXIA-INDUCIBLE FACTOR ALPHA | Cellular_responses_to_external_stimuli | 61 | -2.501 | 0.000 |
| PEPTIDE CHAIN ELONGATION | Metabolism_of_proteins | 88 | -2.921 | 0.000 |
| POST-CHAPERONIN TUBULIN FOLDING PATHWAY | Metabolism_of_proteins | 18 | -1.717 | 0.034 |
| PRC2 METHYLATES HISTONES AND DNA | Gene_expression_(Transcription) | 52 | -1.737 | 0.029 |
| PREFOLDIN MEDIATED TRANSFER OF SUBSTRATE TO CCT/TRIC | Metabolism_of_proteins | 26 | -2.479 | 0.000 |
| PRESYNAPTIC FUNCTION OF KAINATE RECEPTORS | Neuronal_System | 15 | -1.853 | 0.011 |
| PROSTACYCLIN SIGNALLING THROUGH PROSTACYCLIN RECEPTOR | Hemostasis | 14 | -1.708 | 0.037 |
| PROTEIN METHYLATION | Metabolism_of_proteins | 14 | -1.863 | 0.010 |
| PYRUVATE METABOLISM | Metabolism | 17 | -2.139 | 0.000 |
| RECYCLING PATHWAY OF L1 | Developmental_Biology | 40 | -2.192 | 0.000 |
| REGULATION OF ACTIN DYNAMICS FOR PHAGOCYTIC CUP FORMATION | Immune_System | 52 | -2.024 | 0.001 |
| REGULATION OF ACTIVATED PAK-2P34 BY PROTEASOME MEDIATED DEGRADATION | Programmed_Cell_Death | 47 | -2.779 | 0.000 |
| REGULATION OF EXPRESSION OF SLITS AND ROBOS | Developmental_Biology | 160 | -2.859 | 0.000 |
| REGULATION OF ORNITHINE DECARBOXYLASE (ODC) | Metabolism | 48 | -2.674 | 0.000 |
| REGULATION OF PTEN STABILITY AND ACTIVITY | Signal_Transduction | 66 | -2.569 | 0.000 |
| REGULATION OF RAS BY GAPS | Signal_Transduction | 63 | -2.447 | 0.000 |
| REGULATION OF RUNX2 EXPRESSION AND ACTIVITY | Gene_expression_(Transcription) | 68 | -2.478 | 0.000 |
| REGULATION OF RUNX3 EXPRESSION AND ACTIVITY | Gene_expression_(Transcription) | 54 | -2.841 | 0.000 |
| RESPIRATORY ELECTRON TRANSPORT | Metabolism | 78 | -2.691 | 0.000 |
| RHO GTPASES ACTIVATE CIT | Signal_Transduction | 18 | -1.893 | 0.007 |
| RHO GTPASES ACTIVATE IQGAPS | Signal_Transduction | 25 | -2.266 | 0.000 |
| RHO GTPASES ACTIVATE PAKS | Signal_Transduction | 20 | -2.068 | 0.001 |
| RHO GTPASES ACTIVATE PKNS | Signal_Transduction | 27 | -1.965 | 0.003 |
| RHO GTPASES ACTIVATE ROCKS | Signal_Transduction | 18 | -1.815 | 0.015 |
| RHO GTPASES ACTIVATE WASPS AND WAVES | Signal_Transduction | 34 | -1.767 | 0.022 |

| | | | | |
|---|---|---|---|---|
| RIBOSOMAL SCANNING AND START CODON RECOGNITION | Metabolism_of_proteins | 58 | -2.488 | 0.000 |
| RIP-MEDIATED NFKB ACTIVATION VIA ZBP1 | Immune_System | 20 | -1.777 | 0.020 |
| RIPK1-MEDIATED REGULATED NECROSIS | Programmed_Cell_Death | 15 | -1.701 | 0.039 |
| RNA POLYMERASE I CHAIN ELONGATION | Gene_expression_(Transcription) | 69 | -1.690 | 0.042 |
| RNA POLYMERASE I PROMOTER OPENING | Gene_expression_(Transcription) | 42 | -1.806 | 0.016 |
| RRNA MODIFICATION IN THE NUCLEUS AND CYTOSOL | Metabolism_of_RNA | 57 | -2.200 | 0.000 |
| RUNX1 REGULATES TRANSCRIPTION OF GENES INVOLVED IN DIFFERENTIATION OF HSCS | Gene_expression_(Transcription) | 99 | -2.672 | 0.000 |
| SCF(SKP2)-MEDIATED DEGRADATION OF P27/P21 | Cell_Cycle | 59 | -2.694 | 0.000 |
| SELENOCYSTEINE SYNTHESIS | Metabolism | 92 | -2.791 | 0.000 |
| SIRT1 NEGATIVELY REGULATES RRNA EXPRESSION | Gene_expression_(Transcription) | 47 | -1.849 | 0.011 |
| SMOOTH MUSCLE CONTRACTION | Muscle_contraction | 32 | -2.157 | 0.000 |
| SRP-DEPENDENT COTRANSLATIONAL PROTEIN TARGETING TO MEMBRANE | Metabolism_of_proteins | 111 | -2.886 | 0.000 |
| SYNTHESIS OF ACTIVE UBIQUITIN: ROLES OF E1 AND E2 ENZYMES | Metabolism_of_proteins | 30 | -2.145 | 0.000 |
| THE ROLE OF GTSE1 IN G2/M PROGRESSION AFTER G2 CHECKPOINT | Cell_Cycle | 69 | -3.034 | 0.000 |
| TNFR2 NON-CANONICAL NF-KB PATHWAY | Immune_System | 60 | -2.751 | 0.000 |
| TRANSLOCATION OF GLUT4 TO THE PLASMA MEMBRANE | Vesicle-mediated_transport | 64 | -2.230 | 0.000 |
| UB-SPECIFIC PROCESSING PROTEASES | Metabolism_of_proteins | 173 | -2.290 | 0.000 |
| UBIQUITIN-DEPENDENT DEGRADATION OF CYCLIN D1 | Cell_Cycle | 49 | -2.679 | 0.000 |
| UCH PROTEINASES | Metabolism_of_proteins | 92 | -2.494 | 0.000 |
| VEGFA-VEGFR2 PATHWAY | Signal_Transduction | 54 | -1.680 | 0.045 |
| VIF-MEDIATED DEGRADATION OF APOBEC3G | Disease | 49 | -2.864 | 0.000 |
| VPU MEDIATED DEGRADATION OF CD4 | Disease | 49 | -2.738 | 0.000 |
| WNT5A-DEPENDENT INTERNALIZATION OF FZD2; FZD5 AND ROR2 | Signal_Transduction | 13 | -1.751 | 0.026 |

**Table S4. Genes associated with the right olfactory bulb volume in first episode psychosis patients.**
Abbreviations: GLR, generalized linear regression

| Gene | t-value (GLR) | p-value (GLR) | p-value (permutation) |
|---|---|---|---|
| CASP4 | 7.713 | 0.000 | 0.000 |
| COX10-AS1 | -7.138 | 0.000 | 0.000 |
| CKMT2 | -7.056 | 0.000 | 0.000 |
| RP11-342K6.2 | -6.895 | 0.000 | 0.000 |
| STAT6 | 6.712 | 0.001 | 0.000 |
| MORN4 | -6.203 | 0.001 | 0.000 |
| ACSF2 | -6.190 | 0.001 | 0.000 |
| FCGRT | 5.978 | 0.001 | 0.000 |
| RCN3 | 5.978 | 0.001 | 0.000 |
| PARP16 | -5.958 | 0.001 | 0.000 |
| BCLAF1P2 | 5.875 | 0.001 | 0.000 |
| HMGN1P13 | -5.808 | 0.001 | 0.000 |
| ZNRF3 | -5.591 | 0.001 | 0.000 |
| OXCT2 | 5.486 | 0.002 | 0.000 |
| AC079145.4 | -5.430 | 0.002 | 0.000 |
| DYNC2LI1 | -5.418 | 0.002 | 0.000 |
| FLJ13224 | -5.403 | 0.002 | 0.000 |
| PRDX6 | 5.392 | 0.002 | 0.000 |
| UBE2I | 5.266 | 0.002 | 0.000 |
| FAM229B | -5.216 | 0.002 | 0.000 |
| ARPP19 | -5.154 | 0.002 | 0.000 |
| ZFYVE28 | -5.132 | 0.002 | 0.000 |
| LRRK2 | -5.122 | 0.002 | 0.000 |
| UQCR10 | 5.120 | 0.002 | 0.000 |
| MECR | 5.088 | 0.002 | 0.000 |
| KCTD2 | -5.036 | 0.002 | 0.000 |
| FAM109B | -5.030 | 0.002 | 0.000 |
| RP11-474D14.2 | 5.008 | 0.002 | 0.000 |
| VWA1 | -4.979 | 0.003 | 0.000 |
| ZNF532 | -4.979 | 0.003 | 0.000 |
| MGP | -4.947 | 0.003 | 0.000 |
| DDX56 | 4.941 | 0.003 | 0.000 |
| TMED4 | 4.941 | 0.003 | 0.000 |
| ZNF57 | -4.932 | 0.003 | 0.000 |
| EGFR | 4.915 | 0.003 | 0.000 |
| CACNA1H | -4.878 | 0.003 | 0.000 |
| INO80B | -4.856 | 0.003 | 0.000 |
| INO80B-WBP1 | -4.856 | 0.003 | 0.000 |
| WBP1 | -4.856 | 0.003 | 0.000 |
| HMGN4 | 4.844 | 0.003 | 0.000 |
| CH17-140K24.8 | -4.805 | 0.003 | 0.000 |
| PCDHB14 | -4.805 | 0.003 | 0.000 |
| PCDHB15 | -4.805 | 0.003 | 0.000 |

| | | | |
|---|---|---|---|
| SMIM10L2B | -4.788 | 0.003 | 0.000 |
| TXNRD1 | 4.774 | 0.003 | 0.000 |
| SNU13 | 4.705 | 0.003 | 0.000 |
| AMZ2P1 | -4.683 | 0.003 | 0.000 |
| RP3-390M24.1 | 4.629 | 0.004 | 0.000 |
| SDHAF4 | -4.601 | 0.004 | 0.000 |
| CNOT6L | 4.598 | 0.004 | 0.000 |
| HES6 | -4.597 | 0.004 | 0.000 |
| AC011290.5 | 4.563 | 0.004 | 0.000 |
| ADRBK2 | -4.559 | 0.004 | 0.000 |
| CH17-140K24.4 | -4.543 | 0.004 | 0.000 |
| CH17-140K24.5 | -4.543 | 0.004 | 0.000 |
| PCDHB10 | -4.543 | 0.004 | 0.000 |
| PCDHB16 | -4.543 | 0.004 | 0.000 |
| PCDHB8 | -4.543 | 0.004 | 0.000 |
| PCDHB9 | -4.543 | 0.004 | 0.000 |
| NRP1 | 4.499 | 0.004 | 0.000 |
| CTC-550B14.7 | -4.495 | 0.004 | 0.000 |
| AC007228.8 | -4.469 | 0.004 | 0.000 |
| ZNF71 | -4.469 | 0.004 | 0.000 |
| AGBL5-IT1 | -4.395 | 0.005 | 0.000 |
| RP11-69H7.3 | -4.344 | 0.005 | 0.000 |
| BRD9 | 4.338 | 0.005 | 0.000 |
| ZDHHC11 | 4.338 | 0.005 | 0.000 |
| LANCL1 | -4.329 | 0.005 | 0.000 |
| ACKR2 | 4.325 | 0.005 | 0.000 |
| FAM198A | 4.325 | 0.005 | 0.000 |
| RP11-141M3.6 | 4.325 | 0.005 | 0.000 |
| RP11-67L2.2 | -4.304 | 0.005 | 0.000 |
| NUTM2D | -4.272 | 0.005 | 0.000 |
| AD001527.7 | 4.249 | 0.005 | 0.000 |
| ARF4-AS1 | -4.240 | 0.005 | 0.000 |
| GPR75-ASB3 | -4.239 | 0.005 | 0.000 |
| LIMA1 | 4.238 | 0.005 | 0.000 |
| FAM78A | -4.232 | 0.005 | 0.000 |
| FLOT1 | -4.177 | 0.006 | 0.000 |
| XXbac-BPG252P9.9 | -4.177 | 0.006 | 0.000 |
| AC090587.4 | -4.169 | 0.006 | 0.000 |
| MIR4687 | -4.169 | 0.006 | 0.000 |
| PWWP2B | -4.130 | 0.006 | 0.000 |
| ATP6AP2 | -4.130 | 0.006 | 0.000 |
| ST7-OT3_1 | -4.118 | 0.006 | 0.000 |
| ST7-OT4 | -4.118 | 0.006 | 0.000 |
| ST7-OT4_2 | -4.118 | 0.006 | 0.000 |
| SNN | -4.073 | 0.007 | 0.000 |
| HENMT1 | -4.051 | 0.007 | 0.000 |
| IQSEC1 | -4.009 | 0.007 | 0.000 |
| EIF2B5 | 4.003 | 0.007 | 0.000 |
| CTD-2528L19.4 | -3.988 | 0.007 | 0.000 |

| | | | |
|---|---|---|---|
| ZFP30 | -3.988 | 0.007 | 0.000 |
| ZNF781 | -3.988 | 0.007 | 0.000 |
| CNKSR1 | 3.983 | 0.007 | 0.000 |
| TSPAN6 | -3.962 | 0.007 | 0.000 |
| APBA2 | -3.925 | 0.008 | 0.000 |
| BAIAP2-AS1 | -3.875 | 0.008 | 0.000 |
| STARD13 | 3.869 | 0.008 | 0.000 |
| RAD23A | 3.839 | 0.009 | 0.000 |
| KIF3A | -3.796 | 0.009 | 0.000 |
| SEC13 | 3.747 | 0.010 | 0.000 |
| TMEM136 | 3.747 | 0.010 | 0.000 |
| PXMP4 | -3.674 | 0.010 | 0.000 |
| HHAT | -3.636 | 0.011 | 0.000 |
| PTPRA | -3.613 | 0.011 | 0.000 |
| FAM120C | 3.576 | 0.012 | 0.000 |
| C1orf101 | -3.536 | 0.012 | 0.000 |
| CCDC112 | -3.527 | 0.012 | 0.000 |
| CNIH1 | -3.506 | 0.013 | 0.000 |
| ITGB8 | 3.495 | 0.013 | 0.000 |
| RP3-430N8.10 | -3.476 | 0.013 | 0.000 |
| GHR | -3.455 | 0.014 | 0.000 |
| ZNF589 | 3.433 | 0.014 | 0.000 |
| FAM167A | 3.430 | 0.014 | 0.000 |
| SNORD118 | -3.398 | 0.015 | 0.000 |
| TMEM107 | -3.398 | 0.015 | 0.000 |
| PHF7 | -3.398 | 0.015 | 0.000 |
| ZNF322 | -3.385 | 0.015 | 0.000 |
| DBNL | 3.384 | 0.015 | 0.000 |
| FAM66C | -3.275 | 0.017 | 0.000 |
| CTD-2224J9.4 | -3.186 | 0.019 | 0.000 |
| SLC46A3 | -3.039 | 0.023 | 0.000 |
| MBOAT2 | -4.949 | 0.003 | 1.00E-04 |
| RP11-175I6.1 | 4.920 | 0.003 | 1.00E-04 |
| MPLKIP | -4.678 | 0.003 | 1.00E-04 |
| SHANK3 | -4.600 | 0.004 | 1.00E-04 |
| CASP6 | -4.586 | 0.004 | 1.00E-04 |
| PLA2G12A | -4.586 | 0.004 | 1.00E-04 |
| ROGDI | -4.497 | 0.004 | 1.00E-04 |
| NNT-AS1 | -4.411 | 0.005 | 1.00E-04 |
| SLC40A1 | -4.330 | 0.005 | 1.00E-04 |
| RP11-136C24.3 | 4.325 | 0.005 | 1.00E-04 |
| ZNF662 | 4.325 | 0.005 | 1.00E-04 |
| ATP2B1 | -4.292 | 0.005 | 1.00E-04 |
| LINC00863 | -4.272 | 0.005 | 1.00E-04 |
| AC092574.2 | 4.240 | 0.005 | 1.00E-04 |
| ADAMTS10 | -4.225 | 0.006 | 1.00E-04 |
| IER3 | -4.177 | 0.006 | 1.00E-04 |
| STIM1 | -4.169 | 0.006 | 1.00E-04 |
| PDZD4 | -4.160 | 0.006 | 1.00E-04 |

| | | | |
|---|---|---|---|
| ST7 | -4.118 | 0.006 | 1.00E-04 |
| ST7-AS1_2 | -4.118 | 0.006 | 1.00E-04 |
| ST7-OT3_3 | -4.118 | 0.006 | 1.00E-04 |
| ST7-OT4_4 | -4.118 | 0.006 | 1.00E-04 |
| ANKRA2 | -4.030 | 0.007 | 1.00E-04 |
| METTL22 | 4.017 | 0.007 | 1.00E-04 |
| ZNF607 | -3.988 | 0.007 | 1.00E-04 |
| ZNF593 | 3.983 | 0.007 | 1.00E-04 |
| SNORD69 | 3.957 | 0.007 | 1.00E-04 |
| NUDT16L1 | 3.951 | 0.008 | 1.00E-04 |
| NT5DC1 | -3.939 | 0.008 | 1.00E-04 |
| ARHGEF6 | -3.935 | 0.008 | 1.00E-04 |
| FBXO3 | -3.915 | 0.008 | 1.00E-04 |
| TMEM246 | -3.773 | 0.009 | 1.00E-04 |
| C1GALT1C1L | -3.767 | 0.009 | 1.00E-04 |
| LOXL1-AS1 | 3.758 | 0.009 | 1.00E-04 |
| UGDH-AS1 | -3.740 | 0.010 | 1.00E-04 |
| ABCC2 | 3.739 | 0.010 | 1.00E-04 |
| RP11-465N4.5 | 3.722 | 0.010 | 1.00E-04 |
| KDELC1 | -3.707 | 0.010 | 1.00E-04 |
| SNX16 | -3.677 | 0.010 | 1.00E-04 |
| ACP2 | -3.652 | 0.011 | 1.00E-04 |
| TXNL4A | 3.622 | 0.011 | 1.00E-04 |
| CLHC1 | -3.613 | 0.011 | 1.00E-04 |
| VPS16 | -3.613 | 0.011 | 1.00E-04 |
| RP11-21K12.2 | 3.612 | 0.011 | 1.00E-04 |
| SSTR1 | 3.569 | 0.012 | 1.00E-04 |
| SH3BGR | -3.542 | 0.012 | 1.00E-04 |
| RP11-815J21.4 | -3.537 | 0.012 | 1.00E-04 |
| RP11-225H22.4 | -3.528 | 0.012 | 1.00E-04 |
| ECSCR | -3.472 | 0.013 | 1.00E-04 |
| TNK1 | -3.470 | 0.013 | 1.00E-04 |
| PARP3 | 3.431 | 0.014 | 1.00E-04 |
| KIZ | -3.429 | 0.014 | 1.00E-04 |
| DBNDD1 | -3.406 | 0.014 | 1.00E-04 |
| AC074117.10 | 3.401 | 0.014 | 1.00E-04 |
| RP11-299J3.8 | 3.397 | 0.015 | 1.00E-04 |
| AK3P3 | -3.342 | 0.016 | 1.00E-04 |
| ZNF426 | -3.329 | 0.016 | 1.00E-04 |
| RP11-345K20.2 | -3.327 | 0.016 | 1.00E-04 |
| EML6 | -3.288 | 0.017 | 1.00E-04 |
| RP3-437C15.1 | 3.283 | 0.017 | 1.00E-04 |
| PFN1 | 3.249 | 0.017 | 1.00E-04 |
| MIPEPP3 | -3.235 | 0.018 | 1.00E-04 |
| TSPO | 3.231 | 0.018 | 1.00E-04 |
| RP11-5C23.1 | -3.227 | 0.018 | 1.00E-04 |
| MPV17 | 3.193 | 0.019 | 1.00E-04 |
| MGAM | -3.180 | 0.019 | 1.00E-04 |
| DAB2 | 3.139 | 0.020 | 1.00E-04 |

| | | | |
|---|---|---|---|
| CYP4V2 | -3.135 | 0.020 | 1.00E-04 |
| MMD | -4.613 | 0.004 | 2.00E-04 |
| OXCT2P1 | 4.612 | 0.004 | 2.00E-04 |
| KRBOX1 | 4.325 | 0.005 | 2.00E-04 |
| ASB3 | -4.239 | 0.005 | 2.00E-04 |
| GPR75 | -4.239 | 0.005 | 2.00E-04 |
| CPPED1 | 4.212 | 0.006 | 2.00E-04 |
| RP11-312J18.5 | 4.199 | 0.006 | 2.00E-04 |
| ST7-OT3_2 | -4.118 | 0.006 | 2.00E-04 |
| KB-1027C11.4 | -3.998 | 0.007 | 2.00E-04 |
| PRH1-PRR4 | -3.917 | 0.008 | 2.00E-04 |
| NCAM2 | 3.910 | 0.008 | 2.00E-04 |
| RPS11P6 | -3.867 | 0.008 | 2.00E-04 |
| INA | -3.844 | 0.009 | 2.00E-04 |
| AS3MT | -3.833 | 0.009 | 2.00E-04 |
| BORCS7-ASMT | -3.833 | 0.009 | 2.00E-04 |
| KHDRBS3 | 3.729 | 0.010 | 2.00E-04 |
| ANKRD19P | -3.675 | 0.010 | 2.00E-04 |
| CAPN2 | 3.577 | 0.012 | 2.00E-04 |
| CTD-3220F14.3 | -3.525 | 0.012 | 2.00E-04 |
| DNAJC18 | -3.472 | 0.013 | 2.00E-04 |
| KNSTRN | 3.450 | 0.014 | 2.00E-04 |
| COPZ1 | 3.443 | 0.014 | 2.00E-04 |
| MIR6775 | 3.434 | 0.014 | 2.00E-04 |
| CDC40 | -3.407 | 0.014 | 2.00E-04 |
| AC017116.11 | 3.384 | 0.015 | 2.00E-04 |
| CD276 | -3.370 | 0.015 | 2.00E-04 |
| MRPL27 | 3.290 | 0.017 | 2.00E-04 |
| UCN | 3.193 | 0.019 | 2.00E-04 |
| R3HDM1 | 3.191 | 0.019 | 2.00E-04 |
| ZNF813 | -3.186 | 0.019 | 2.00E-04 |
| KLKB1 | -3.135 | 0.020 | 2.00E-04 |
| RP11-515O17.3 | -4.613 | 0.004 | 3.00E-04 |
| RP11-475I24.1 | 4.293 | 0.005 | 3.00E-04 |
| NOX4 | -4.273 | 0.005 | 3.00E-04 |
| MIR6132 | -4.118 | 0.006 | 3.00E-04 |
| ST7-OT4_1 | -4.118 | 0.006 | 3.00E-04 |
| ST7-OT4_3 | -4.118 | 0.006 | 3.00E-04 |
| PNCK | -4.115 | 0.006 | 3.00E-04 |
| HIC2 | -4.108 | 0.006 | 3.00E-04 |
| SLC2A1-AS1 | -4.056 | 0.007 | 3.00E-04 |
| RP11-667F14.1 | -4.013 | 0.007 | 3.00E-04 |
| RP11-734K2.4 | -3.987 | 0.007 | 3.00E-04 |
| BMS1P3 | 3.951 | 0.008 | 3.00E-04 |
| FAM117B | -3.875 | 0.008 | 3.00E-04 |
| BORCS7 | -3.833 | 0.009 | 3.00E-04 |
| MIR6777 | 3.786 | 0.009 | 3.00E-04 |
| BMF | -3.755 | 0.009 | 3.00E-04 |
| OAS2 | 3.711 | 0.010 | 3.00E-04 |

| | | | |
|---|---|---|---|
| LRRC1 | -3.711 | 0.010 | 3.00E-04 |
| RP5-1160K1.6 | 3.626 | 0.011 | 3.00E-04 |
| SGCE | -3.572 | 0.012 | 3.00E-04 |
| GJA1 | -3.558 | 0.012 | 3.00E-04 |
| RP11-230B22.1 | -3.512 | 0.013 | 3.00E-04 |
| ATP6AP1 | -3.489 | 0.013 | 3.00E-04 |
| MARCH1 | 3.453 | 0.014 | 3.00E-04 |
| RABEPK | 3.442 | 0.014 | 3.00E-04 |
| SENP7 | -3.394 | 0.015 | 3.00E-04 |
| MIR6837 | 3.384 | 0.015 | 3.00E-04 |
| BEX1 | 3.343 | 0.016 | 3.00E-04 |
| SCML2P2 | -3.326 | 0.016 | 3.00E-04 |
| CHRDL1 | 3.293 | 0.017 | 3.00E-04 |
| SPATA4 | -3.292 | 0.017 | 3.00E-04 |
| CTD-2033A16.1 | 3.283 | 0.017 | 3.00E-04 |
| HNRNPA3P12 | 3.274 | 0.017 | 3.00E-04 |
| NQO1 | 3.252 | 0.017 | 3.00E-04 |
| MT-TQ | 3.248 | 0.018 | 3.00E-04 |
| AKAP13 | -3.217 | 0.018 | 3.00E-04 |
| KRT10 | 3.215 | 0.018 | 3.00E-04 |
| CTD-2337A12.1 | 3.190 | 0.019 | 3.00E-04 |
| C9 | 3.139 | 0.020 | 3.00E-04 |
| CTC-339F2.2 | -3.103 | 0.021 | 3.00E-04 |
| LSM11 | -3.035 | 0.023 | 3.00E-04 |
| MVB12B | -3.012 | 0.024 | 3.00E-04 |
| RP11-430B1.2 | -3.000 | 0.024 | 3.00E-04 |
| KANTR | -4.116 | 0.006 | 4.00E-04 |
| RP11-258C19.7 | -3.836 | 0.009 | 4.00E-04 |
| C3orf67 | -3.788 | 0.009 | 4.00E-04 |
| RIT1 | 3.555 | 0.012 | 4.00E-04 |
| ATG4C | -3.512 | 0.013 | 4.00E-04 |
| AC004967.7 | 3.503 | 0.013 | 4.00E-04 |
| AAMDC | -3.418 | 0.014 | 4.00E-04 |
| RP11-496H1.2 | -3.392 | 0.015 | 4.00E-04 |
| LINC00957 | 3.384 | 0.015 | 4.00E-04 |
| SCD | 3.354 | 0.015 | 4.00E-04 |
| ARHGEF25 | -3.336 | 0.016 | 4.00E-04 |
| CLEC4A | -3.275 | 0.017 | 4.00E-04 |
| MIR7706 | -3.217 | 0.018 | 4.00E-04 |
| RP11-463C8.4 | -3.187 | 0.019 | 4.00E-04 |
| TMEM63C | -3.187 | 0.019 | 4.00E-04 |
| VPS9D1-AS1 | 3.172 | 0.019 | 4.00E-04 |
| RP11-87H9.5 | -3.111 | 0.021 | 4.00E-04 |
| SF3A3 | 3.049 | 0.023 | 4.00E-04 |
| ITM2B | -4.363 | 0.005 | 5.00E-04 |
| ARHGAP42 | -4.078 | 0.007 | 5.00E-04 |
| KCNIP4 | -4.057 | 0.007 | 5.00E-04 |
| AGAP11 | 3.951 | 0.008 | 5.00E-04 |
| RP11-96C23.13 | 3.951 | 0.008 | 5.00E-04 |

| | | | |
|---|---|---|---|
| BAZ2B | -3.865 | 0.008 | 5.00E-04 |
| BX322557.10 | -3.798 | 0.009 | 5.00E-04 |
| MIR647 | 3.741 | 0.010 | 5.00E-04 |
| GBP4 | 3.730 | 0.010 | 5.00E-04 |
| RP11-36C20.1 | 3.557 | 0.012 | 5.00E-04 |
| AC010976.2 | 3.544 | 0.012 | 5.00E-04 |
| NSDHL | 3.516 | 0.013 | 5.00E-04 |
| OR7E38P | 3.503 | 0.013 | 5.00E-04 |
| BEX2 | 3.343 | 0.016 | 5.00E-04 |
| WFS1 | -3.343 | 0.016 | 5.00E-04 |
| KBTBD3 | -3.339 | 0.016 | 5.00E-04 |
| SLC26A10 | -3.336 | 0.016 | 5.00E-04 |
| DZIP3 | -3.333 | 0.016 | 5.00E-04 |
| TRIM61 | -3.210 | 0.018 | 5.00E-04 |
| KLF17P2 | 3.205 | 0.018 | 5.00E-04 |
| LINC01085 | 3.205 | 0.018 | 5.00E-04 |
| SNRPA | 3.138 | 0.020 | 5.00E-04 |
| TRAF3IP2-AS1 | -3.124 | 0.020 | 5.00E-04 |
| RP11-195O1.5 | -3.027 | 0.023 | 5.00E-04 |
| PITRM1 | 3.006 | 0.024 | 5.00E-04 |
| RNF14 | -3.001 | 0.024 | 5.00E-04 |
| MANEA | -2.973 | 0.025 | 5.00E-04 |
| RP11-350N15.4 | 2.897 | 0.027 | 5.00E-04 |
| EDNRA | -3.817 | 0.009 | 6.00E-04 |
| LINC00205 | -3.798 | 0.009 | 6.00E-04 |
| PCDHB7 | -3.697 | 0.010 | 6.00E-04 |
| RP11-46H11.3 | -3.589 | 0.012 | 6.00E-04 |
| SRP72 | 3.468 | 0.013 | 6.00E-04 |
| SIRPB2 | 3.372 | 0.015 | 6.00E-04 |
| IFI27L2 | 3.353 | 0.015 | 6.00E-04 |
| CASK | -3.347 | 0.015 | 6.00E-04 |
| VLDLR-AS1 | -3.263 | 0.017 | 6.00E-04 |
| FNBP1L | -3.220 | 0.018 | 6.00E-04 |
| CFAP36 | -3.208 | 0.018 | 6.00E-04 |
| RP11-669M16.2 | 3.205 | 0.018 | 6.00E-04 |
| RP5-1061H20.4 | -3.161 | 0.020 | 6.00E-04 |
| USP27X | -3.141 | 0.020 | 6.00E-04 |
| AQP3 | -3.141 | 0.020 | 6.00E-04 |
| FANK1 | -3.090 | 0.021 | 6.00E-04 |
| AZIN2 | -3.066 | 0.022 | 6.00E-04 |
| CTB-58E17.5 | -2.990 | 0.024 | 6.00E-04 |
| MYPN | 2.983 | 0.025 | 6.00E-04 |
| C22orf23 | -2.973 | 0.025 | 6.00E-04 |
| SEH1L | 2.941 | 0.026 | 6.00E-04 |
| RP3-331H24.7 | 2.939 | 0.026 | 6.00E-04 |
| DCTPP1 | 2.931 | 0.026 | 6.00E-04 |
| AP001437.1 | 2.927 | 0.026 | 6.00E-04 |
| AGA | 4.335 | 0.005 | 7.00E-04 |
| RP11-96C23.14 | 3.951 | 0.008 | 7.00E-04 |

| | | | |
|---|---|---|---|
| RAB11B-AS1 | -3.727 | 0.010 | 7.00E-04 |
| MDM1 | -3.559 | 0.012 | 7.00E-04 |
| ZNF354A | -3.420 | 0.014 | 7.00E-04 |
| ISCA1P4 | -3.414 | 0.014 | 7.00E-04 |
| RP11-47I22.4 | -3.392 | 0.015 | 7.00E-04 |
| AC139887.4 | 3.379 | 0.015 | 7.00E-04 |
| NSFL1C | 3.372 | 0.015 | 7.00E-04 |
| PPP2R1A | 3.326 | 0.016 | 7.00E-04 |
| AKAP6 | -3.290 | 0.017 | 7.00E-04 |
| RP11-366M4.11 | -3.210 | 0.018 | 7.00E-04 |
| RP5-1061H20.3 | -3.161 | 0.020 | 7.00E-04 |
| ZBTB44 | -3.107 | 0.021 | 7.00E-04 |
| TES | -3.092 | 0.021 | 7.00E-04 |
| RP11-278C7.5 | -3.017 | 0.023 | 7.00E-04 |
| VPS37D | -2.988 | 0.024 | 7.00E-04 |
| AC005944.2 | 2.962 | 0.025 | 7.00E-04 |
| CAPNS1 | 2.923 | 0.027 | 7.00E-04 |
| MPST | 2.923 | 0.027 | 7.00E-04 |
| MECP2 | -2.888 | 0.028 | 7.00E-04 |
| XXbac-BPG246D15.9 | 2.747 | 0.033 | 7.00E-04 |
| RP11-676J12.7 | 4.798 | 0.003 | 8.00E-04 |
| ADIRF | 3.951 | 0.008 | 8.00E-04 |
| TAS2R12 | -3.917 | 0.008 | 8.00E-04 |
| TAS2R14 | -3.917 | 0.008 | 8.00E-04 |
| AC009506.2 | -3.865 | 0.008 | 8.00E-04 |
| LINC01232 | -3.858 | 0.008 | 8.00E-04 |
| ZNF696 | -3.743 | 0.010 | 8.00E-04 |
| LGMN | -3.696 | 0.010 | 8.00E-04 |
| YIF1A | 3.555 | 0.012 | 8.00E-04 |
| RP11-190A12.8 | 3.471 | 0.013 | 8.00E-04 |
| PHYH | -3.406 | 0.014 | 8.00E-04 |
| SQLE | 3.399 | 0.015 | 8.00E-04 |
| APLP1 | -3.397 | 0.015 | 8.00E-04 |
| TNC | 3.305 | 0.016 | 8.00E-04 |
| ZNF705A | -3.275 | 0.017 | 8.00E-04 |
| DNAJA1 | 3.272 | 0.017 | 8.00E-04 |
| LIPE-AS1 | 3.203 | 0.019 | 8.00E-04 |
| CIPC | -3.187 | 0.019 | 8.00E-04 |
| AC066694.1 | -3.172 | 0.019 | 8.00E-04 |
| TRPC1 | -3.150 | 0.020 | 8.00E-04 |
| SOD2 | 3.125 | 0.020 | 8.00E-04 |
| CTSF | -2.997 | 0.024 | 8.00E-04 |
| RP11-422P24.12 | -2.971 | 0.025 | 8.00E-04 |
| EIF5AP4 | 2.841 | 0.030 | 8.00E-04 |
| MRPS14 | 2.804 | 0.031 | 8.00E-04 |
| PTBP1 | 2.626 | 0.039 | 8.00E-04 |
| CASKIN1 | -3.730 | 0.010 | 9.00E-04 |
| DIMT1 | 3.489 | 0.013 | 9.00E-04 |
| ZFP90 | -3.466 | 0.013 | 9.00E-04 |

| | | | |
|---|---|---|---|
| FLJ22447 | -3.392 | 0.015 | 9.00E-04 |
| GEM | -3.308 | 0.016 | 9.00E-04 |
| NEGR1 | 3.307 | 0.016 | 9.00E-04 |
| CTPS2 | -3.169 | 0.019 | 9.00E-04 |
| RNF20 | -3.166 | 0.019 | 9.00E-04 |
| INSIG1 | 3.135 | 0.020 | 9.00E-04 |
| RP11-517C16.2 | 3.129 | 0.020 | 9.00E-04 |
| RAB40A | -3.053 | 0.022 | 9.00E-04 |
| RP11-1398P2.1 | -2.986 | 0.024 | 9.00E-04 |
| RP11-330H6.5 | 2.962 | 0.025 | 9.00E-04 |
| TWF2 | 2.962 | 0.025 | 9.00E-04 |
| KLHDC2 | -2.889 | 0.028 | 9.00E-04 |
| TMCO3 | -2.843 | 0.029 | 9.00E-04 |
| RP5-1112D6.7 | -2.839 | 0.030 | 9.00E-04 |
| C20orf96 | -2.837 | 0.030 | 9.00E-04 |
| RGMB-AS1 | 2.824 | 0.030 | 9.00E-04 |
| DDX60L | 2.764 | 0.033 | 9.00E-04 |
| ARPC1A | 2.759 | 0.033 | 9.00E-04 |
| HNRNPA3P5 | 2.663 | 0.037 | 9.00E-04 |
| RNF5 | -2.591 | 0.041 | 9.00E-04 |
| HS3ST3A1 | 4.179 | 0.006 | 0.001 |
| PRH1 | -3.917 | 0.008 | 0.001 |
| XXbac-BPG283O16.9 | -3.459 | 0.013 | 0.001 |
| TMEM214 | 3.426 | 0.014 | 0.001 |
| PRKCH | -3.392 | 0.015 | 0.001 |
| SLC30A4 | -3.355 | 0.015 | 0.001 |
| AC234582.1 | -3.309 | 0.016 | 0.001 |
| RBBP9 | -3.292 | 0.017 | 0.001 |
| UBQLNL | -3.235 | 0.018 | 0.001 |
| RP11-330M2.4 | -3.224 | 0.018 | 0.001 |
| SYT16 | -3.200 | 0.019 | 0.001 |
| PBXIP1 | -3.175 | 0.019 | 0.001 |
| COL3A1 | -3.172 | 0.019 | 0.001 |
| COPG1 | 3.114 | 0.021 | 0.001 |
| NCBP3 | 3.049 | 0.023 | 0.001 |
| RNF8 | -3.040 | 0.023 | 0.001 |
| TMEM182 | -2.993 | 0.024 | 0.001 |
| SLC25A22 | 2.946 | 0.026 | 0.001 |
| CKMT2-AS1 | -2.861 | 0.029 | 0.001 |
| RP3-331H24.5 | 2.794 | 0.031 | 0.001 |
| FADS1 | 2.740 | 0.034 | 0.001 |
| ZNF235 | -2.699 | 0.036 | 0.001 |
| FAM25A | 3.951 | 0.008 | 0.001 |
| ZNF517 | -3.541 | 0.012 | 0.001 |
| C16orf87 | -3.519 | 0.013 | 0.001 |
| FSCN1 | 3.318 | 0.016 | 0.001 |
| AC234582.2 | -3.309 | 0.016 | 0.001 |
| CTD-2277K2.1 | -3.200 | 0.019 | 0.001 |
| TAF4 | -3.189 | 0.019 | 0.001 |

| | | | |
|---|---|---|---|
| FUT2 | -3.109 | 0.021 | 0.001 |
| PCYOX1 | -3.067 | 0.022 | 0.001 |
| ANAPC13 | 3.033 | 0.023 | 0.001 |
| CCSAP | -3.021 | 0.023 | 0.001 |
| RP5-1042K10.13 | 3.019 | 0.023 | 0.001 |
| CTA-941F9.10 | 3.006 | 0.024 | 0.001 |
| RP11-452L6.1 | 2.902 | 0.027 | 0.001 |
| CADPS2 | 2.882 | 0.028 | 0.001 |
| ALG5 | 2.836 | 0.030 | 0.001 |
| TSEN2 | 2.835 | 0.030 | 0.001 |
| RP11-154D6.1 | 2.794 | 0.031 | 0.001 |
| ARPC1B | 2.759 | 0.033 | 0.001 |
| MIR1908 | 2.740 | 0.034 | 0.001 |
| GCNT2 | -2.723 | 0.035 | 0.001 |
| MIR4745 | 2.626 | 0.039 | 0.001 |
| KCNK2 | 4.368 | 0.005 | 0.001 |
| NUTM2A | -3.829 | 0.009 | 0.001 |
| S100A11 | 3.528 | 0.012 | 0.001 |
| MSRA | 3.207 | 0.018 | 0.001 |
| TMEM59L | -3.135 | 0.020 | 0.001 |
| AGAP2-AS1 | 3.106 | 0.021 | 0.001 |
| C20orf202 | 3.070 | 0.022 | 0.001 |
| MIR339 | 3.045 | 0.023 | 0.001 |
| MEN1 | 3.011 | 0.024 | 0.001 |
| RPS18P9 | -2.991 | 0.024 | 0.001 |
| AC007318.5 | 2.921 | 0.027 | 0.001 |
| LLNLR-268E12.1 | -2.824 | 0.030 | 0.001 |
| LINC00472 | 2.794 | 0.031 | 0.001 |
| NME5 | -2.750 | 0.033 | 0.001 |
| MRPS25 | 2.647 | 0.038 | 0.001 |
| RAB40B | -2.561 | 0.043 | 0.001 |
| CTD-2184D3.6 | -2.539 | 0.044 | 0.001 |
| CBY3 | -3.709 | 0.010 | 0.001 |
| ANXA5 | 3.506 | 0.013 | 0.001 |
| NLRC3 | -3.306 | 0.016 | 0.001 |
| PCDHB13 | -3.239 | 0.018 | 0.001 |
| C1orf52 | -3.129 | 0.020 | 0.001 |
| TMSB10 | 2.994 | 0.024 | 0.001 |
| RCBTB1 | -2.993 | 0.024 | 0.001 |
| ANTXR1 | -2.954 | 0.025 | 0.001 |
| AC009963.4 | 2.876 | 0.028 | 0.001 |
| TOMM34 | 2.839 | 0.030 | 0.001 |
| TMEM175 | -2.700 | 0.036 | 0.001 |
| IMP4 | 2.659 | 0.038 | 0.001 |
| RPL23AP4 | -3.653 | 0.011 | 0.001 |
| RPP38 | -3.564 | 0.012 | 0.001 |
| CACFD1 | -3.435 | 0.014 | 0.001 |
| RPS6P25 | 3.315 | 0.016 | 0.001 |
| TFDP2 | -3.138 | 0.020 | 0.001 |

| | | | |
|---|---|---|---|
| RP11-20I23.10 | 3.110 | 0.021 | 0.001 |
| XXYLT1 | -3.055 | 0.022 | 0.001 |
| DTNA | -2.994 | 0.024 | 0.001 |
| RP11-56B16.4 | 2.975 | 0.025 | 0.001 |
| TLR9 | 2.962 | 0.025 | 0.001 |
| DHCR24 | 2.936 | 0.026 | 0.001 |
| KDM4D | -2.921 | 0.027 | 0.001 |
| RPS2P55 | 2.886 | 0.028 | 0.001 |
| TP53TG1 | -2.867 | 0.029 | 0.001 |
| AC021224.1 | 2.862 | 0.029 | 0.001 |
| RP13-131K19.6 | -2.848 | 0.029 | 0.001 |
| CTC-559E9.5 | -2.811 | 0.031 | 0.001 |
| RP5-877J2.1 | -2.786 | 0.032 | 0.001 |
| RP11-134G8.7 | -2.769 | 0.032 | 0.001 |
| TAP2 | 2.747 | 0.033 | 0.001 |
| PMS2 | -2.722 | 0.035 | 0.001 |
| SUSD5 | -2.697 | 0.036 | 0.001 |
| CD109 | 2.674 | 0.037 | 0.001 |
| DERA | 2.639 | 0.039 | 0.001 |
| RP11-96C23.11 | 3.951 | 0.008 | 0.002 |
| RP11-295D4.3 | 3.451 | 0.014 | 0.002 |
| AGBL5 | 3.426 | 0.014 | 0.002 |
| HECTD2 | -3.344 | 0.016 | 0.002 |
| NUTM2B | -3.319 | 0.016 | 0.002 |
| LMLN | -3.163 | 0.019 | 0.002 |
| WWP1 | -3.129 | 0.020 | 0.002 |
| UBL7-AS1 | -3.086 | 0.021 | 0.002 |
| RP4-545L17.12 | 3.070 | 0.022 | 0.002 |
| C15orf65 | -2.994 | 0.024 | 0.002 |
| AC005037.3 | 2.955 | 0.025 | 0.002 |
| AC096772.6 | -2.936 | 0.026 | 0.002 |
| DAAM2 | -2.933 | 0.026 | 0.002 |
| XXyac-YR38GF2.1 | -2.933 | 0.026 | 0.002 |
| RP11-493P1.2 | -2.928 | 0.026 | 0.002 |
| NEK5 | -2.927 | 0.026 | 0.002 |
| XPR1 | -2.923 | 0.027 | 0.002 |
| PPFIA3 | -2.905 | 0.027 | 0.002 |
| DDB1 | 2.877 | 0.028 | 0.002 |
| RNF215 | -2.876 | 0.028 | 0.002 |
| MPP2 | -2.874 | 0.028 | 0.002 |
| ELP5 | 2.838 | 0.030 | 0.002 |
| RP11-110I1.13 | 2.815 | 0.031 | 0.002 |
| EIF1B | -2.810 | 0.031 | 0.002 |
| KRTAP1-1 | 2.807 | 0.031 | 0.002 |
| LIX1L | 2.750 | 0.033 | 0.002 |
| RP11-525A16.4 | 2.740 | 0.034 | 0.002 |
| PRR4 | -3.917 | 0.008 | 0.002 |
| ADAMTS13 | -3.435 | 0.014 | 0.002 |
| C8orf37 | -3.415 | 0.014 | 0.002 |

| | | | |
|---|---|---|---|
| PRKCZ | -3.406 | 0.014 | 0.002 |
| C10orf88 | -3.263 | 0.017 | 0.002 |
| THAP9 | -3.198 | 0.019 | 0.002 |
| MIR1257 | -3.189 | 0.019 | 0.002 |
| IGHMBP2 | -3.158 | 0.020 | 0.002 |
| PTH1R | -3.055 | 0.022 | 0.002 |
| RANP1 | 3.052 | 0.022 | 0.002 |
| RN7SL273P | -3.040 | 0.023 | 0.002 |
| LARS2 | 3.016 | 0.024 | 0.002 |
| RAVER2 | -2.927 | 0.026 | 0.002 |
| MAP3K3 | -2.892 | 0.028 | 0.002 |
| FAR1 | -2.889 | 0.028 | 0.002 |
| TP53INP1 | -2.814 | 0.031 | 0.002 |
| ZNF618 | -2.725 | 0.034 | 0.002 |
| DAZAP1 | 2.722 | 0.035 | 0.002 |
| SEPT9 | 2.651 | 0.038 | 0.002 |
| SNORA36A | 2.630 | 0.039 | 0.002 |
| SNORA56 | 2.630 | 0.039 | 0.002 |
| CTD-2008P7.6 | 2.610 | 0.040 | 0.002 |
| ARMCX5-GPRASP2 | -2.584 | 0.042 | 0.002 |
| PI4KA | 3.085 | 0.022 | 0.002 |
| RP11-575F12.2 | 3.053 | 0.022 | 0.002 |
| CAAP1 | -3.033 | 0.023 | 0.002 |
| CCDC7 | -3.027 | 0.023 | 0.002 |
| CTD-3126B10.5 | 2.971 | 0.025 | 0.002 |
| PDK3 | -2.930 | 0.026 | 0.002 |
| PAXIP1-AS2 | -2.890 | 0.028 | 0.002 |
| NFIC | 2.846 | 0.029 | 0.002 |
| ANKRD34A | 2.750 | 0.033 | 0.002 |
| TRIO | 2.694 | 0.036 | 0.002 |
| ZC3H8 | -2.691 | 0.036 | 0.002 |
| ARHGEF37 | -3.542 | 0.012 | 0.002 |
| SPRY1 | -3.376 | 0.015 | 0.002 |
| EPPK1 | -3.275 | 0.017 | 0.002 |
| PPP1R14BP3 | 3.190 | 0.019 | 0.002 |
| ATXN10 | 3.006 | 0.024 | 0.002 |
| MEG9 | 2.958 | 0.025 | 0.002 |
| GS1-358P8.4 | -2.930 | 0.026 | 0.002 |
| TBC1D32 | -2.908 | 0.027 | 0.002 |
| HFE | -2.856 | 0.029 | 0.002 |
| MIR4530 | -2.794 | 0.031 | 0.002 |
| INMT-FAM188B | -2.786 | 0.032 | 0.002 |
| RP1-239B22.5 | 2.774 | 0.032 | 0.002 |
| AP001205.1 | 2.770 | 0.032 | 0.002 |
| TTC39A | -2.744 | 0.034 | 0.002 |
| MMP16 | -2.714 | 0.035 | 0.002 |
| SIM2 | -2.700 | 0.036 | 0.002 |
| SETD1A | -2.658 | 0.038 | 0.002 |
| MIR664B | 2.630 | 0.039 | 0.002 |

| | | | |
|---|---|---|---|
| NF2 | 2.610 | 0.040 | 0.002 |
| WNK3 | -3.650 | 0.011 | 0.002 |
| DDB2 | -3.552 | 0.012 | 0.002 |
| PCSK5 | -3.332 | 0.016 | 0.002 |
| RNFT1 | -3.252 | 0.017 | 0.002 |
| UTP15 | 2.993 | 0.024 | 0.002 |
| VASP | 2.954 | 0.025 | 0.002 |
| MTIF3 | -2.910 | 0.027 | 0.002 |
| CMC2 | 2.900 | 0.027 | 0.002 |
| RP3-510D11.4 | -2.887 | 0.028 | 0.002 |
| PWP2 | 2.874 | 0.028 | 0.002 |
| SRM | 2.861 | 0.029 | 0.002 |
| TCTN2 | -2.854 | 0.029 | 0.002 |
| NPR2 | -2.833 | 0.030 | 0.002 |
| LL0XNC01-116E7.2 | 2.807 | 0.031 | 0.002 |
| RP11-338K17.8 | -2.799 | 0.031 | 0.002 |
| SPRTN | -2.797 | 0.031 | 0.002 |
| SNTA1 | -2.796 | 0.031 | 0.002 |
| INMT | -2.786 | 0.032 | 0.002 |
| DCAF5 | -2.779 | 0.032 | 0.002 |
| RSPH10B | -2.722 | 0.035 | 0.002 |
| AC135048.13 | -2.658 | 0.038 | 0.002 |
| DKC1 | 2.630 | 0.039 | 0.002 |
| PIPOX | -2.546 | 0.044 | 0.002 |
| TNFAIP2 | 3.731 | 0.010 | 0.002 |
| CCDC3 | -3.629 | 0.011 | 0.002 |
| SMARCA1 | -3.549 | 0.012 | 0.002 |
| RP11-307L3.2 | 3.366 | 0.015 | 0.002 |
| RP11-1060J15.4 | -3.135 | 0.020 | 0.002 |
| XX-C00717C00720L.1 | -3.078 | 0.022 | 0.002 |
| SEPT8 | -2.999 | 0.024 | 0.002 |
| ELOVL2 | 2.959 | 0.025 | 0.002 |
| PELO | -2.937 | 0.026 | 0.002 |
| KLC4 | -2.936 | 0.026 | 0.002 |
| RNF44 | -2.913 | 0.027 | 0.002 |
| KLHL29 | -2.912 | 0.027 | 0.002 |
| CABLES2 | -2.838 | 0.030 | 0.002 |
| ZC3H6 | -2.795 | 0.031 | 0.002 |
| AQP1 | -2.786 | 0.032 | 0.002 |
| FAM188B | -2.786 | 0.032 | 0.002 |
| ENTPD1 | -2.750 | 0.033 | 0.002 |
| CCNE1 | -2.722 | 0.035 | 0.002 |
| MCU | -2.709 | 0.035 | 0.002 |
| COG2 | -2.695 | 0.036 | 0.002 |
| LA16c-358B7.3 | 2.687 | 0.036 | 0.002 |
| ADH5 | 2.674 | 0.037 | 0.002 |
| FBXL19 | -2.658 | 0.038 | 0.002 |
| RP1-30M3.5 | 2.653 | 0.038 | 0.002 |
| TEX9 | -2.628 | 0.039 | 0.002 |

| | | | |
|---|---|---|---|
| IL6R | 2.547 | 0.044 | 0.002 |
| DHX40P1 | -3.252 | 0.017 | 0.002 |
| MDGA1 | -3.123 | 0.020 | 0.002 |
| G6PD | 3.077 | 0.022 | 0.002 |
| PSMF1 | 3.070 | 0.022 | 0.002 |
| TMEM256-PLSCR3 | 3.055 | 0.022 | 0.002 |
| AP4B1-AS1 | 3.047 | 0.023 | 0.002 |
| RP11-589M4.4 | 2.911 | 0.027 | 0.002 |
| CTD-2006C1.10 | -2.905 | 0.027 | 0.002 |
| ERMP1 | -2.903 | 0.027 | 0.002 |
| LPXN | 2.879 | 0.028 | 0.002 |
| NKILA | -2.879 | 0.028 | 0.002 |
| BAIAP2L2 | 2.835 | 0.030 | 0.002 |
| RBM26-AS1 | -2.799 | 0.031 | 0.002 |
| HLA-DOB | 2.747 | 0.033 | 0.002 |
| MPPE1 | -2.733 | 0.034 | 0.002 |
| PRSS27 | -2.690 | 0.036 | 0.002 |
| C3orf35 | -2.679 | 0.037 | 0.002 |
| ZDHHC9 | -2.650 | 0.038 | 0.002 |
| ZBTB34 | -2.647 | 0.038 | 0.002 |
| SNX3 | -2.607 | 0.040 | 0.002 |
| PGD | 2.599 | 0.041 | 0.002 |
| PYCR1 | 2.566 | 0.043 | 0.002 |
| C4B | -3.143 | 0.020 | 0.002 |
| RP11-665C16.9 | 2.847 | 0.029 | 0.002 |
| ZNF513 | 2.847 | 0.029 | 0.002 |
| PELI1 | -2.823 | 0.030 | 0.002 |
| SPHK1 | 2.805 | 0.031 | 0.002 |
| HACE1 | -2.772 | 0.032 | 0.002 |
| RFK | -2.673 | 0.037 | 0.002 |
| ORAI3 | -2.658 | 0.038 | 0.002 |
| NATD1 | -2.594 | 0.041 | 0.002 |
| RP11-961A15.3 | 2.567 | 0.043 | 0.002 |
| MIR612 | 2.565 | 0.043 | 0.002 |
| ZC3H12B | -2.558 | 0.043 | 0.002 |
| TRPA1 | -2.521 | 0.045 | 0.002 |
| CALM3 | 3.322 | 0.016 | 0.002 |
| SLC25A1 | 3.294 | 0.017 | 0.002 |
| RCBTB2 | -3.281 | 0.017 | 0.002 |
| ZNF177 | -3.043 | 0.023 | 0.002 |
| ARHGAP44 | -2.951 | 0.026 | 0.002 |
| ITGA1 | -2.937 | 0.026 | 0.002 |
| RPL26P30 | 2.757 | 0.033 | 0.002 |
| POLR2E | 2.731 | 0.034 | 0.002 |
| SH3BP5 | 2.705 | 0.035 | 0.002 |
| NOB1 | 2.655 | 0.038 | 0.002 |
| RP11-810M2.2 | -2.609 | 0.040 | 0.002 |
| CDCA7L | 2.583 | 0.042 | 0.002 |
| RP11-360F5.3 | -2.566 | 0.043 | 0.002 |

| | | | |
|---|---|---|---|
| NPTN | -2.550 | 0.043 | 0.002 |
| AC092574.1 | -3.121 | 0.021 | 0.002 |
| TBX18 | -3.052 | 0.022 | 0.002 |
| NAB2 | 2.949 | 0.026 | 0.002 |
| C8orf31 | -2.923 | 0.027 | 0.002 |
| DARS-AS1 | -2.909 | 0.027 | 0.002 |
| RP11-303E16.3 | 2.900 | 0.027 | 0.002 |
| NANOS1 | -2.840 | 0.030 | 0.002 |
| RNF122 | -2.817 | 0.030 | 0.002 |
| RP11-677I18.3 | 2.812 | 0.031 | 0.002 |
| ADAMTSL4-AS1 | -2.694 | 0.036 | 0.002 |
| NAA10 | 2.670 | 0.037 | 0.002 |
| BAD | -2.658 | 0.038 | 0.002 |
| GOLGA8K | -3.517 | 0.013 | 0.003 |
| TBC1D3P1-DHX40P1 | -3.252 | 0.017 | 0.003 |
| LAP3 | 2.944 | 0.026 | 0.003 |
| RB1CC1 | -2.883 | 0.028 | 0.003 |
| TMCC2 | -2.873 | 0.028 | 0.003 |
| CDYL | -2.754 | 0.033 | 0.003 |
| AL365273.1 | -2.750 | 0.033 | 0.003 |
| LYSMD1 | -2.659 | 0.038 | 0.003 |
| CD68 | 2.655 | 0.038 | 0.003 |
| KCNN1 | 2.594 | 0.041 | 0.003 |
| AC034220.3 | -2.505 | 0.046 | 0.003 |
| PDIA3 | -2.503 | 0.046 | 0.003 |
| AC108463.2 | -2.492 | 0.047 | 0.003 |
| SPRED1 | -3.444 | 0.014 | 0.003 |
| SMG6 | -2.950 | 0.026 | 0.003 |
| RP11-548H18.2 | -2.856 | 0.029 | 0.003 |
| LINC01341 | -2.806 | 0.031 | 0.003 |
| XXbac-BPG157A10.21 | -2.677 | 0.037 | 0.003 |
| L1CAM | 2.670 | 0.037 | 0.003 |
| ADGRL4 | -2.592 | 0.041 | 0.003 |
| ARMCX5 | -2.584 | 0.042 | 0.003 |
| HPS1 | 2.582 | 0.042 | 0.003 |
| MIR1287 | 2.582 | 0.042 | 0.003 |
| TET2 | -2.578 | 0.042 | 0.003 |
| KLHL5 | -2.566 | 0.043 | 0.003 |
| LINC00324 | -3.074 | 0.022 | 0.003 |
| ZNF433 | -2.905 | 0.027 | 0.003 |
| CTC-444N24.7 | -2.865 | 0.029 | 0.003 |
| NR1I3 | -2.816 | 0.031 | 0.003 |
| RALBP1 | -2.808 | 0.031 | 0.003 |
| FN3K | -2.767 | 0.033 | 0.003 |
| RP11-429G19.3 | -2.750 | 0.033 | 0.003 |
| MCL1 | -2.694 | 0.036 | 0.003 |
| GPRASP2 | -2.584 | 0.042 | 0.003 |
| MYADML2 | 2.566 | 0.043 | 0.003 |
| NEAT1_3 | 2.565 | 0.043 | 0.003 |

| | | | |
|---|---|---|---|
| ZNF829 | -2.554 | 0.043 | 0.003 |
| RALGAPA2 | -2.523 | 0.045 | 0.003 |
| RP11-420L9.2 | -3.380 | 0.015 | 0.003 |
| TMEM256 | 3.055 | 0.022 | 0.003 |
| RP11-303E16.9 | 2.900 | 0.027 | 0.003 |
| SRSF4 | 2.879 | 0.028 | 0.003 |
| SWT1 | -2.838 | 0.030 | 0.003 |
| S100A13 | 2.759 | 0.033 | 0.003 |
| C10orf131 | -2.750 | 0.033 | 0.003 |
| MIR6832 | 2.698 | 0.036 | 0.003 |
| PUM3 | 2.575 | 0.042 | 0.003 |
| FAM168A | -2.552 | 0.043 | 0.003 |
| RASSF7 | -2.533 | 0.044 | 0.003 |
| NIPSNAP3B | -2.519 | 0.045 | 0.003 |
| RTBDN | -2.492 | 0.047 | 0.003 |
| RP11-420L9.5 | -3.380 | 0.015 | 0.003 |
| CNNM2 | -3.210 | 0.018 | 0.003 |
| PLBD2 | -2.916 | 0.027 | 0.003 |
| ATG16L1 | -2.858 | 0.029 | 0.003 |
| STT3B | -2.850 | 0.029 | 0.003 |
| CPOX | -2.839 | 0.030 | 0.003 |
| C5orf15 | -2.824 | 0.030 | 0.003 |
| ING1 | -2.815 | 0.031 | 0.003 |
| SYT3 | 2.778 | 0.032 | 0.003 |
| CEP162 | -2.721 | 0.035 | 0.003 |
| RAP1GDS1 | -2.651 | 0.038 | 0.003 |
| RAF1 | 2.571 | 0.042 | 0.003 |
| NEAT1_1 | 2.565 | 0.043 | 0.003 |
| PMS2P2 | -2.528 | 0.045 | 0.003 |
| MIR6756 | -2.495 | 0.047 | 0.003 |
| RP11-524F11.1 | 2.464 | 0.049 | 0.003 |
| TBC1D3P1 | -3.252 | 0.017 | 0.003 |
| ZNF559-ZNF177 | -3.043 | 0.023 | 0.003 |
| KLHDC9 | -2.873 | 0.028 | 0.003 |
| SMCHD1 | -2.747 | 0.033 | 0.003 |
| POLR2L | 2.668 | 0.037 | 0.003 |
| MAP2K4 | -2.651 | 0.038 | 0.003 |
| KPNA6 | 2.642 | 0.038 | 0.003 |
| AC123768.1 | -2.634 | 0.039 | 0.003 |
| RPSAP54 | 2.570 | 0.042 | 0.003 |
| NEAT1_2 | 2.565 | 0.043 | 0.003 |
| RP11-254I22.1 | 2.521 | 0.045 | 0.003 |
| RCOR2 | -3.032 | 0.023 | 0.003 |
| UNC119 | -2.975 | 0.025 | 0.003 |
| RP11-112L6.3 | -2.840 | 0.030 | 0.003 |
| RP11-968O1.5 | -2.836 | 0.030 | 0.003 |
| RP11-248J23.7 | -2.750 | 0.033 | 0.003 |
| RP13-1032I1.7 | 2.675 | 0.037 | 0.003 |
| KBTBD2 | -2.662 | 0.037 | 0.003 |

| | | | |
|---|---|---|---|
| SENP3-EIF4A1 | 2.655 | 0.038 | 0.003 |
| PYROXD2 | 2.582 | 0.042 | 0.003 |
| ABI2 | 2.532 | 0.045 | 0.003 |
| CTD-3184A7.4 | 2.468 | 0.049 | 0.003 |
| PIGS | -2.461 | 0.049 | 0.003 |
| ZNF559 | -3.043 | 0.023 | 0.003 |
| NBPF12 | -3.028 | 0.023 | 0.003 |
| GRAMD4P7 | 2.988 | 0.024 | 0.003 |
| GCC2-AS1 | -2.889 | 0.028 | 0.003 |
| GNAI2 | 2.835 | 0.030 | 0.003 |
| B3GNT4 | -2.829 | 0.030 | 0.003 |
| MFHAS1 | -2.796 | 0.031 | 0.003 |
| RP1-178F15.4 | 2.759 | 0.033 | 0.003 |
| SNORD10 | 2.655 | 0.038 | 0.003 |
| RP11-690I21.1 | 2.572 | 0.042 | 0.003 |
| CES4A | 2.531 | 0.045 | 0.003 |
| CUL1 | -2.494 | 0.047 | 0.003 |
| FUT7 | 2.472 | 0.048 | 0.003 |
| RP11-500C11.3 | 3.089 | 0.021 | 0.003 |
| CCL2 | 3.021 | 0.023 | 0.003 |
| TWF1P1 | 2.918 | 0.027 | 0.003 |
| WDPCP | -2.702 | 0.035 | 0.003 |
| PSAP | -2.681 | 0.036 | 0.003 |
| ZNF610 | -2.672 | 0.037 | 0.003 |
| SENP3 | 2.655 | 0.038 | 0.003 |
| KRT80 | -2.617 | 0.040 | 0.003 |
| CYB5D2 | -2.609 | 0.040 | 0.003 |
| TRAP1 | 2.576 | 0.042 | 0.003 |
| LINC00894 | 3.270 | 0.017 | 0.003 |
| DTD1 | 3.001 | 0.024 | 0.003 |
| PARD3 | -2.936 | 0.026 | 0.003 |
| ACTG1 | 2.818 | 0.030 | 0.003 |
| RP1-178F15.5 | 2.759 | 0.033 | 0.003 |
| MSANTD3-TMEFF1 | 2.737 | 0.034 | 0.003 |
| DOCK10 | 2.716 | 0.035 | 0.003 |
| SGPL1 | -2.666 | 0.037 | 0.003 |
| EIF3I | 2.642 | 0.038 | 0.003 |
| RALGPS1 | -2.604 | 0.040 | 0.003 |
| S100A6 | 2.585 | 0.042 | 0.003 |
| CORO1C | 2.566 | 0.043 | 0.003 |
| EIF5A | 2.561 | 0.043 | 0.003 |
| DLL1 | 2.559 | 0.043 | 0.003 |
| GREM2 | 2.544 | 0.044 | 0.003 |
| DPY19L2 | -2.517 | 0.045 | 0.003 |
| U2 | 2.499 | 0.047 | 0.003 |
| MCAM | -2.495 | 0.047 | 0.003 |
| PPP2R3A | 2.450 | 0.050 | 0.003 |
| SCRN3 | -2.727 | 0.034 | 0.004 |
| STK38L | -2.696 | 0.036 | 0.004 |

| | | | |
|---|---|---|---|
| ARHGAP4 | 2.670 | 0.037 | 0.004 |
| SNORA67 | 2.655 | 0.038 | 0.004 |
| GOLGA8N | -2.634 | 0.039 | 0.004 |
| TUG1 | -2.629 | 0.039 | 0.004 |
| CCDC124 | 2.594 | 0.041 | 0.004 |
| VLDLR | -2.566 | 0.043 | 0.004 |
| RAB11FIP1 | -2.546 | 0.044 | 0.004 |
| TAPT1 | -2.543 | 0.044 | 0.004 |
| PDHX | -2.537 | 0.044 | 0.004 |
| LINC01358 | 2.934 | 0.026 | 0.004 |
| SHANK1 | 2.778 | 0.032 | 0.004 |
| RP11-12A20.6 | -2.776 | 0.032 | 0.004 |
| PHTF2 | -2.751 | 0.033 | 0.004 |
| NREP | -2.713 | 0.035 | 0.004 |
| PIGA | -2.667 | 0.037 | 0.004 |
| SNORA48 | 2.655 | 0.038 | 0.004 |
| TXLNA | 2.642 | 0.038 | 0.004 |
| COL5A3 | -2.638 | 0.039 | 0.004 |
| NEAT1 | 2.565 | 0.043 | 0.004 |
| SCOC-AS1 | -2.523 | 0.045 | 0.004 |
| AC020934.1 | -2.492 | 0.047 | 0.004 |
| NPDC1 | 2.472 | 0.048 | 0.004 |
| RP11-229P13.27 | 2.472 | 0.048 | 0.004 |
| PPIB | 2.463 | 0.049 | 0.004 |
| CNBP | 2.461 | 0.049 | 0.004 |
| CTD-2207L17.1 | -2.937 | 0.026 | 0.004 |
| PPIC | -2.675 | 0.037 | 0.004 |
| SAR1B | -2.660 | 0.038 | 0.004 |
| PLXNB2 | -2.642 | 0.038 | 0.004 |
| ABCC3 | 2.634 | 0.039 | 0.004 |
| SNORD6 | 2.610 | 0.040 | 0.004 |
| LSM7 | 2.550 | 0.043 | 0.004 |
| MIR21 | -2.536 | 0.044 | 0.004 |
| ASIC3 | -2.524 | 0.045 | 0.004 |
| RP5-1125A11.7 | -3.127 | 0.020 | 0.004 |
| LINC00493 | 3.001 | 0.024 | 0.004 |
| RP11-204C16.4 | 2.928 | 0.026 | 0.004 |
| UBA3 | -2.730 | 0.034 | 0.004 |
| ST3GAL1 | 2.685 | 0.036 | 0.004 |
| RP11-479O9.4 | -2.678 | 0.037 | 0.004 |
| CINP | 2.656 | 0.038 | 0.004 |
| AC044907.1 | 2.618 | 0.040 | 0.004 |
| NRROS | -2.589 | 0.041 | 0.004 |
| TRIM4 | -2.487 | 0.047 | 0.004 |
| GCLM | 2.472 | 0.048 | 0.004 |
| RP4-751H13.7 | -3.121 | 0.021 | 0.004 |
| SHANK2 | 3.033 | 0.023 | 0.004 |
| AC087884.1 | 2.870 | 0.028 | 0.004 |
| CNIH4 | -2.683 | 0.036 | 0.004 |

| | | | |
|---|---|---|---|
| TUG1_4 | -2.629 | 0.039 | 0.004 |
| RP11-539L10.3 | 2.626 | 0.039 | 0.004 |
| RP3-395M20.12 | 2.590 | 0.041 | 0.004 |
| RP11-461A8.4 | 2.576 | 0.042 | 0.004 |
| RP11-391L3.3 | -2.523 | 0.045 | 0.004 |
| AL022341.3 | 2.516 | 0.046 | 0.004 |
| TTC9C | 2.501 | 0.046 | 0.004 |
| RMND5A | -2.496 | 0.047 | 0.004 |
| TMED3 | 3.172 | 0.019 | 0.004 |
| B3GNT2 | -3.046 | 0.023 | 0.004 |
| TBC1D10C | -3.001 | 0.024 | 0.004 |
| ZNF878 | -2.905 | 0.027 | 0.004 |
| LMAN2L | -2.716 | 0.035 | 0.004 |
| SERPING1 | -2.685 | 0.036 | 0.004 |
| ACTL10 | 2.681 | 0.036 | 0.004 |
| DUSP4 | 2.617 | 0.040 | 0.004 |
| SCG2 | -2.589 | 0.041 | 0.004 |
| PIGX | -2.589 | 0.041 | 0.004 |
| ERBB2 | -2.571 | 0.042 | 0.004 |
| LSM12 | 2.521 | 0.045 | 0.004 |
| PIK3IP1 | -2.516 | 0.046 | 0.004 |
| ALDH9A1 | 2.486 | 0.047 | 0.004 |
| NDUFS6 | 2.474 | 0.048 | 0.004 |
| SCAPER | -2.458 | 0.049 | 0.004 |
| TDG | -2.702 | 0.035 | 0.004 |
| MTERF1 | -2.660 | 0.038 | 0.004 |
| SETBP1 | -2.511 | 0.046 | 0.004 |
| HOOK2 | -2.492 | 0.047 | 0.004 |
| ZNF527 | -2.487 | 0.047 | 0.004 |
| SIK2 | -2.642 | 0.038 | 0.004 |
| NUTM2E | -2.629 | 0.039 | 0.004 |
| TUG1_1 | -2.629 | 0.039 | 0.004 |
| ARPC4-TTLL3 | 2.602 | 0.041 | 0.004 |
| AC011043.1 | -2.550 | 0.044 | 0.004 |
| C6orf132 | -2.484 | 0.048 | 0.004 |
| EPB41L5 | -2.464 | 0.049 | 0.004 |
| S100A16 | 3.514 | 0.013 | 0.004 |
| DLX1 | 3.251 | 0.017 | 0.004 |
| EXOSC4 | 2.840 | 0.030 | 0.004 |
| CLDND1 | -2.839 | 0.030 | 0.004 |
| CTNNA1 | -2.586 | 0.041 | 0.004 |
| ZNF667 | -2.585 | 0.041 | 0.004 |
| AC015987.1 | -2.525 | 0.045 | 0.004 |
| NSRP1 | 2.521 | 0.045 | 0.004 |
| SCO2 | 3.133 | 0.020 | 0.004 |
| BBS4 | -3.037 | 0.023 | 0.004 |
| BMP1 | -2.885 | 0.028 | 0.004 |
| CDKL1 | 2.774 | 0.032 | 0.004 |
| GRAMD1A | -2.591 | 0.041 | 0.004 |

| | | | |
|---|---|---|---|
| RNF135 | -2.525 | 0.045 | 0.004 |
| MBNL1-AS1 | 2.510 | 0.046 | 0.004 |
| OR2AE1 | -2.487 | 0.047 | 0.004 |
| TYMP | 3.133 | 0.020 | 0.005 |
| PLCL1 | -2.907 | 0.027 | 0.005 |
| MIR6802 | -2.844 | 0.029 | 0.005 |
| GDNF | -2.688 | 0.036 | 0.005 |
| TUG1_3 | -2.629 | 0.039 | 0.005 |
| RP11-867G23.12 | 2.592 | 0.041 | 0.005 |
| FAM189B | 2.564 | 0.043 | 0.005 |
| FER | -2.564 | 0.043 | 0.005 |
| VMP1 | -2.536 | 0.044 | 0.005 |
| MED8 | 2.462 | 0.049 | 0.005 |
| PALM | -2.985 | 0.024 | 0.005 |
| CDK10 | 2.645 | 0.038 | 0.005 |
| MAP9 | -2.618 | 0.040 | 0.005 |
| KLF7 | -2.605 | 0.040 | 0.005 |
| MEX3B | -2.597 | 0.041 | 0.005 |
| SMIM12 | 2.573 | 0.042 | 0.005 |
| TRAPPC12-AS1 | -2.546 | 0.044 | 0.005 |
| GANC | -2.536 | 0.044 | 0.005 |
| FGF7 | 2.532 | 0.045 | 0.005 |
| AC120045.1 | -2.520 | 0.045 | 0.005 |
| WDR19 | -2.474 | 0.048 | 0.005 |
| KIAA1024 | 3.172 | 0.019 | 0.005 |
| AC139149.1 | 2.818 | 0.030 | 0.005 |
| CDC27P1 | 2.809 | 0.031 | 0.005 |
| MIA2 | -2.666 | 0.037 | 0.005 |
| RP11-407N17.3 | -2.666 | 0.037 | 0.005 |
| RP11-186B7.4 | 2.655 | 0.038 | 0.005 |
| TUG1_2 | -2.629 | 0.039 | 0.005 |
| ZNF793-AS1 | -2.621 | 0.040 | 0.005 |
| TTLL3 | 2.602 | 0.041 | 0.005 |
| SPATA1 | 2.697 | 0.036 | 0.005 |
| EIF4A1 | 2.655 | 0.038 | 0.005 |
| LAGE3 | 2.614 | 0.040 | 0.005 |
| HSF2BP | -2.601 | 0.041 | 0.005 |
| JUNB | -2.563 | 0.043 | 0.005 |
| METTL25 | -2.539 | 0.044 | 0.005 |
| IKZF4 | -2.535 | 0.044 | 0.005 |
| IFNGR1 | -2.531 | 0.045 | 0.005 |
| RP11-848P1.2 | -2.525 | 0.045 | 0.005 |
| LINC01137 | -2.483 | 0.048 | 0.005 |
| RP11-541N10.3 | -2.468 | 0.049 | 0.005 |
| CTC-463A16.1 | -2.457 | 0.049 | 0.005 |
| MEIS2 | -2.673 | 0.037 | 0.005 |
| ARPC4 | 2.602 | 0.041 | 0.005 |
| GLI4 | -2.595 | 0.041 | 0.005 |
| TAF11 | -2.558 | 0.043 | 0.005 |

| | | | |
|---|---|---|---|
| CAPN3 | -2.536 | 0.044 | 0.005 |
| MIR656 | 2.492 | 0.047 | 0.005 |
| TBC1D12 | -2.488 | 0.047 | 0.005 |
| FAM171A1 | -3.158 | 0.020 | 0.005 |
| ZNF652 | -2.975 | 0.025 | 0.005 |
| RP11-425D10.10 | -2.945 | 0.026 | 0.005 |
| CTD-2530N21.5 | -2.885 | 0.028 | 0.005 |
| ZFP41 | -2.595 | 0.041 | 0.005 |
| SNCG | 2.537 | 0.044 | 0.005 |
| HSD17B1 | -2.489 | 0.047 | 0.005 |
| MLH3 | -3.161 | 0.020 | 0.005 |
| USP25 | -2.835 | 0.030 | 0.005 |
| SNORA23 | 2.649 | 0.038 | 0.005 |
| RIMS3 | -2.624 | 0.039 | 0.005 |
| PPARGC1B | 2.549 | 0.044 | 0.005 |
| BNIP3P5 | -2.536 | 0.044 | 0.005 |
| RP11-164J13.1 | -2.536 | 0.044 | 0.005 |
| MIR423 | 2.521 | 0.045 | 0.005 |
| ZNF425 | -2.471 | 0.048 | 0.005 |
| NCBP2-AS1 | 2.469 | 0.049 | 0.005 |
| NDUFA6-AS1 | -2.457 | 0.049 | 0.005 |
| MIR6847 | 2.840 | 0.030 | 0.005 |
| CASP9 | -2.665 | 0.037 | 0.005 |
| RP11-635N19.1 | -2.645 | 0.038 | 0.005 |
| SCN1B | -2.591 | 0.041 | 0.005 |
| ARMC5 | -2.545 | 0.044 | 0.005 |
| MYCL | -2.505 | 0.046 | 0.005 |
| TXNDC16 | -2.466 | 0.049 | 0.005 |
| USP2-AS1 | -2.605 | 0.040 | 0.005 |
| TMEM67 | -2.594 | 0.041 | 0.005 |
| AL162151.3 | 2.588 | 0.041 | 0.005 |
| RP4-798A10.2 | -2.492 | 0.047 | 0.005 |
| CTD-2517M22.14 | -2.491 | 0.047 | 0.005 |
| AC007750.5 | -3.027 | 0.023 | 0.005 |
| TMEFF1 | 2.737 | 0.034 | 0.005 |
| MMP24 | -2.684 | 0.036 | 0.005 |
| IFT74 | -2.600 | 0.041 | 0.005 |
| RP13-582O9.6 | -2.595 | 0.041 | 0.005 |
| OSCAR | -2.524 | 0.045 | 0.005 |
| RP4-669P10.19 | -2.457 | 0.049 | 0.005 |
| ODF3B | 3.133 | 0.020 | 0.006 |
| ZNF345 | -2.820 | 0.030 | 0.006 |
| SCAMP3 | 2.564 | 0.043 | 0.006 |
| TAS1R3 | -2.561 | 0.043 | 0.006 |
| CTSB | 2.488 | 0.047 | 0.006 |
| RP4-622L5.2 | 2.642 | 0.038 | 0.006 |
| RP11-166D19.1 | 2.600 | 0.041 | 0.006 |
| CTD-2527I21.9 | -2.591 | 0.041 | 0.006 |
| RP11-1149O23.3 | 2.581 | 0.042 | 0.006 |

| | | | |
|---|---|---|---|
| TYW5 | -2.557 | 0.043 | 0.006 |
| AC010894.3 | -2.527 | 0.045 | 0.006 |
| GOLGA8J | -2.520 | 0.045 | 0.006 |
| MAU2 | -2.507 | 0.046 | 0.006 |
| BCL11A | -2.480 | 0.048 | 0.006 |
| ADAMTS4 | 2.447 | 0.050 | 0.006 |
| CHMP7 | 2.581 | 0.042 | 0.006 |
| RNF32 | -2.510 | 0.046 | 0.006 |
| 8-Mar | -2.471 | 0.048 | 0.006 |
| LA16c-360H6.1 | -2.772 | 0.032 | 0.006 |
| MSANTD3 | 2.737 | 0.034 | 0.006 |
| SLC9A5 | 2.570 | 0.042 | 0.006 |
| RP4-669P10.20 | -2.609 | 0.040 | 0.006 |
| PSKH1 | -2.592 | 0.041 | 0.006 |
| CYSRT1 | -2.957 | 0.025 | 0.006 |
| BLID | 2.600 | 0.041 | 0.006 |
| RPL34P18 | 2.583 | 0.042 | 0.006 |
| ZNF614 | -2.510 | 0.046 | 0.006 |
| NOXA1 | -2.477 | 0.048 | 0.006 |
| QRICH2 | -2.456 | 0.049 | 0.006 |
| MIR100HG | 2.600 | 0.041 | 0.006 |
| CCDC121 | -2.519 | 0.045 | 0.006 |
| ZRANB2-AS2 | -2.515 | 0.046 | 0.006 |
| KIAA1143 | -2.495 | 0.047 | 0.006 |
| YPEL1 | -2.623 | 0.039 | 0.006 |
| HSD17B1P1 | -2.489 | 0.047 | 0.006 |
| PABPC1 | 2.454 | 0.050 | 0.006 |
| FAM3C | 2.709 | 0.035 | 0.006 |
| SNAPC1 | -2.706 | 0.035 | 0.006 |
| MAP3K8 | -2.469 | 0.048 | 0.006 |
| PARP2 | -2.731 | 0.034 | 0.006 |
| RUSC2 | -2.536 | 0.044 | 0.006 |
| RP11-192H23.8 | -2.534 | 0.044 | 0.006 |
| CMIP | -2.523 | 0.045 | 0.006 |
| LTBP4 | -2.479 | 0.048 | 0.006 |
| MIR7705 | 2.454 | 0.050 | 0.006 |
| CERS6 | -2.942 | 0.026 | 0.007 |
| CTAGE5 | -2.666 | 0.037 | 0.007 |
| DRAM2 | -2.591 | 0.041 | 0.007 |
| RABL3 | -2.587 | 0.041 | 0.007 |
| AP2S1 | 2.504 | 0.046 | 0.007 |
| RP11-259P20.1 | -2.498 | 0.047 | 0.007 |
| GPAA1 | 2.840 | 0.030 | 0.007 |
| AC005795.1 | 2.682 | 0.036 | 0.007 |
| NSMCE1 | 2.652 | 0.038 | 0.007 |
| HMGCL | -2.607 | 0.040 | 0.007 |
| SYDE2 | -2.547 | 0.044 | 0.007 |
| MANBA | -2.535 | 0.044 | 0.007 |
| B3GALNT2 | -2.477 | 0.048 | 0.007 |

| | | | |
|---|---|---|---|
| P2RX7 | -3.154 | 0.020 | 0.007 |
| CTC-277H1.7 | 2.570 | 0.042 | 0.007 |
| EVA1C | 2.541 | 0.044 | 0.007 |
| NDUFAF2 | 2.516 | 0.046 | 0.007 |
| RP11-50C13.1 | -2.478 | 0.048 | 0.007 |
| HIF1A | -2.706 | 0.035 | 0.007 |
| RP11-57H14.2 | 2.467 | 0.049 | 0.007 |
| LRRC17 | -2.876 | 0.028 | 0.007 |
| RP11-618G20.1 | -2.706 | 0.035 | 0.007 |
| CHST15 | 2.604 | 0.040 | 0.007 |
| TMEM39A | 2.503 | 0.046 | 0.007 |
| PTMS | 2.481 | 0.048 | 0.007 |
| C5orf42 | -2.477 | 0.048 | 0.007 |
| CHADL | -2.470 | 0.048 | 0.007 |
| BNC1 | 2.683 | 0.036 | 0.007 |
| SLC12A6 | -3.163 | 0.019 | 0.007 |
| LINC00933 | -2.550 | 0.043 | 0.007 |
| PTCD1 | 2.455 | 0.049 | 0.007 |
| CEP104 | 2.722 | 0.035 | 0.007 |
| GNAO1 | -2.548 | 0.044 | 0.007 |
| ZNF507 | -2.504 | 0.046 | 0.007 |
| LINC00853 | -2.485 | 0.047 | 0.007 |
| CCDC43 | 2.592 | 0.041 | 0.008 |
| GPLD1 | -2.565 | 0.043 | 0.008 |
| DNASE2 | -2.525 | 0.045 | 0.008 |
| FGFR1 | 2.999 | 0.024 | 0.008 |
| C3orf70 | -2.495 | 0.047 | 0.008 |
| KLHL42 | -2.634 | 0.039 | 0.008 |
| MBIP | -2.556 | 0.043 | 0.008 |
| RP11-686D22.7 | -2.531 | 0.045 | 0.008 |
| FAM26E | -2.498 | 0.047 | 0.008 |
| AP001189.4 | -2.597 | 0.041 | 0.008 |
| SKAP2 | -2.543 | 0.044 | 0.008 |
| RP11-686D22.8 | -2.531 | 0.045 | 0.008 |
| DICER1-AS1 | -2.481 | 0.048 | 0.008 |
| AL450992.2 | -2.450 | 0.050 | 0.008 |
| ARL6 | -2.872 | 0.028 | 0.008 |
| EEF1E1 | 2.815 | 0.031 | 0.008 |
| KLF3 | -2.584 | 0.042 | 0.008 |
| MST1P2 | -2.553 | 0.043 | 0.008 |
| SUCLG2 | 2.531 | 0.045 | 0.008 |
| SDC4 | 2.513 | 0.046 | 0.008 |
| RP11-634H22.1 | -2.507 | 0.046 | 0.008 |
| RP11-1021N1.1 | -2.476 | 0.048 | 0.008 |
| ATP5J2-PTCD1 | 2.455 | 0.049 | 0.008 |
| AC120045.2 | -2.454 | 0.050 | 0.008 |
| EIF4BP3 | 2.512 | 0.046 | 0.008 |
| RPPH1 | 2.506 | 0.046 | 0.008 |
| ZNF354C | -2.490 | 0.047 | 0.008 |

| | | | |
|---|---|---|---|
| ASB14 | -2.457 | 0.049 | 0.008 |
| RP11-686D22.3 | -2.531 | 0.045 | 0.008 |
| HDAC6 | -2.561 | 0.043 | 0.009 |
| OTUD1 | -2.560 | 0.043 | 0.009 |
| WSB1 | -2.614 | 0.040 | 0.009 |
| ATP5J2 | 2.455 | 0.049 | 0.009 |
| RP11-540D14.8 | -2.575 | 0.042 | 0.009 |
| SLFN12 | -2.531 | 0.045 | 0.009 |
| SCRG1 | -3.073 | 0.022 | 0.009 |
| SPATC1L | 2.947 | 0.026 | 0.009 |
| PPIAP31 | 2.498 | 0.047 | 0.009 |
| BLOC1S5-TXNDC5 | 2.815 | 0.031 | 0.009 |
| EFNA4 | -2.575 | 0.042 | 0.009 |
| RPSAP3 | 2.640 | 0.039 | 0.009 |
| LINC01473 | -2.706 | 0.035 | 0.009 |
| RP11-488C13.1 | 2.569 | 0.042 | 0.009 |
| NIP7 | 2.499 | 0.047 | 0.009 |
| LYSMD2 | 2.487 | 0.047 | 0.009 |
| POLR1B | 2.485 | 0.047 | 0.009 |
| FTCD | 2.947 | 0.026 | 0.009 |
| EFNA3 | -2.575 | 0.042 | 0.009 |
| GOLGA8T | -2.454 | 0.050 | 0.009 |
| BLOC1S5 | 2.815 | 0.031 | 0.010 |
| MFSD8 | -2.686 | 0.036 | 0.010 |
| RUNDC3A | -2.488 | 0.047 | 0.010 |
| RPL23AP42 | 3.110 | 0.021 | 0.010 |
| PRORSD1P | -2.479 | 0.048 | 0.010 |
| TXNDC5 | 2.815 | 0.031 | 0.010 |
| AC012358.8 | -2.479 | 0.048 | 0.010 |
| SETSIP | 2.461 | 0.049 | 0.010 |
| VPS41 | -2.481 | 0.048 | 0.010 |
| C16orf45 | -2.476 | 0.048 | 0.010 |
| EEF1E1-BLOC1S5 | 2.815 | 0.031 | 0.010 |
| RP11-395G23.3 | -2.515 | 0.046 | 0.010 |
| OXR1 | -2.515 | 0.046 | 0.011 |
| MAP6 | -2.718 | 0.035 | 0.011 |
| AKIRIN1 | -2.516 | 0.046 | 0.011 |
| MPV17L | -2.476 | 0.048 | 0.011 |
| HS3ST3B1 | 2.776 | 0.032 | 0.011 |
| RPL24P4 | 2.643 | 0.038 | 0.011 |
| RP11-1000B6.5 | -2.608 | 0.040 | 0.011 |
| HOMER1 | -2.883 | 0.028 | 0.012 |
| STAC | -2.608 | 0.040 | 0.012 |
| A2M | -2.557 | 0.043 | 0.012 |
| MAPKAPK5-AS1 | -2.557 | 0.043 | 0.012 |
| AC006538.1 | -2.646 | 0.038 | 0.012 |
| KCNQ4 | -2.448 | 0.050 | 0.012 |
| ADARB1 | -2.592 | 0.041 | 0.012 |
| RP11-596C23.6 | 2.496 | 0.047 | 0.012 |

| | | | |
|---|---|---|---|
| RP11-566K11.2 | 2.609 | 0.040 | 0.013 |
| ABCG4 | -2.750 | 0.033 | 0.013 |
| NAIP | -2.447 | 0.050 | 0.013 |
| SGCA | -2.735 | 0.034 | 0.013 |
| TUBB3 | 2.609 | 0.040 | 0.013 |
| AL627171.1 | 2.496 | 0.047 | 0.013 |
| RP11-566K11.7 | 2.609 | 0.040 | 0.013 |
| KLHL21 | -2.455 | 0.049 | 0.013 |
| POPDC3 | 2.533 | 0.044 | 0.015 |
| TCF25 | 2.609 | 0.040 | 0.015 |
| RN7SL3 | 2.496 | 0.047 | 0.015 |
| TRIM14 | 2.663 | 0.037 | 0.015 |
| MC1R | 2.609 | 0.040 | 0.016 |
| RN7SL2 | 2.496 | 0.047 | 0.016 |
| PRSS35 | -2.546 | 0.044 | 0.016 |
| DYNLL1P1 | 2.517 | 0.045 | 0.017 |
| ZFP1 | -2.490 | 0.047 | 0.017 |
| TOR4A | 2.532 | 0.045 | 0.018 |
| SDAD1P1 | -2.514 | 0.046 | 0.018 |
| ESPNL | -2.960 | 0.025 | 0.019 |
| RPL4P4 | 2.448 | 0.050 | 0.020 |
| SOX30 | -2.596 | 0.041 | 0.020 |
| NELFB | 2.532 | 0.045 | 0.021 |
| IVNS1ABP | -2.603 | 0.041 | 0.021 |
| GALNT6 | 2.607 | 0.040 | 0.025 |
| CCDC181 | -2.693 | 0.036 | 0.028 |
| RP3-469D22.1 | -2.472 | 0.048 | 0.040 |

**Table S5. Genes associated with the right olfactory bulb volume in healthy controls.** Abbreviations: GLR, generalized linear regression.

| **Gene** | **t-value (GLR)** | **p-value (GLR)** | **p-value (permutation)** |
|---|---|---|---|
| GSKIP | -5.067 | 9.52E-05 | 0.00 |
| PPP1R35 | 4.454 | 3.49E-04 | 1.00E-04 |
| RNU6-1262P | -4.327 | 4.57E-04 | 1.00E-04 |
| C7orf73 | 3.801 | 1.43E-03 | 1.00E-04 |
| RPL12P38 | 4.544 | 2.87E-04 | 2.00E-04 |
| RP11-175I6.1 | 3.870 | 0.001 | 3.00E-04 |
| RP11-447L10.1 | -3.723 | 0.002 | 3.00E-04 |
| TM4SF19-TCTEX1D2 | -3.723 | 0.002 | 4.00E-04 |
| MESP1 | 3.650 | 0.002 | 4.00E-04 |
| SNORD41 | -3.505 | 0.003 | 4.00E-04 |
| TCTEX1D2 | -3.723 | 0.002 | 0.001 |
| RP11-475I24.1 | 3.719 | 0.002 | 0.001 |
| SNORD15A | 3.400 | 0.003 | 0.001 |
| MTA3 | 3.952 | 0.001 | 0.001 |
| PCYT1A | -3.723 | 0.002 | 0.001 |
| HILPDA | 3.517 | 0.003 | 0.001 |
| TM4SF19 | -3.723 | 0.002 | 0.001 |
| RP11-459F6.3 | -3.532 | 0.003 | 0.001 |
| TMUB1 | 3.394 | 0.003 | 0.001 |
| STK35 | -3.353 | 0.004 | 0.001 |
| MT-CYB | -3.099 | 0.007 | 0.001 |
| RP11-155G14.5 | 3.517 | 0.003 | 0.001 |
| RPS3 | 3.400 | 0.003 | 0.001 |
| ZNF592 | -3.727 | 0.002 | 0.001 |
| METTL2B | 3.517 | 0.003 | 0.001 |
| RP5-1050D4.5 | -3.401 | 0.003 | 0.001 |
| GREM1 | 3.560 | 0.002 | 0.001 |
| ZNF398 | -3.374 | 0.004 | 0.001 |
| RPS6 | 3.398 | 0.003 | 0.001 |
| NBPF20 | 3.350 | 0.004 | 0.001 |
| PMS2P11 | -3.287 | 0.004 | 0.001 |
| MAP3K3 | -3.569 | 0.002 | 0.002 |
| RP11-212P7.3 | 3.517 | 0.003 | 0.002 |
| ZNF845 | -3.366 | 0.004 | 0.002 |
| DTX2P1-UPK3BP1-PMS2P11 | -3.287 | 0.004 | 0.002 |
| ZNF224 | -3.549 | 0.002 | 0.002 |
| DST | -3.414 | 0.003 | 0.002 |
| RP11-266L9.8 | 3.325 | 0.004 | 0.002 |
| RP11-155G14.6 | 3.517 | 0.003 | 0.002 |
| RP11-173C1.1 | 3.453 | 0.003 | 0.002 |
| RP11-467H10.2 | -3.287 | 0.004 | 0.002 |
| RN7SKP203 | 3.282 | 0.004 | 0.002 |
| RN7SK | 3.288 | 0.004 | 0.002 |
| PEF1 | 3.270 | 0.005 | 0.002 |

| | | | |
|---|---|---|---|
| CLCN5 | -3.149 | 0.006 | 0.002 |
| AP5M1 | -3.105 | 0.006 | 0.002 |
| UPK3BP1 | -3.287 | 0.004 | 0.002 |
| RP11-272L13.3 | -3.490 | 0.003 | 0.002 |
| DTX2P1 | -3.287 | 0.004 | 0.002 |
| RBM24 | -3.161 | 0.006 | 0.002 |
| C10orf35 | 2.994 | 0.008 | 0.002 |
| TSC2 | -2.871 | 0.011 | 0.002 |
| PMS2P9 | -3.287 | 0.004 | 0.003 |
| NOMO3 | -3.180 | 0.005 | 0.003 |
| MIR3179-2 | -3.180 | 0.005 | 0.003 |
| MT-ATP8 | -3.048 | 0.007 | 0.003 |
| CCDC146 | -3.287 | 0.004 | 0.003 |
| PLEKHH3 | 3.234 | 0.005 | 0.003 |
| ISG20L2 | -2.757 | 0.013 | 0.003 |
| NBPF25P | 3.350 | 0.004 | 0.003 |
| AL353644.10 | 3.136 | 0.006 | 0.003 |
| AL353644.11 | 3.136 | 0.006 | 0.003 |
| AL353644.3 | 3.136 | 0.006 | 0.003 |
| AL353644.4 | 3.136 | 0.006 | 0.003 |
| AL353644.1 | 3.136 | 0.006 | 0.003 |
| AL353644.9 | 3.136 | 0.006 | 0.003 |
| CTD-2031P19.5 | -3.120 | 0.006 | 0.003 |
| ZXDC | -3.094 | 0.007 | 0.003 |
| SOX15 | -3.154 | 0.006 | 0.003 |
| DCLRE1B | -3.136 | 0.006 | 0.003 |
| ZNF707 | -2.509 | 0.023 | 0.003 |
| AL353644.8 | 3.136 | 0.006 | 0.003 |
| RP11-146B14.1 | -3.039 | 0.007 | 0.003 |
| NAAA | -3.104 | 0.006 | 0.003 |
| MT-ND4L | -3.048 | 0.007 | 0.003 |
| RP11-390F4.3 | -3.039 | 0.007 | 0.003 |
| hsa-mir-3687-1 | 3.136 | 0.006 | 0.003 |
| WDR7 | -3.196 | 0.005 | 0.004 |
| CTD-3035K23.3 | -3.158 | 0.006 | 0.004 |
| RP11-757O6.6 | 3.002 | 0.008 | 0.004 |
| ZNF667-AS1 | 2.958 | 0.009 | 0.004 |
| RPS4XP17 | 3.037 | 0.007 | 0.004 |
| RHOBTB3 | 2.998 | 0.008 | 0.004 |
| MT-ATP6 | -3.048 | 0.007 | 0.004 |
| SNHG8 | 3.040 | 0.007 | 0.004 |
| PHOSPHO2 | 3.032 | 0.008 | 0.004 |
| KIAA1755 | -3.087 | 0.007 | 0.004 |
| KDM4C | -3.039 | 0.007 | 0.004 |
| KLHL23 | 3.032 | 0.008 | 0.004 |
| PLEKHM3 | -2.982 | 0.008 | 0.004 |
| RP5-858L17.1 | 2.860 | 0.011 | 0.004 |
| RP13-554M15.7 | -3.124 | 0.006 | 0.004 |
| MT-ND4 | -3.048 | 0.007 | 0.004 |

| | | | |
|---|---|---|---|
| FHDC1 | 3.027 | 0.008 | 0.004 |
| TRAPPC10 | -2.958 | 0.009 | 0.004 |
| AL592188.3 | 2.957 | 0.009 | 0.004 |
| NXF3 | 2.892 | 0.010 | 0.004 |
| MT-CO3 | -3.048 | 0.007 | 0.004 |
| RPSAP17 | 3.013 | 0.008 | 0.004 |
| RP11-219A15.1 | 2.974 | 0.009 | 0.004 |
| AL592188.8 | 2.957 | 0.009 | 0.004 |
| WDR7-OT1 | -3.196 | 0.005 | 0.004 |
| RP11-629O1.2 | 3.057 | 0.007 | 0.004 |
| AL513412.1 | -3.039 | 0.007 | 0.004 |
| RASA3 | -2.844 | 0.011 | 0.004 |
| SDAD1 | -3.104 | 0.006 | 0.004 |
| TMEM79 | -3.010 | 0.008 | 0.004 |
| AL592188.11 | 2.957 | 0.009 | 0.004 |
| AL592188.7 | 2.957 | 0.009 | 0.004 |
| RP11-361D15.2 | 2.935 | 0.009 | 0.004 |
| CEBPB | 2.883 | 0.010 | 0.004 |
| AL592188.6 | 2.957 | 0.009 | 0.004 |
| RBMS3-AS3 | -2.906 | 0.010 | 0.004 |
| CCIN | -2.970 | 0.009 | 0.005 |
| MLXIP | -2.939 | 0.009 | 0.005 |
| VPS53 | 3.037 | 0.007 | 0.005 |
| FBXL18 | -2.772 | 0.013 | 0.005 |
| TMSB4XP8 | 2.953 | 0.009 | 0.005 |
| HINFP | -3.023 | 0.008 | 0.005 |
| CCDC144A | 2.974 | 0.009 | 0.005 |
| CAB39L | -2.903 | 0.010 | 0.005 |
| RPS29 | 2.804 | 0.012 | 0.005 |
| USP32P1 | 2.974 | 0.009 | 0.005 |
| CRYBG3 | -2.851 | 0.011 | 0.005 |
| CTSA | -2.678 | 0.016 | 0.005 |
| SNORA24 | 3.040 | 0.007 | 0.005 |
| NOS2P4 | 2.974 | 0.009 | 0.005 |
| SMG6 | -2.905 | 0.010 | 0.005 |
| ZNF155 | -2.829 | 0.012 | 0.006 |
| HIST2H2BF | -2.855 | 0.011 | 0.006 |
| LONP2 | -2.832 | 0.012 | 0.006 |
| RP11-219A15.2 | 2.974 | 0.009 | 0.006 |
| PROCA1 | -2.835 | 0.011 | 0.006 |
| FTH1P16 | 2.806 | 0.012 | 0.006 |
| PRRC1 | -2.983 | 0.008 | 0.006 |
| RP11-219A15.4 | 2.974 | 0.009 | 0.006 |
| RNU4-32P | 2.740 | 0.014 | 0.006 |
| WIZ | -2.868 | 0.011 | 0.006 |
| FAM89A | 2.798 | 0.012 | 0.006 |
| ANKRD27 | 2.851 | 0.011 | 0.006 |
| RP11-130F10.1 | 2.662 | 0.016 | 0.006 |
| FASTK | 2.822 | 0.012 | 0.007 |

| | | | |
|---|---|---|---|
| MIR1182 | 2.798 | 0.012 | 0.007 |
| ZNF579 | 2.860 | 0.011 | 0.007 |
| MED1 | -2.857 | 0.011 | 0.007 |
| MMAB | 2.740 | 0.014 | 0.007 |
| PCCA | 2.726 | 0.014 | 0.007 |
| FOXD4 | -2.871 | 0.011 | 0.007 |
| RIBC1 | 2.745 | 0.014 | 0.007 |
| METTL16 | -2.741 | 0.014 | 0.007 |
| DENND2D | 2.687 | 0.016 | 0.007 |
| ZNF213 | -2.574 | 0.020 | 0.007 |
| RP11-148K1.12 | 2.822 | 0.012 | 0.007 |
| RP11-433J22.2 | 2.818 | 0.012 | 0.007 |
| PPCS | 2.786 | 0.013 | 0.007 |
| TBC1D20 | -2.745 | 0.014 | 0.007 |
| RPL7 | 2.740 | 0.014 | 0.007 |
| RALY | -2.767 | 0.013 | 0.008 |
| UACA | -2.739 | 0.014 | 0.008 |
| CACNA2D1 | -2.685 | 0.016 | 0.008 |
| KB-318B8.7 | -2.683 | 0.016 | 0.008 |
| RP9 | 2.637 | 0.017 | 0.008 |
| SPEN | -2.776 | 0.013 | 0.008 |
| DGCR2 | -2.673 | 0.016 | 0.008 |
| RPL32P29 | 2.804 | 0.012 | 0.008 |
| ASB16-AS1 | -2.776 | 0.013 | 0.008 |
| AARS | 2.720 | 0.015 | 0.008 |
| FAM227A | -2.710 | 0.015 | 0.008 |
| UBA52P6 | -2.568 | 0.020 | 0.008 |
| EXOSC6 | 2.720 | 0.015 | 0.008 |
| LONRF1 | 2.667 | 0.016 | 0.008 |
| RP9P | 2.637 | 0.017 | 0.008 |
| RPL23 | 2.629 | 0.018 | 0.008 |
| 44256 | -2.179 | 0.044 | 0.008 |
| MARK3 | 2.765 | 0.013 | 0.008 |
| ZNF3 | 2.679 | 0.016 | 0.008 |
| PDGFC | -2.669 | 0.016 | 0.008 |
| LMTK2 | -2.616 | 0.018 | 0.009 |
| PRR14L | -2.615 | 0.018 | 0.009 |
| CHM | -2.508 | 0.023 | 0.009 |
| MYL6 | 2.694 | 0.015 | 0.009 |
| RP11-177J6.1 | -2.693 | 0.015 | 0.009 |
| RPL24 | 2.688 | 0.016 | 0.009 |
| FAM109A | -2.554 | 0.021 | 0.009 |
| CCDC30 | 2.786 | 0.013 | 0.009 |
| AC005071.3 | -2.718 | 0.015 | 0.009 |
| SETX | -2.570 | 0.020 | 0.009 |
| DMTN | 2.784 | 0.013 | 0.009 |
| SNORD3C | 2.703 | 0.015 | 0.009 |
| PLOD3 | -2.629 | 0.018 | 0.009 |
| RP11-574K11.32 | 2.582 | 0.019 | 0.009 |

| | | | |
|---|---|---|---|
| SNORD3D | 2.703 | 0.015 | 0.009 |
| MFAP3 | -2.673 | 0.016 | 0.009 |
| RP11-160E2.6 | 2.703 | 0.015 | 0.009 |
| ASCC2 | -2.650 | 0.017 | 0.009 |
| CAMSAP1 | -2.707 | 0.015 | 0.009 |
| RP11-111F5.5 | 2.646 | 0.017 | 0.009 |
| FP236383.3 | 2.546 | 0.021 | 0.009 |
| GALNT10 | -2.673 | 0.016 | 0.009 |
| RP11-539L10.3 | -2.667 | 0.016 | 0.009 |
| C11orf65 | -2.758 | 0.013 | 0.010 |
| MYL6B | 2.694 | 0.015 | 0.010 |
| SLC45A4 | -2.670 | 0.016 | 0.010 |
| CH507-513H4.6 | 2.618 | 0.018 | 0.010 |
| ZNFX1 | -2.614 | 0.018 | 0.010 |
| DNAJB6 | 2.677 | 0.016 | 0.010 |
| ZNF79 | -2.706 | 0.015 | 0.010 |
| PTP4A2 | 2.647 | 0.017 | 0.010 |
| CH507-513H4.5 | 2.601 | 0.019 | 0.010 |
| CTD-2530H12.4 | 2.573 | 0.020 | 0.010 |
| FP236383.12 | 2.546 | 0.021 | 0.010 |
| SUMF2 | -2.630 | 0.018 | 0.010 |
| FP671120.7 | 2.546 | 0.021 | 0.010 |
| FP236383.9 | 2.546 | 0.021 | 0.010 |
| HIST2H2BD | -2.532 | 0.021 | 0.010 |
| RP11-57H14.2 | -2.623 | 0.018 | 0.010 |
| HDDC3 | 2.596 | 0.019 | 0.010 |
| SMG1P7 | 2.720 | 0.015 | 0.010 |
| RP11-90L1.8 | -2.708 | 0.015 | 0.010 |
| DGCR11 | -2.673 | 0.016 | 0.010 |
| ZNF736 | -2.560 | 0.020 | 0.010 |
| FP671120.4 | 2.546 | 0.021 | 0.010 |
| ARHGEF12 | -2.693 | 0.015 | 0.010 |
| SEC24B | -2.634 | 0.017 | 0.010 |
| ADD3 | -2.628 | 0.018 | 0.010 |
| SUSD2 | 2.677 | 0.016 | 0.011 |
| USP35 | -2.479 | 0.024 | 0.011 |
| AC008440.5 | -2.539 | 0.021 | 0.011 |
| NOMO2 | -2.512 | 0.022 | 0.011 |
| CHAD | -2.634 | 0.017 | 0.011 |
| ZNF879 | -2.602 | 0.019 | 0.011 |
| GPSM1 | -2.562 | 0.020 | 0.011 |
| CTD-2369P2.5 | 2.494 | 0.023 | 0.011 |
| IER3IP1 | 2.621 | 0.018 | 0.011 |
| IGIP | -2.597 | 0.019 | 0.011 |
| ADAMTS7P3 | 2.665 | 0.016 | 0.011 |
| CSPG4P11 | -2.571 | 0.020 | 0.011 |
| ZFAND5 | 2.511 | 0.022 | 0.011 |
| SNORA21 | 2.629 | 0.018 | 0.011 |
| PURA | -2.597 | 0.019 | 0.011 |

| | | | |
|---|---|---|---|
| ATP7A | -2.610 | 0.018 | 0.011 |
| FP236383.10 | 2.546 | 0.021 | 0.011 |
| FP236383.4 | 2.546 | 0.021 | 0.011 |
| RP5-1000K24.2 | -2.610 | 0.018 | 0.011 |
| CH507-513H4.1 | 2.546 | 0.021 | 0.011 |
| SLC25A20 | -2.541 | 0.021 | 0.011 |
| HSBP1L1 | 2.480 | 0.024 | 0.011 |
| ZBTB3 | -2.560 | 0.020 | 0.012 |
| POLB | 2.532 | 0.022 | 0.012 |
| RP11-865I6.2 | 2.521 | 0.022 | 0.012 |
| OSBPL11 | -2.584 | 0.019 | 0.012 |
| MOB2 | -2.580 | 0.019 | 0.012 |
| GALNT2 | -2.518 | 0.022 | 0.012 |
| ZNF134 | -2.487 | 0.024 | 0.012 |
| RP11-111M22.2 | -2.549 | 0.021 | 0.012 |
| KLF7-IT1 | -2.506 | 0.023 | 0.012 |
| COX5B | 2.633 | 0.017 | 0.012 |
| HDHD2 | 2.621 | 0.018 | 0.012 |
| BCL7B | 2.578 | 0.020 | 0.012 |
| C1orf189 | 2.421 | 0.027 | 0.012 |
| SIRT5 | 2.615 | 0.018 | 0.012 |
| RPL24P2 | 2.551 | 0.021 | 0.012 |
| RPS23 | 2.536 | 0.021 | 0.012 |
| RP11-574F21.2 | -2.470 | 0.024 | 0.012 |
| RAPH1 | -2.522 | 0.022 | 0.013 |
| RP11-49K24.6 | 2.621 | 0.018 | 0.013 |
| CTD-2622I13.3 | -2.501 | 0.023 | 0.013 |
| ZNF84 | -2.382 | 0.029 | 0.013 |
| MIR3179-3 | -2.512 | 0.022 | 0.013 |
| ZNF211 | -2.487 | 0.024 | 0.013 |
| HNRNPA1P7 | 2.435 | 0.026 | 0.013 |
| RP5-855D21.3 | -2.524 | 0.022 | 0.013 |
| MIR3179-4 | -2.512 | 0.022 | 0.013 |
| CH507-528H12.1 | 2.546 | 0.021 | 0.013 |
| SLC7A11-AS1 | 2.515 | 0.022 | 0.013 |
| FTLP3 | 2.490 | 0.023 | 0.013 |
| SLC38A10 | -2.499 | 0.023 | 0.013 |
| DICER1 | -2.356 | 0.031 | 0.013 |
| CTB-131B5.5 | -2.597 | 0.019 | 0.013 |
| AC007969.5 | 2.455 | 0.025 | 0.013 |
| LAMC1 | -2.516 | 0.022 | 0.014 |
| RPS16 | 2.400 | 0.028 | 0.014 |
| BCAT2 | 2.472 | 0.024 | 0.014 |
| STK32C | 2.467 | 0.025 | 0.014 |
| TPI1 | 2.541 | 0.021 | 0.014 |
| C1orf43 | 2.421 | 0.027 | 0.014 |
| ADD1 | -2.403 | 0.028 | 0.014 |
| RNU1-1 | 2.250 | 0.038 | 0.014 |
| SPTBN1 | -2.539 | 0.021 | 0.014 |

| | | | |
|---|---|---|---|
| RP11-298D21.3 | 2.458 | 0.025 | 0.014 |
| ZNF281 | 2.435 | 0.026 | 0.014 |
| PTPN1 | -2.523 | 0.022 | 0.014 |
| NECAB3 | 2.517 | 0.022 | 0.014 |
| RPL7P1 | 2.453 | 0.025 | 0.014 |
| RPSAP54 | 2.442 | 0.026 | 0.014 |
| CDC42BPB | -2.457 | 0.025 | 0.014 |
| TTC7B | -2.382 | 0.029 | 0.014 |
| AGAP1-IT1 | -2.304 | 0.034 | 0.014 |
| CCM2 | -2.531 | 0.022 | 0.014 |
| SPIRE1 | -2.460 | 0.025 | 0.014 |
| RP11-849H4.4 | -2.459 | 0.025 | 0.014 |
| ZNF705E | -2.461 | 0.025 | 0.015 |
| FAM157A | 2.462 | 0.025 | 0.015 |
| NPTX1 | -2.329 | 0.032 | 0.015 |
| CTD-3157E16.2 | 2.517 | 0.022 | 0.015 |
| RP11-343L5.2 | -2.510 | 0.022 | 0.015 |
| RPS25 | 2.484 | 0.024 | 0.015 |
| ZNF41 | -2.452 | 0.025 | 0.015 |
| CARD6 | -2.464 | 0.025 | 0.015 |
| PHKA2-AS1 | -2.540 | 0.021 | 0.015 |
| ALG1L9P | -2.461 | 0.025 | 0.015 |
| FP671120.6 | 2.464 | 0.025 | 0.015 |
| TFPI2 | -2.480 | 0.024 | 0.015 |
| PGBD4 | -2.456 | 0.025 | 0.015 |
| SLC27A5 | 2.392 | 0.029 | 0.015 |
| FKBP15 | -2.393 | 0.029 | 0.015 |
| TNFSF13B | -2.537 | 0.021 | 0.015 |
| SNORA14B | -2.485 | 0.024 | 0.015 |
| HELB | 2.441 | 0.026 | 0.015 |
| WBSCR27 | -2.474 | 0.024 | 0.015 |
| RP11-121P12.1 | 2.435 | 0.026 | 0.015 |
| CCNC | -2.433 | 0.026 | 0.015 |
| SMIM10L1 | -2.263 | 0.037 | 0.016 |
| PPIAP22 | 2.416 | 0.027 | 0.016 |
| RP11-547D13.1 | 2.398 | 0.028 | 0.016 |
| FAM71E2 | 2.359 | 0.031 | 0.016 |
| EEF1A1P13 | 2.451 | 0.025 | 0.016 |
| FSCN3 | 2.417 | 0.027 | 0.016 |
| IGF1R | -2.453 | 0.025 | 0.016 |
| MORF4 | 2.430 | 0.026 | 0.016 |
| TNFRSF1A | 2.388 | 0.029 | 0.016 |
| SLC9A1 | -2.451 | 0.025 | 0.016 |
| KRT8P33 | -2.448 | 0.026 | 0.016 |
| RP11-359P5.1 | -2.413 | 0.027 | 0.016 |
| MT-ND3 | -2.379 | 0.029 | 0.016 |
| CTD-2105E13.6 | 2.359 | 0.031 | 0.016 |
| BICD2 | -2.448 | 0.026 | 0.016 |
| HAUS7 | -2.333 | 0.032 | 0.016 |

| | | | |
|---|---|---|---|
| C6orf132 | 2.331 | 0.032 | 0.017 |
| RPSAP15 | 2.447 | 0.026 | 0.017 |
| ARF5 | 2.417 | 0.027 | 0.017 |
| RP11-195B17.1 | -2.400 | 0.028 | 0.017 |
| GPR107 | -2.453 | 0.025 | 0.017 |
| SNORA52 | 2.381 | 0.029 | 0.017 |
| TSN | 2.368 | 0.030 | 0.017 |
| LA16c-349E10.1 | -2.439 | 0.026 | 0.017 |
| DDT | -2.382 | 0.029 | 0.017 |
| SLC16A1-AS1 | 2.407 | 0.028 | 0.017 |
| SNORD83B | 2.404 | 0.028 | 0.017 |
| AP000351.3 | -2.382 | 0.029 | 0.017 |
| COX6B2 | 2.359 | 0.031 | 0.017 |
| RP11-753H16.5 | -2.441 | 0.026 | 0.017 |
| AC005944.2 | -2.434 | 0.026 | 0.017 |
| CIRBP-AS1 | -2.423 | 0.027 | 0.017 |
| ZBTB7A | 2.439 | 0.026 | 0.017 |
| HMGB1P1 | -2.387 | 0.029 | 0.017 |
| RPLP2 | 2.381 | 0.029 | 0.017 |
| RP11-115C10.1 | 2.380 | 0.029 | 0.017 |
| HMGN3-AS1 | 2.461 | 0.025 | 0.018 |
| RPL3 | 2.404 | 0.028 | 0.018 |
| COMMD7 | 2.383 | 0.029 | 0.018 |
| CDH6 | -2.395 | 0.028 | 0.018 |
| EIF4HP1 | 2.407 | 0.028 | 0.018 |
| SNORD43 | 2.404 | 0.028 | 0.018 |
| ATP9B | -2.367 | 0.030 | 0.018 |
| TGFBR2 | -2.373 | 0.030 | 0.018 |
| CARF | -2.401 | 0.028 | 0.018 |
| AF038458.4 | 2.368 | 0.030 | 0.018 |
| RP11-356J5.12 | 2.342 | 0.032 | 0.018 |
| CCDC39 | -2.382 | 0.029 | 0.018 |
| KIAA2026 | -2.323 | 0.033 | 0.018 |
| RSL24D1 | 2.398 | 0.028 | 0.018 |
| AP002495.1 | -2.386 | 0.029 | 0.018 |
| ZNF431 | -2.295 | 0.035 | 0.018 |
| RPL13P12 | 2.412 | 0.027 | 0.019 |
| RP11-235E17.2 | 2.337 | 0.032 | 0.019 |
| C6orf163 | -2.419 | 0.027 | 0.019 |
| RPS3AP26 | 2.404 | 0.028 | 0.019 |
| ZCCHC7 | 2.384 | 0.029 | 0.019 |
| RP1-257A7.4 | 2.312 | 0.034 | 0.019 |
| RBM33 | -2.400 | 0.028 | 0.019 |
| DUX4L51 | -2.395 | 0.028 | 0.019 |
| RANBP6 | -2.323 | 0.033 | 0.019 |
| POM121B | -2.320 | 0.033 | 0.019 |
| TRPV1 | 2.337 | 0.032 | 0.019 |
| TECPR2 | -2.321 | 0.033 | 0.019 |
| TNFSF12-TNFSF13 | 2.317 | 0.033 | 0.019 |

| | | | |
|---|---|---|---|
| RNU6-244P | -2.321 | 0.033 | 0.019 |
| TBC1D7 | 2.312 | 0.034 | 0.019 |
| RP11-110I1.13 | -2.356 | 0.031 | 0.019 |
| RGS7 | 2.349 | 0.031 | 0.019 |
| RHOBTB2 | -2.334 | 0.032 | 0.019 |
| RABGGTB | 2.258 | 0.037 | 0.019 |
| TREX2 | -2.333 | 0.032 | 0.019 |
| RP1-292L20.3 | -2.330 | 0.032 | 0.019 |
| TBC1D3C | 2.323 | 0.033 | 0.019 |
| AL022326.1 | 2.404 | 0.028 | 0.019 |
| UBR3 | -2.406 | 0.028 | 0.019 |
| SHPK | 2.337 | 0.032 | 0.020 |
| SNORD94 | 2.314 | 0.033 | 0.020 |
| RPL21P75 | 2.309 | 0.034 | 0.020 |
| AC007750.5 | 2.219 | 0.040 | 0.020 |
| DCP2 | -2.360 | 0.030 | 0.020 |
| LINC00667 | -2.331 | 0.032 | 0.020 |
| CRAT | -2.324 | 0.033 | 0.020 |
| MAST2 | -2.239 | 0.039 | 0.020 |
| MIF-AS1 | -2.382 | 0.029 | 0.020 |
| GLI4 | -2.346 | 0.031 | 0.020 |
| RPS7 | 2.291 | 0.035 | 0.020 |
| RP11-686D22.5 | 2.367 | 0.030 | 0.020 |
| TRIM38 | -2.356 | 0.031 | 0.020 |
| GNL3L | -2.313 | 0.033 | 0.020 |
| RAG1 | -2.340 | 0.032 | 0.020 |
| TNFSF13 | 2.317 | 0.033 | 0.020 |
| TUBD1 | -2.308 | 0.034 | 0.020 |
| TNFSF12 | 2.317 | 0.033 | 0.020 |
| RP13-228J13.5 | 2.358 | 0.031 | 0.020 |
| UBR1 | -2.346 | 0.031 | 0.020 |
| EVC | -2.331 | 0.032 | 0.021 |
| LNPEP | -2.269 | 0.037 | 0.021 |
| RP13-582O9.6 | -2.346 | 0.031 | 0.021 |
| GLYATL1P4 | -2.311 | 0.034 | 0.021 |
| ATP5D | 2.264 | 0.037 | 0.021 |
| CEP170B | -2.288 | 0.035 | 0.021 |
| RP11-407N17.5 | -2.284 | 0.035 | 0.021 |
| TAOK2 | -2.264 | 0.037 | 0.021 |
| XIAP | -2.343 | 0.032 | 0.021 |
| ATP5E | 2.329 | 0.032 | 0.021 |
| CEP170 | -2.324 | 0.033 | 0.021 |
| SRRM2 | -2.285 | 0.035 | 0.021 |
| RNU1-3 | 2.118 | 0.049 | 0.021 |
| TSHZ1 | -2.385 | 0.029 | 0.021 |
| ZFP41 | -2.346 | 0.031 | 0.021 |
| LINC01545 | -2.316 | 0.033 | 0.021 |
| TBC1D3L | 2.323 | 0.033 | 0.021 |
| bP-21201H5.1 | 2.264 | 0.037 | 0.021 |

| | | | |
|---|---|---|---|
| LINC00271 | 2.332 | 0.032 | 0.021 |
| ROMO1 | 2.361 | 0.030 | 0.021 |
| GLYATL1 | -2.311 | 0.034 | 0.021 |
| EIF2S3L | 2.326 | 0.033 | 0.021 |
| TRAF6 | -2.306 | 0.034 | 0.022 |
| RP11-1148L6.9 | 2.295 | 0.035 | 0.022 |
| RP5-1187M17.10 | 2.164 | 0.045 | 0.022 |
| POM121 | -2.320 | 0.033 | 0.022 |
| SLC16A13 | 2.280 | 0.036 | 0.022 |
| SPRY3 | -2.264 | 0.037 | 0.022 |
| SNRPA | 2.258 | 0.037 | 0.022 |
| IQCC | -2.292 | 0.035 | 0.022 |
| HMGB1P31 | 2.237 | 0.039 | 0.022 |
| TMCO6 | 2.324 | 0.033 | 0.022 |
| IL17RA | -2.322 | 0.033 | 0.022 |
| RP11-815J21.4 | -2.304 | 0.034 | 0.022 |
| RAB11B | -2.313 | 0.033 | 0.022 |
| RP11-171I2.2 | -2.272 | 0.036 | 0.022 |
| C16orf13 | 2.243 | 0.038 | 0.022 |
| IGF2R | -2.242 | 0.039 | 0.022 |
| NENF | 2.232 | 0.039 | 0.022 |
| RP11-288H12.3 | -2.242 | 0.039 | 0.022 |
| KB-1458E12.1 | 2.213 | 0.041 | 0.022 |
| VN1R82P | -2.295 | 0.035 | 0.023 |
| SIAE | -2.333 | 0.032 | 0.023 |
| RPE | 2.163 | 0.045 | 0.023 |
| RPL21P28 | 2.213 | 0.041 | 0.023 |
| RAB32 | 2.281 | 0.036 | 0.023 |
| SLC7A5P1 | 2.277 | 0.036 | 0.023 |
| SLC26A1 | -2.263 | 0.037 | 0.023 |
| PARGP1 | -2.255 | 0.038 | 0.023 |
| UTRN | -2.197 | 0.042 | 0.023 |
| KANSL2 | 2.262 | 0.037 | 0.023 |
| PRELID3B | 2.329 | 0.032 | 0.023 |
| RPL3P2 | 2.300 | 0.034 | 0.023 |
| SNORD45A | 2.258 | 0.037 | 0.023 |
| SLC7A2 | -2.220 | 0.040 | 0.023 |
| RP11-893F2.5 | -2.274 | 0.036 | 0.023 |
| TRIM26 | -2.262 | 0.037 | 0.023 |
| NDST1 | -2.255 | 0.038 | 0.023 |
| MTHFD2 | 2.281 | 0.036 | 0.024 |
| ZNF782 | -2.280 | 0.036 | 0.024 |
| SPTLC2 | -2.216 | 0.041 | 0.024 |
| RP11-819M15.2 | 2.144 | 0.047 | 0.024 |
| CYGB | -2.210 | 0.041 | 0.024 |
| ACADM | 2.258 | 0.037 | 0.024 |
| ZNF320 | -2.232 | 0.039 | 0.024 |
| RP11-1391J7.1 | -2.211 | 0.041 | 0.024 |
| PDPK2P | -2.282 | 0.036 | 0.024 |

| | | | |
|---|---|---|---|
| SNORD45C | 2.258 | 0.037 | 0.024 |
| CTC-308K20.1 | 2.211 | 0.041 | 0.024 |
| ZNF256 | -2.299 | 0.034 | 0.024 |
| WDPCP | -2.238 | 0.039 | 0.024 |
| HSD17B12 | -2.238 | 0.039 | 0.024 |
| AC092117.2 | -2.285 | 0.035 | 0.024 |
| RPSAP19 | 2.211 | 0.041 | 0.024 |
| SUPT6H | -2.254 | 0.038 | 0.024 |
| RP11-876N24.3 | -2.268 | 0.037 | 0.024 |
| MIR1231 | -2.210 | 0.041 | 0.024 |
| IFFO1 | -2.247 | 0.038 | 0.025 |
| PSMD5-AS1 | 2.280 | 0.036 | 0.025 |
| MIR670HG | -2.238 | 0.039 | 0.025 |
| AP001189.4 | -2.194 | 0.042 | 0.025 |
| GCC2 | -2.279 | 0.036 | 0.025 |
| LL0XNC01-36H8.1 | -2.266 | 0.037 | 0.025 |
| ZNF816-ZNF321P | -2.232 | 0.039 | 0.025 |
| ZNF486 | -2.199 | 0.042 | 0.025 |
| DPYSL4 | 2.251 | 0.038 | 0.025 |
| CTD-2331H12.8 | -2.232 | 0.039 | 0.025 |
| FAUP1 | 2.217 | 0.041 | 0.025 |
| NCOR2 | -2.245 | 0.038 | 0.025 |
| NMNAT3 | 2.231 | 0.039 | 0.025 |
| TAF7 | 2.191 | 0.043 | 0.025 |
| SAP30L-AS1 | -2.178 | 0.044 | 0.025 |
| CTC-350I8.1 | -2.155 | 0.046 | 0.025 |
| RP4-761J14.8 | -2.282 | 0.036 | 0.025 |
| FGFR4 | 2.246 | 0.038 | 0.025 |
| NUDT6 | -2.213 | 0.041 | 0.025 |
| MGST3 | 2.185 | 0.043 | 0.025 |
| GLUD1P3 | -2.249 | 0.038 | 0.025 |
| PCID2 | 2.197 | 0.042 | 0.025 |
| KIAA1683 | 2.306 | 0.034 | 0.025 |
| AC004556.1 | 2.255 | 0.038 | 0.025 |
| ZNF702P | -2.232 | 0.039 | 0.025 |
| ASB14 | 2.169 | 0.045 | 0.026 |
| ATRX | -2.255 | 0.038 | 0.026 |
| INPP4A | -2.252 | 0.038 | 0.026 |
| RPSAP47 | 2.250 | 0.038 | 0.026 |
| THOC6 | 2.165 | 0.045 | 0.026 |
| SNORD36B | 2.189 | 0.043 | 0.026 |
| C17orf107 | 2.239 | 0.039 | 0.026 |
| PPARA | 2.228 | 0.040 | 0.026 |
| ADAM21 | 2.186 | 0.043 | 0.026 |
| SNORD45B | 2.258 | 0.037 | 0.026 |
| ZNF321P | -2.232 | 0.039 | 0.026 |
| SNORD36C | 2.189 | 0.043 | 0.026 |
| DYNLT1 | 2.206 | 0.041 | 0.026 |
| CEP89 | -2.204 | 0.042 | 0.026 |

| | | | |
|---|---|---|---|
| SPRYD3 | -2.184 | 0.043 | 0.026 |
| TMEM138 | -2.221 | 0.040 | 0.027 |
| RP5-832C2.5 | -2.213 | 0.041 | 0.027 |
| NRG1 | -2.191 | 0.043 | 0.027 |
| CENPO | -2.216 | 0.041 | 0.027 |
| RWDD2B | 2.215 | 0.041 | 0.027 |
| RP1-139D8.6 | 2.207 | 0.041 | 0.027 |
| RPL4P6 | 2.196 | 0.042 | 0.027 |
| SNORA3A | 2.173 | 0.044 | 0.027 |
| SCN9A | -2.166 | 0.045 | 0.027 |
| ACAP2 | -2.194 | 0.042 | 0.027 |
| CTD-2620I22.1 | -2.232 | 0.039 | 0.027 |
| BNIP3L | -2.197 | 0.042 | 0.027 |
| HIST2H3PS2 | -2.191 | 0.043 | 0.027 |
| APP | -2.232 | 0.039 | 0.027 |
| RP11-666A8.8 | -2.210 | 0.041 | 0.027 |
| GUCA1A | 2.207 | 0.041 | 0.027 |
| CPSF3 | 2.183 | 0.043 | 0.027 |
| MAP7D1 | -2.203 | 0.042 | 0.027 |
| SNORA3B | 2.173 | 0.044 | 0.027 |
| AC090498.1 | 2.153 | 0.046 | 0.027 |
| RP5-998N21.10 | -2.191 | 0.043 | 0.028 |
| GSTM2 | 2.173 | 0.044 | 0.028 |
| ACVR2B-AS1 | 2.159 | 0.045 | 0.028 |
| AC091053.1 | 2.173 | 0.044 | 0.028 |
| WDSUB1 | 2.161 | 0.045 | 0.028 |
| HIST2H2BB | -2.191 | 0.043 | 0.028 |
| TPRG1L | -2.141 | 0.047 | 0.028 |
| AL449212.1 | -2.160 | 0.045 | 0.028 |
| ZFYVE26 | -2.159 | 0.045 | 0.028 |
| RP11-234A1.1 | 2.212 | 0.041 | 0.028 |
| IAH1 | 2.183 | 0.043 | 0.028 |
| MAGI3 | -2.183 | 0.043 | 0.028 |
| AL121895.1 | -2.178 | 0.044 | 0.028 |
| GSTM3 | 2.171 | 0.044 | 0.028 |
| SLC7A9 | -2.204 | 0.042 | 0.028 |
| GNG12-AS1 | -2.201 | 0.042 | 0.028 |
| CCDC28A | 2.191 | 0.043 | 0.028 |
| SNORD24 | 2.189 | 0.043 | 0.028 |
| CEBPZOS | -2.172 | 0.044 | 0.028 |
| MZT2B | 2.122 | 0.049 | 0.028 |
| KDM6A | -2.221 | 0.040 | 0.029 |
| TRIM61 | -2.160 | 0.045 | 0.029 |
| STK3 | 2.213 | 0.041 | 0.029 |
| RP11-553K23.2 | 2.231 | 0.039 | 0.029 |
| KHDRBS3 | 2.196 | 0.042 | 0.029 |
| AAR2 | -2.178 | 0.044 | 0.029 |
| SNORA7B | -2.160 | 0.045 | 0.029 |
| ZNF816 | -2.232 | 0.039 | 0.029 |

| | | | |
|---|---|---|---|
| ZSWIM3 | -2.209 | 0.041 | 0.029 |
| SNORD36A | 2.189 | 0.043 | 0.029 |
| RP11-34P13.13 | -2.161 | 0.045 | 0.029 |
| RP11-863K10.7 | -2.150 | 0.046 | 0.029 |
| RPL7A | 2.189 | 0.043 | 0.030 |
| ZNF595 | -2.178 | 0.044 | 0.030 |
| SUGP2 | -2.162 | 0.045 | 0.030 |
| GUSBP3 | -2.138 | 0.047 | 0.030 |
| LTBR | -2.183 | 0.043 | 0.030 |
| ZFAND1 | 2.133 | 0.048 | 0.030 |
| LA16c-17H1.3 | 2.183 | 0.043 | 0.030 |
| CRMP1 | -2.183 | 0.043 | 0.030 |
| SH2D2A | 2.168 | 0.045 | 0.030 |
| TGIF1 | 2.167 | 0.045 | 0.030 |
| RPL27A | 2.173 | 0.044 | 0.030 |
| RP11-366M4.11 | -2.160 | 0.045 | 0.030 |
| PPP2R5A | 2.122 | 0.049 | 0.030 |
| LAMP2 | -2.165 | 0.045 | 0.030 |
| RP11-296E7.1 | 2.186 | 0.043 | 0.030 |
| CASD1 | 2.178 | 0.044 | 0.030 |
| ZCCHC9 | 2.168 | 0.045 | 0.030 |
| SNORA59A | -2.199 | 0.042 | 0.030 |
| UBE2D1 | -2.196 | 0.042 | 0.030 |
| CFAP44 | -2.173 | 0.044 | 0.030 |
| IRAK1BP1 | 2.163 | 0.045 | 0.030 |
| TRIB1 | 2.210 | 0.041 | 0.031 |
| MAP2K7 | -2.178 | 0.044 | 0.031 |
| RP1-102E24.8 | -2.183 | 0.043 | 0.031 |
| GSTM5 | 2.173 | 0.044 | 0.031 |
| OSTM1 | -2.160 | 0.045 | 0.031 |
| MAST1 | 2.142 | 0.047 | 0.031 |
| EPM2A | 2.140 | 0.047 | 0.031 |
| GSTM1 | 2.173 | 0.044 | 0.031 |
| PTPN3 | 2.161 | 0.045 | 0.031 |
| GSTM4 | 2.173 | 0.044 | 0.031 |
| B9D1 | 2.140 | 0.047 | 0.031 |
| ZNF330 | 2.141 | 0.047 | 0.031 |
| RP13-270P17.1 | 2.149 | 0.046 | 0.031 |
| AC091053.2 | 2.173 | 0.044 | 0.032 |
| RP11-423P10.2 | -2.172 | 0.044 | 0.032 |
| CTD-3247H4.2 | -2.159 | 0.045 | 0.032 |
| H3F3BP1 | 2.153 | 0.046 | 0.032 |
| PPP2R5E | -2.143 | 0.047 | 0.032 |
| RPPH1 | 2.112 | 0.050 | 0.032 |
| RP4-735C1.4 | 2.173 | 0.044 | 0.032 |
| MAML2 | -2.145 | 0.047 | 0.032 |
| RP11-33N14.3 | 2.138 | 0.047 | 0.032 |
| RP1-111B22.3 | -2.160 | 0.045 | 0.032 |
| RP5-1142A6.2 | -2.137 | 0.047 | 0.032 |

| | | | |
|---|---|---|---|
| RP11-50D9.1 | 2.129 | 0.048 | 0.033 |
| KIF3B | -2.123 | 0.049 | 0.033 |
| RP11-796G6.1 | 2.122 | 0.049 | 0.033 |
| BMPER | -2.119 | 0.049 | 0.033 |
| RP11-566K11.5 | -2.123 | 0.049 | 0.033 |
| CPPED1 | -2.127 | 0.048 | 0.033 |
| ZSCAN23 | -2.154 | 0.046 | 0.034 |
| ZFHX4 | -2.139 | 0.047 | 0.034 |
| MIR656 | -2.134 | 0.048 | 0.034 |
| BTF3 | 2.128 | 0.048 | 0.034 |
| CEP250 | -2.115 | 0.050 | 0.034 |
| TIGD6 | -2.113 | 0.050 | 0.035 |
| RP11-968O1.5 | -2.121 | 0.049 | 0.035 |
| CD44 | -2.122 | 0.049 | 0.035 |

**Table S6. Genes that were significantly correlated with the right olfactory bulb (OB_R) volume in FEP patients and also identified by a genome-wide association study (GWAS) as risk genes for brain disorders.**

Abbreviations: SZ, schizophrenia; BP, bipolar disorder; OR, odd ratio.

| | Correlation with OB_R | | | GWAS | | |
|---|---|---|---|---|---|---|
| **Gene** | **t-value** | **p-value** | **p-value (permutation)** | **Trait** | **-log10(p-value)** | **OR** |
| AS3MT | -3.833 | 0.009 | 2.00E-04 | SZ | 19.398 | 1.091 |
| BORCS7 | -3.833 | 0.009 | 3.00E-04 | SZ | 16.000 | 1.087 |
| BORCS7-ASMT | -3.833 | 0.009 | 2.00E-04 | SZ | 16.000 | 1.087 |
| C3orf70 | -2.495 | 0.047 | 0.008 | unipolar depression | 10.301 | 1.160 |
| C4B | -3.143 | 0.020 | 0.002 | SZ | 17.222 | 1.163 |
| CERS6 | -2.942 | 0.026 | 0.007 | Parkinson | 15.523 | 1.209 |
| CHADL | -2.470 | 0.048 | 0.007 | SZ | 10.699 | 1.087 |
| CNNM2 | -3.210 | 0.018 | 0.003 | SZ | 19.398 | 1.091 |
| CTNNA1 | -2.586 | 0.041 | 0.004 | SZ | 8.301 | 1.063 |
| CTSB | 2.488 | 0.047 | 0.006 | Parkinson | 11.222 | 1.090 |
| ERBB2 | -2.571 | 0.042 | 0.004 | BP | 8.301 | 1.130 |
| FADS1 | 2.740 | 0.034 | 0.001 | BP | 10.000 | 1.126 |
| FAM109B | -5.030 | 0.002 | 0.000 | SZ | 8.398 | 1.064 |
| FAM171A1 | -3.158 | 0.020 | 0.005 | Parkinson | 8.000 | 1.075 |
| FGFR1 | 2.999 | 0.024 | 0.008 | SZ | 8.398 | 1.075 |
| FLOT1 | -4.177 | 0.006 | 0.000 | SZ | 22.000 | 1.240 |
| HACE1 | -2.772 | 0.032 | 0.002 | SZ | 8.097 | 1.056 |
| HFE | -2.856 | 0.029 | 0.002 | SZ | 23.097 | 1.210 |
| HHAT | -3.636 | 0.011 | 0.000 | SZ | 8.222 | 2.630 |
| HLA-DOB | 2.747 | 0.033 | 0.002 | SZ | 17.222 | 1.163 |
| HMGN4 | 4.844 | 0.003 | 0.000 | SZ | 23.097 | 1.210 |
| IER3 | -4.177 | 0.006 | 1.00E-04 | SZ | 22.000 | 1.240 |
| INA | -3.844 | 0.009 | 2.00E-04 | SZ | 18.222 | 1.104 |
| LINC00933 | -2.550 | 0.043 | 0.007 | SZ | 10.222 | 1.075 |
| LMAN2L | -2.716 | 0.035 | 0.004 | BP | 9.523 | 1.120 |
| LRRK2 | -5.122 | 0.002 | 0.000 | Parkinson | 39.000 | 3.115 |
| MAU2 | -2.507 | 0.046 | 0.006 | SZ | 9.398 | 1.071 |
| MED8 | 2.462 | 0.049 | 0.005 | SZ | 12.000 | 1.072 |
| MIR6832 | 2.698 | 0.036 | 0.003 | SZ | 19.097 | 1.163 |
| MMP16 | -2.714 | 0.035 | 0.002 | SZ | 8.000 | 1.065 |
| NAB2 | 2.949 | 0.026 | 0.002 | SZ | 11.699 | 1.182 |
| NDUFA6-AS1 | -2.457 | 0.049 | 0.005 | SZ | 8.398 | 1.064 |
| NDUFAF2 | 2.516 | 0.046 | 0.007 | Parkinson | 8.000 | 1.163 |
| NME5 | -2.750 | 0.033 | 0.001 | SZ | 8.000 | 1.064 |
| PHF7 | -3.398 | 0.015 | 0.000 | SZ | 11.523 | 1.075 |
| PLCL1 | -2.907 | 0.027 | 0.005 | SZ | 10.699 | 1.076 |
| PPP2R3A | 2.450 | 0.050 | 0.003 | SZ | 10.155 | 1.072 |
| RNF5 | -2.591 | 0.041 | 0.001 | SZ | 17.222 | 1.163 |
| SDAD1P1 | -2.514 | 0.046 | 0.018 | SZ | 10.699 | 1.078 |

| | | | | | | |
|---|---|---|---|---|---|---|
| SERPING1 | -2.685 | 0.036 | 0.004 | SZ | 8.699 | 1.068 |
| SMG6 | -2.950 | 0.026 | 0.003 | SZ | 9.523 | 1.071 |
| SNORD69 | 3.957 | 0.007 | 1.00E-04 | SZ | 11.523 | 1.075 |
| STAT6 | 6.712 | 0.001 | 0.000 | SZ | 11.699 | 1.182 |
| TAP2 | 2.747 | 0.033 | 0.001 | SZ | 17.222 | 1.163 |
| TLR9 | 2.962 | 0.025 | 0.001 | SZ | 11.523 | 1.075 |
| TMEM175 | -2.700 | 0.036 | 0.001 | Parkinson | 50.000 | 1.230 |
| TWF2 | 2.962 | 0.025 | 0.001 | SZ | 11.523 | 1.075 |
| TYW5 | -2.557 | 0.043 | 0.006 | SZ | 15.699 | 1.096 |
| ZNF322 | -3.385 | 0.015 | 0.000 | SZ | 23.398 | 1.200 |

**Table S7. Significant pathways overrepresented in genes associated with the right olfactory bulb volume in first episode psychosis patients.**
Meaning of columns: size, the number of genes in the child pathway; NES, normalized enrichment score; FDR, false discovery rate

| Child pathway | Parent network | size | NES | FDR |
|---|---|---|---|---|
| ABC-FAMILY PROTEINS MEDIATED TRANSPORT | Transport_of_small_molecules | 72 | 2.213 | 0.000 |
| ACTIVATION OF NF-KAPPAB IN B CELLS | Immune_System | 62 | 2.336 | 0.000 |
| APC/C:CDH1 MEDIATED DEGRADATION OF CDC20 AND OTHER APC/C:CDH1 TARGETED PROTEINS IN LATE MITOSIS/EARLY G1 | Cell_Cycle | 69 | 2.220 | 0.000 |
| ASYMMETRIC LOCALIZATION OF PCP PROTEINS | Signal_Transduction | 59 | 2.330 | 0.000 |
| AUF1 (HNRNP D0) BINDS AND DESTABILIZES MRNA | Metabolism_of_RNA | 52 | 2.563 | 0.000 |
| AUTODEGRADATION OF CDH1 BY CDH1:APC/C | Cell_Cycle | 62 | 2.226 | 0.000 |
| CDK-MEDIATED PHOSPHORYLATION AND REMOVAL OF CDC6 | DNA_Replication | 69 | 2.154 | 0.001 |
| CDT1 ASSOCIATION WITH THE CDC6:ORC:ORIGIN COMPLEX | DNA_Replication | 56 | 2.384 | 0.000 |
| CHOLESTEROL BIOSYNTHESIS | Metabolism | 20 | 1.975 | 0.005 |
| CITRIC ACID CYCLE (TCA CYCLE) | Metabolism | 22 | 2.206 | 0.000 |
| CLEC7A (DECTIN-1) SIGNALING | Immune_System | 72 | 2.055 | 0.002 |
| COMPLEX I BIOGENESIS | Metabolism | 55 | 2.083 | 0.001 |
| CROSS-PRESENTATION OF SOLUBLE EXOGENOUS ANTIGENS (ENDOSOMES) | Immune_System | 43 | 2.479 | 0.000 |
| DECTIN-1 MEDIATED NONCANONICAL NF-KB SIGNALING | Immune_System | 57 | 2.284 | 0.000 |
| DEFECTIVE CFTR CAUSES CYSTIC FIBROSIS | Disease | 57 | 2.129 | 0.001 |
| DEGRADATION OF AXIN | Signal_Transduction | 52 | 2.162 | 0.001 |
| DEGRADATION OF BETA-CATENIN BY THE DESTRUCTION COMPLEX | Signal_Transduction | 65 | 1.994 | 0.004 |
| DEGRADATION OF DVL | Signal_Transduction | 54 | 2.311 | 0.000 |
| DEGRADATION OF GLI1 BY THE PROTEASOME | Signal_Transduction | 56 | 2.210 | 0.000 |
| DEGRADATION OF GLI2 BY THE PROTEASOME | Signal_Transduction | 56 | 2.133 | 0.001 |
| DEREGULATED CDK5 TRIGGERS MULTIPLE NEURODEGENERATIVE PATHWAYS IN ALZHEIMER'S DISEASE MODELS | Disease | 21 | 1.767 | 0.034 |

| | | | | |
|---|---|---|---|---|
| DOWNSTREAM TCR SIGNALING | Immune_System | 75 | 2.025 | 0.003 |
| EPHB-MEDIATED FORWARD SIGNALING | Developmental_Biology | 39 | 1.718 | 0.046 |
| ER-PHAGOSOME PATHWAY | Immune_System | 78 | 1.952 | 0.007 |
| EUKARYOTIC TRANSLATION TERMINATION | Metabolism_of_proteins | 90 | 2.708 | 0.000 |
| FATTY ACYL-COA BIOSYNTHESIS | Metabolism | 11 | 1.791 | 0.028 |
| FBXL7 DOWN-REGULATES AURKA DURING MITOTIC ENTRY AND IN EARLY MITOSIS | Cell_Cycle | 52 | 2.325 | 0.000 |
| FCERI MEDIATED NF-KB ACTIVATION | Immune_System | 74 | 2.068 | 0.002 |
| FOLDING OF ACTIN BY CCT/TRIC | Metabolism_of_proteins | 10 | 1.725 | 0.044 |
| FORMATION OF A POOL OF FREE 40S SUBUNITS | Metabolism_of_proteins | 98 | 2.767 | 0.000 |
| FORMATION OF TC-NER PRE-INCISION COMPLEX | DNA_Repair | 53 | 1.752 | 0.037 |
| FORMATION OF THE TERNARY COMPLEX; AND SUBSEQUENTLY; THE 43S COMPLEX | Metabolism_of_proteins | 50 | 2.418 | 0.000 |
| FORMATION OF TUBULIN FOLDING INTERMEDIATES BY CCT/TRIC | Metabolism_of_proteins | 20 | 1.910 | 0.009 |
| G2/M CHECKPOINTS | Cell_Cycle | 48 | 2.519 | 0.000 |
| GENERIC TRANSCRIPTION PATHWAY | Gene_expression_(Transcription) | 322 | -2.534 | 0.000 |
| GLI3 IS PROCESSED TO GLI3R BY THE PROTEASOME | Signal_Transduction | 56 | 2.167 | 0.001 |
| GLUTATHIONE CONJUGATION | Metabolism | 19 | 1.865 | 0.014 |
| GTP HYDROLYSIS AND JOINING OF THE 60S RIBOSOMAL SUBUNIT | Metabolism_of_proteins | 109 | 2.733 | 0.000 |
| HEDGEHOG 'ON' STATE | Signal_Transduction | 67 | 2.151 | 0.001 |
| HEDGEHOG LIGAND BIOGENESIS | Signal_Transduction | 54 | 2.160 | 0.001 |
| HH MUTANTS THAT DON'T UNDERGO AUTOCATALYTIC PROCESSING ARE DEGRADED BY ERAD | Disease | 52 | 2.354 | 0.000 |
| INTERLEUKIN-1 SIGNALING | Immune_System | 78 | 1.910 | 0.009 |
| L13A-MEDIATED TRANSLATIONAL SILENCING OF CERULOPLASMIN EXPRESSION | Metabolism_of_proteins | 108 | 2.709 | 0.000 |
| MAJOR PATHWAY OF RRNA PROCESSING IN THE NUCLEOLUS AND CYTOSOL | Metabolism_of_RNA | 176 | 2.832 | 0.000 |
| MAPK6/MAPK4 SIGNALING | Signal_Transduction | 83 | 1.883 | 0.012 |
| MITOCHONDRIAL PROTEIN IMPORT | Metabolism_of_proteins | 60 | 1.995 | 0.004 |
| MITOCHONDRIAL TRANSLATION ELONGATION | Metabolism_of_proteins | 88 | 2.626 | 0.000 |
| MITOCHONDRIAL TRANSLATION INITIATION | Metabolism_of_proteins | 88 | 2.571 | 0.000 |

| | | | | |
|---|---|---|---|---|
| MITOCHONDRIAL TRANSLATION TERMINATION | Metabolism_of_proteins | 88 | 2.675 | 0.000 |
| MRNA SPLICING - MAJOR PATHWAY | Metabolism_of_RNA | 178 | 2.367 | 0.000 |
| MRNA SPLICING - MINOR PATHWAY | Metabolism_of_RNA | 52 | 2.174 | 0.000 |
| NIK-->NONCANONICAL NF-KB SIGNALING | Immune_System | 56 | 2.281 | 0.000 |
| NONSENSE MEDIATED DECAY (NMD) ENHANCED BY THE EXON JUNCTION COMPLEX (EJC) | Metabolism_of_RNA | 112 | 2.445 | 0.000 |
| NONSENSE MEDIATED DECAY (NMD) INDEPENDENT OF THE EXON JUNCTION COMPLEX (EJC) | Metabolism_of_RNA | 92 | 2.697 | 0.000 |
| ORC1 REMOVAL FROM CHROMATIN | DNA_Replication | 68 | 2.411 | 0.000 |
| OXYGEN-DEPENDENT PROLINE HYDROXYLATION OF HYPOXIA-INDUCIBLE FACTOR ALPHA | Cellular_responses_to_external_stimuli | 60 | 2.130 | 0.001 |
| PENTOSE PHOSPHATE PATHWAY (HEXOSE MONOPHOSPHATE SHUNT) | Metabolism | 11 | 1.851 | 0.016 |
| PEPTIDE CHAIN ELONGATION | Metabolism_of_proteins | 86 | 2.745 | 0.000 |
| POST-CHAPERONIN TUBULIN FOLDING PATHWAY | Metabolism_of_proteins | 17 | 1.753 | 0.037 |
| PREFOLDIN MEDIATED TRANSFER OF SUBSTRATE TO CCT/TRIC | Metabolism_of_proteins | 25 | 2.199 | 0.000 |
| PROCESSING OF CAPPED INTRON-CONTAINING PRE-MRNA | Metabolism_of_RNA | 37 | 1.966 | 0.006 |
| PURINE RIBONUCLEOSIDE MONOPHOSPHATE BIOSYNTHESIS | Metabolism | 12 | 1.949 | 0.007 |
| RECYCLING PATHWAY OF L1 | Developmental_Biology | 38 | 1.925 | 0.008 |
| REGULATION OF ACTIN DYNAMICS FOR PHAGOCYTIC CUP FORMATION | Immune_System | 50 | 1.789 | 0.028 |
| REGULATION OF ACTIVATED PAK-2P34 BY PROTEASOME MEDIATED DEGRADATION | Programmed_Cell_Death | 47 | 2.478 | 0.000 |
| REGULATION OF EXPRESSION OF SLITS AND ROBOS | Developmental_Biology | 156 | 2.734 | 0.000 |
| REGULATION OF ORNITHINE DECARBOXYLASE (ODC) | Metabolism | 48 | 2.516 | 0.000 |
| REGULATION OF PTEN STABILITY AND ACTIVITY | Signal_Transduction | 64 | 2.061 | 0.002 |
| REGULATION OF RAS BY GAPS | Signal_Transduction | 63 | 2.048 | 0.002 |
| REGULATION OF RUNX2 EXPRESSION AND ACTIVITY | Gene_expression_(Transcription) | 66 | 2.189 | 0.000 |
| REGULATION OF RUNX3 EXPRESSION AND ACTIVITY | Gene_expression_(Transcription) | 54 | 2.412 | 0.000 |
| RESPIRATORY ELECTRON TRANSPORT | Metabolism | 88 | 2.675 | 0.000 |

| | | | | |
|---|---|---|---|---|
| RIBOSOMAL SCANNING AND START CODON RECOGNITION | Metabolism_of_proteins | 57 | 2.307 | 0.000 |
| RNA POLYMERASE I PROMOTER ESCAPE | Gene_expression_(Transcription) | 30 | 1.717 | 0.046 |
| RRNA MODIFICATION IN THE NUCLEUS AND CYTOSOL | Metabolism_of_RNA | 58 | 2.791 | 0.000 |
| RUNX1 REGULATES TRANSCRIPTION OF GENES INVOLVED IN DIFFERENTIATION OF HSCS | Gene_expression_(Transcription) | 95 | 2.189 | 0.000 |
| SCF(SKP2)-MEDIATED DEGRADATION OF P27/P21 | Cell_Cycle | 58 | 2.161 | 0.001 |
| SELENOCYSTEINE SYNTHESIS | Metabolism | 90 | 2.562 | 0.000 |
| SIGNAL TRANSDUCTION BY L1 | Developmental_Biology | 18 | 1.751 | 0.036 |
| SIGNALING BY PDGF | Signal_Transduction | 23 | -1.994 | 0.042 |
| SIGNALING BY ROBO RECEPTORS | Developmental_Biology | 12 | 1.860 | 0.015 |
| SRP-DEPENDENT COTRANSLATIONAL PROTEIN TARGETING TO MEMBRANE | Metabolism_of_proteins | 109 | 2.643 | 0.000 |
| THE ROLE OF GTSE1 IN G2/M PROGRESSION AFTER G2 CHECKPOINT | Cell_Cycle | 68 | 2.540 | 0.000 |
| TNFR2 NON-CANONICAL NF-KB PATHWAY | Immune_System | 58 | 2.525 | 0.000 |
| UBIQUITIN-DEPENDENT DEGRADATION OF CYCLIN D1 | Cell_Cycle | 49 | 2.454 | 0.000 |
| UCH PROTEINASES | Metabolism_of_proteins | 93 | 1.962 | 0.006 |
| VIF-MEDIATED DEGRADATION OF APOBEC3G | Disease | 49 | 2.480 | 0.000 |
| VPU MEDIATED DEGRADATION OF CD4 | Disease | 49 | 2.446 | 0.000 |

**Table S8. Significant pathways overrepresented in genes associated with the right olfactory bulb volume in healthy controls.**

Meaning of columns: size, the number of genes in the child pathway; NES, normalized enrichment score; FDR, false discovery rate

| Child pathway | Parent network | size | NES | FDR |
|---|---|---|---|---|
| ANCHORING OF THE BASAL BODY TO THE PLASMA MEMBRANE | Organelle_biogenesis_and_maintenance | 96 | -1.934 | 0.045 |
| CYTOSOLIC TRNA AMINOACYLATION | Metabolism_of_proteins | 24 | 1.911 | 0.023 |
| EUKARYOTIC TRANSLATION TERMINATION | Metabolism_of_proteins | 90 | 3.313 | 0.000 |
| FORMATION OF A POOL OF FREE 40S SUBUNITS | Metabolism_of_proteins | 98 | 3.257 | 0.000 |
| FORMATION OF THE TERNARY COMPLEX; AND SUBSEQUENTLY; THE 43S COMPLEX | Metabolism_of_proteins | 50 | 2.721 | 0.000 |
| GENERIC TRANSCRIPTION PATHWAY | Gene_expression_(Transcription) | 322 | -2.141 | 0.006 |
| GTP HYDROLYSIS AND JOINING OF THE 60S RIBOSOMAL SUBUNIT | Metabolism_of_proteins | 109 | 3.231 | 0.000 |
| L13A-MEDIATED TRANSLATIONAL SILENCING OF CERULOPLASMIN EXPRESSION | Metabolism_of_proteins | 108 | 3.219 | 0.000 |
| MAJOR PATHWAY OF RRNA PROCESSING IN THE NUCLEOLUS AND CYTOSOL | Metabolism_of_RNA | 176 | 2.715 | 0.000 |
| MYOCLONIC EPILEPSY OF LAFORA | Disease | 10 | 1.925 | 0.021 |
| NONSENSE MEDIATED DECAY (NMD) ENHANCED BY THE EXON JUNCTION COMPLEX (EJC) | Metabolism_of_RNA | 112 | 2.922 | 0.000 |
| NONSENSE MEDIATED DECAY (NMD) INDEPENDENT OF THE EXON JUNCTION COMPLEX (EJC) | Metabolism_of_RNA | 92 | 3.317 | 0.000 |
| PEPTIDE CHAIN ELONGATION | Metabolism_of_proteins | 86 | 3.400 | 0.000 |
| REGULATION OF EXPRESSION OF SLITS AND ROBOS | Developmental_Biology | 156 | 2.787 | 0.000 |
| RIBOSOMAL SCANNING AND START CODON RECOGNITION | Metabolism_of_proteins | 57 | 2.631 | 0.000 |
| SELENOCYSTEINE SYNTHESIS | Metabolism | 90 | 3.356 | 0.000 |
| SRP-DEPENDENT COTRANSLATIONAL PROTEIN TARGETING TO MEMBRANE | Metabolism_of_proteins | 109 | 3.224 | 0.000 |

**Supplementary figure legends**

**Figure S1. The volcano plot for differential expression analysis between first episode psychosis patients and healthy controls.**
Black dots represent significant genes with false discovery rate smaller then 0.05, while gray dots represent genes that were below the significant cutoff.

**Figure S2. Gene set enrichment analysis (GSEA) results of genes associated with the right olfactory bulb (OB_R) volume.**
GSEA identified 88 and 17 significant pathways overrepresented in genes associated with the OB_R in FEP patients (A) and healthy controls (B), respectively. Significant pathways were further grouped based on the parent-child hierarchical structure of pathways provided by the Reactome Pathway Database. The y-axis has all the parent networks from Reactome Pathway Database and the x-axis showed the number of significant child pathways under the corresponding parent network.

**Figure S1.**

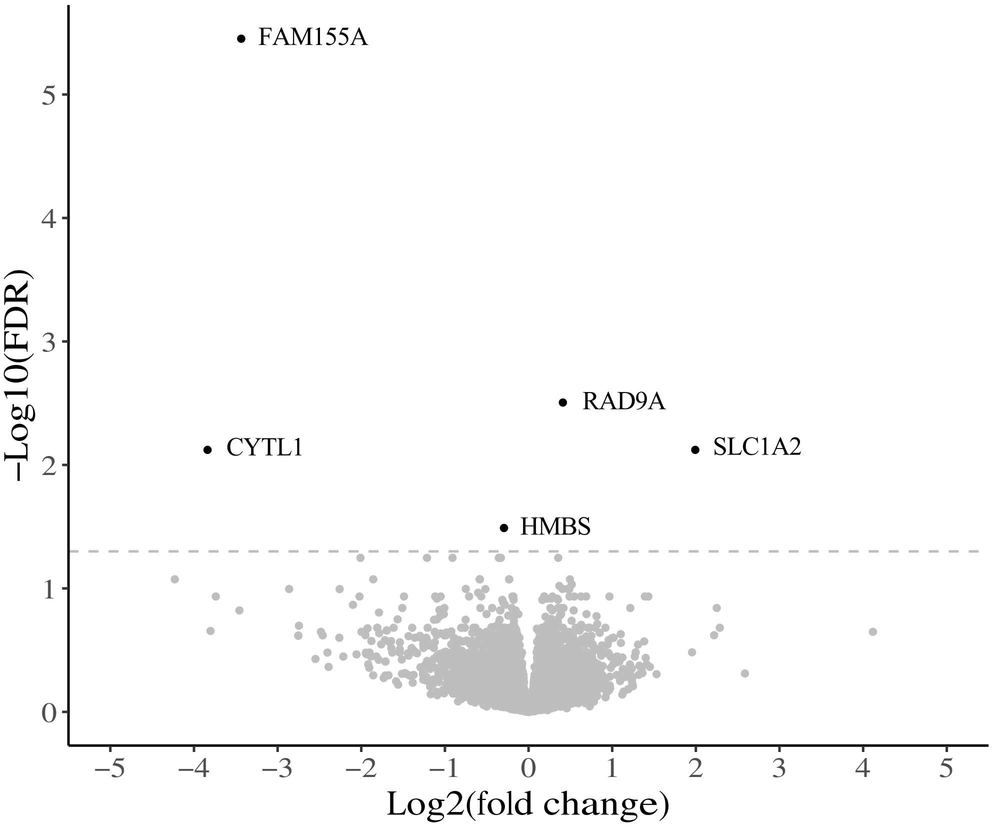
FAM155A
RAD9A
CYTL1
SLC1A2
HMBS
5
4
3
2
1
0
−Log10(FDR)
−5
−4
−3
−2
−1
0
1
2
3
4
5
Log2(fold change)

**Figure S2.**

A

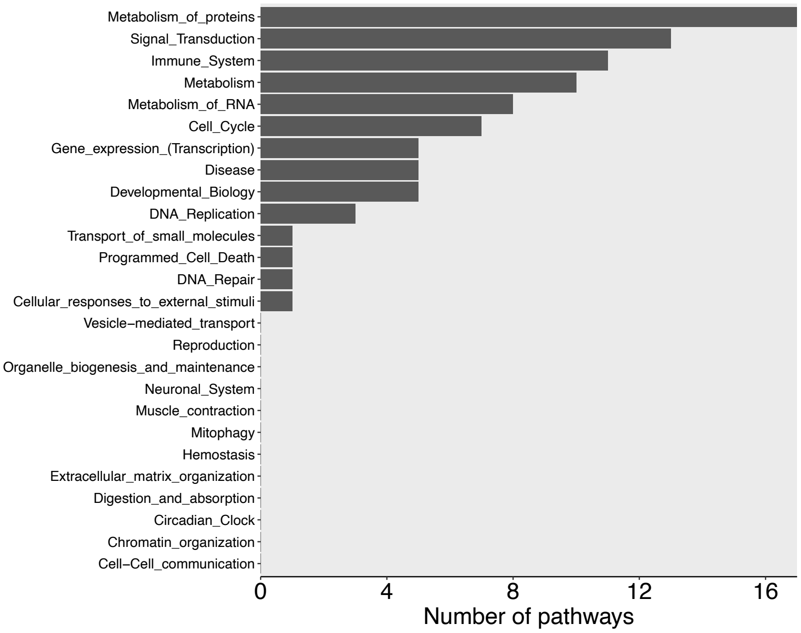
Metabolism_of_proteins
Signal_Transduction
Immune_System
Metabolism
Metabolism_of_RNA
Cell_Cycle
Gene_expression_(Transcription)
Disease
Developmental_Biology
DNA_Replication
Transport_of_small_molecules
Programmed_Cell_Death
DNA_Repair
Cellular_responses_to_external_stimuli
Vesicle−mediated_transport
Reproduction
Organelle_biogenesis_and_maintenance
Neuronal_System
Muscle_contraction
Mitophagy
Hemostasis
Extracellular_matrix_organization
Digestion_and_absorption
Circadian_Clock
Chromatin_organization
Cell−Cell_communication
0
4
8
12
16
Number of pathways

B

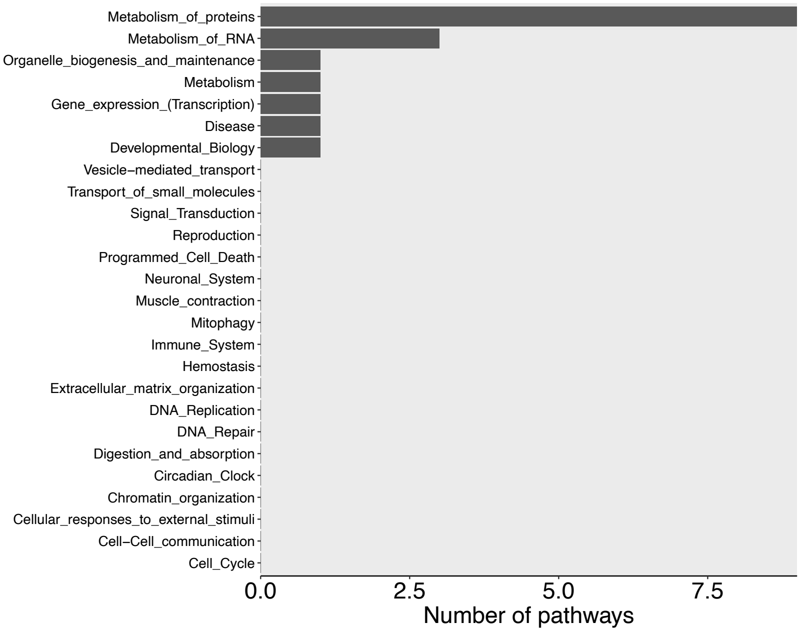
Metabolism_of_proteins
Metabolism_of_RNA
Organelle_biogenesis_and_maintenance
Metabolism
Gene_expression_(Transcription)
Disease
Developmental_Biology
Vesicle−mediated_transport
Transport_of_small_molecules
Signal_Transduction
Reproduction
Programmed_Cell_Death
Neuronal_System
Muscle_contraction
Mitophagy
Immune_System
Hemostasis
Extracellular_matrix_organization
DNA_Replication
DNA_Repair
Digestion_and_absorption
Circadian_Clock
Chromatin_organization
Cellular_responses_to_external_stimuli
Cell−Cell_communication
Cell_Cycle
0.0
2.5
5.0
7.5
Number of pathways
